# Suppression of upstream ORF translation is not a widespread mechanism of translational stimulation by yeast helicase Ded1

**DOI:** 10.64898/2026.04.10.717766

**Authors:** Rakesh Kumar, Gemma May, Neelam Dabas Sen, C. Joel McManus, Alan G. Hinnebusch

## Abstract

Ded1 is an essential DEAD-box helicase in yeast that broadly stimulates translation initiation and is critical for mRNAs with structured 5’UTRs. We have evaluated the proposal that Ded1 stimulates translation primarily by preventing initiation at upstream ORFs (uORFs) associated with stable secondary structures. By Ribo-Seq analysis under experimental conditions designed to suppress artifactual 5’UTR translation, we found that reduced translation of the main open-reading-frames (mORFs) in native mRNAs is generally not accompanied by increased 5’UTR translation in *ded1* mutant cells, and that the presence of translated uORFs in yeast mRNAs generally does not confer heightened dependence on Ded1 for efficient translation of mORFs. Results from a high-throughput reporter assay examining native 5’UTRs reinforce the importance of Ded1 in initiation from structured 5’ UTRs and show that impairing Ded1 has minimal effects on translational repression by uORFs. Our results demonstrate that, in cells growing vegetatively in rich medium, translational stimulation by suppression of inhibitory uORFs is restricted to a minority of Ded1 targets, and that unwinding of 5’ UTR secondary structures *per se* is the principal mechanism for Ded1 stimulation of translation initiation.

## INTRODUCTION

Most eukaryotic mRNAs are translated by the scanning mechanism, which begins with assembly of a 43S preinitiation complex (PIC) containing the small (40S) ribosomal subunit, a ternary complex (TC) of eukaryotic initiation factor 2 (eIF2), Met-tRNA_i_^Met^, and GTP, and factors eIF1, eIF1A, eIF5, and eIF3. The 43S PIC binds to the 5′-end of mRNA, with eIF4F bound to the m^7^G cap (comprised of cap-binding protein eIF4E, scaffolding subunit eIF4G, and DEAD-box RNA helicase eIF4A), and scans the 5′-untranslated region (UTR) to identify the start codon. The 48S PIC joins with the large (60S) subunit to form an 80S initiation complex ready to begin protein synthesis (reviewed in (1,2)). Both 43S PIC attachment to mRNA and subsequent scanning are inhibited by secondary structures in the mRNA 5’UTR (3) thought to be resolved by DEAD-box RNA helicases. The association of eIF4A with eIF4F should facilitate unwinding of cap-proximal structures; however, other helicases, including Dhx29 in mammals and Ded1/Ddx3 in budding yeast, are additionally required to resolve cap-distal structures that impede scanning (reviewed in (4,5)).

eIF4A is required for 48S PIC assembly in vitro for all mRNAs tested regardless of their structural complexity (6,7). Evidence suggests that mammalian eIF4A remodels the 40S subunit to enhance PIC attachment (8), and might facilitate threading of the 5’-end of mRNA into the 40S entry channel (9). Ribosome footprint profiling of a yeast eIF4A temperature-sensitive mutant (*tif1-ts)* revealed that, despite a strong reduction in bulk polysome assembly, inactivation of eIF4A markedly reduced the relative translational efficiencies (TEs) of less than 40 mRNAs (10), suggesting that the majority of mRNAs have similar requirements for eIF4A in yeast cells. In the same study, inactivation of Ded1 in a cold-sensitive Ded1 mutant (*ded1-cs*) conferred a comparable large reduction in bulk translation but produced a much broader reprogramming of translation, reducing the relative TEs of >1100 mRNAs. These Ded1-hyperdependent mRNAs displayed a marked tendency for long, structure-prone 5’UTRs, suggesting that Ded1 plays a larger role than eIF4A in promoting translation of mRNAs harboring stable 5’UTR structures (5,10). This mRNA-specific role was even more evident when Ded1 was inactivated in a mutant lacking the closely related paralog Dbp1, conferring more widespread and severe reductions in relative TEs of the Ded1-hyperdependent mRNAs than observed in temperature-sensitive *ded1-ts* mutant cells containing Dbp1. Profiling of 40S subunits showed that both the attachment of 43S PICs and subsequent scanning to the AUG start codon were preferentially impaired on Ded1-hyperdependent mRNAs in *ded1-cs* and *ded1-ts dbp1Δ* mutants (5). The heightened requirement for Ded1 to stimulate assembly of 48S PICs on Ded1-hyperdependent mRNAs with structured 5’UTRs was reconstituted in a purified yeast translation initiation system. Whereas Ded1 accelerated PIC assembly on all mRNAs tested, it had a much stronger effect on the Ded1-hyperdependent vs. Ded1-hypodependent transcripts, and the greater stimulation of several Ded1-hyperdependent transcripts was eliminated by destabilizing stem-loop structures in their 5’UTRs (11). Interestingly, Ded1 stimulation of 48S PIC assembly in this in vitro system was enhanced by eIF4F and by domains in eIF4G and Ded1 that mediate Ded1-eIF4F association (11), including Ded1 N-terminal amino acids necessary for its interactions with eIF4A and eIF4E in vivo (12). These last findings support previous biochemical evidence that Ded1 unwinding activity on a model substrate is stimulated by eIF4A and eIF4G (13).

It was first established by Kozak that secondary structures can enhance rather than impair translation initiation when positioned immediately downstream of a poorly recognized start codon, including AUGs in poor context and near-cognate (non-AUG) start codons (NCCs), most likely by increasing the dwell time of the scanning PIC at the poor start codon (14). This mechanism of translational control, which has been dubbed START (<u>st</u>ructure-<u>a</u>ssisted <u>R</u>NA translation”) initiation (15), was implicated in promoting initiation at an upstream AUU codon in *PTEN* mRNA to produce the PTENβ isoform in human cells (16). More recently, it was reported that yeast Ded1 unwinds such 5’UTR structures to suppress initiation at upstream open-reading-frames (uORFs) that would otherwise inhibit initiation at downstream coding sequences, as the principal mechanism of translational stimulation by Ded1 in vivo. Guenther et al. reached this conclusion based on a combination of (i) Ribo-Seq analysis of the temperature-sensitive (Ts^-^) *ded1-95* mutant interpreted to indicate widespread increased 5’UTR translation correlated with decreased translation of downstream main open reading frames (mORFs), (ii) in vivo DMS mapping of RNA structures in *ded1-95* cells suggesting enrichment of Ded1-resolved structures located 3’ of upstream alternative translation initiation sites (ATISs) whose translation is suppressed by Ded1, (iii) iCLIP data suggesting enrichment of Ded1 binding sites in proximity to ATISs and sites of Ded1 unwinding, and (iv) evidence that eliminating the ATISs from two native mRNAs diminished their reduced translation in *ded1* cells (17). However, mRNAs containing high-stringency ATISs had only average reductions in translational efficiency (TE) compared to all mRNAs in *ded1-95* cells (17), making it unclear whether this uORF-dependent mechanism of Ded1 function, referred to below as the Ded1-START model, applies broadly in the yeast translatome.

We recently employed a purified system for genome-wide analysis of translation initiation (Rec-Seq) and reconstituted Ded1 function in preferentially stimulating assembly of 48S PICs on hundreds of native mRNAs judged to be hyperdependent on Ded1 for efficient translation in vivo on the basis of prior ribosome profiling (Ribo-Seq analysis) of *ded1-cs* cells. In this reconstituted system, Ded1 preferentially promoted 48S PIC formation at the main start codons of mRNAs with long, structured 5’UTRs. Although Ded1 enhanced leaky scanning at numerous uORFs, consistent with the conclusions of Guenther et al. summarized above, the magnitude of this effect was far too small to account for the observed Ded1 stimulation of PIC assembly at mORF start codons downstream (18). These in vitro data challenged the notion that Ded1 broadly stimulates initiation primarily by suppressing translation of uORFs. However, because these last findings were obtained using a minimal translation initiation system it remains possible that, in cells, uORFs would be more inhibitory or that Ded1 would have a greater effect in promoting leaky scanning of uORF start codons than was observed in vitro.

Accordingly, we have addressed the applicability of the Ded1-START model in vivo using two complementary approaches. First, we carried out new Ribo-Seq experiments on *ded1* mutants, omitting cycloheximide treatment of cells to avoid artifactually enhancing ribosome occupancies of uORFs (19), and examined (i) whether reductions in translational efficiencies (TEs) of mORFs conferred by impairing Ded1 are generally associated with increased translation of uORFs in the affected mRNAs; and (ii) whether mRNAs with repressive uORFs show a heightened dependence on Ded1 for efficient translation of their mORFs (Fig. 1A). Second, we carried out a massively parallel reporter analysis (MPRA) called FACS-uORF that simultaneously compares YFP expression from thousands of reporters containing endogenous 5’UTRs harboring uORFs to the expression of the corresponding mutant reporters lacking uORF start codons. By this approach, we previously identified ∼560 mRNAs containing functional AUG-initiated uORFs, and ∼190 mRNAs with functional uORFs initiated with near-cognate start codons (NCCs), whose presence in the 5’UTRs significantly affected YFP expression from the downstream mORFs (20). Here, we have conducted FACS-uORF analysis of isogenic *ded1* and wild-type (WT) *DED1* cells to determine whether repressive uORFs inhibit translation of the downstream mORFs to a greater extent when Ded1 function is impaired (Fig. 1B) in the manner expected from the reduced leaky scanning of uORFs in *ded1* cells predicted by the Ded1-START model. Our combined results from these two orthogonal approaches indicate that Ded1 generally stimulates translation of mORFs independently of the presence of inhibitory uORFs and that, at least in non-stressed yeast cells, the Ded1-START model appears to operate only on a small number of mRNAs.

**Figure 1.**
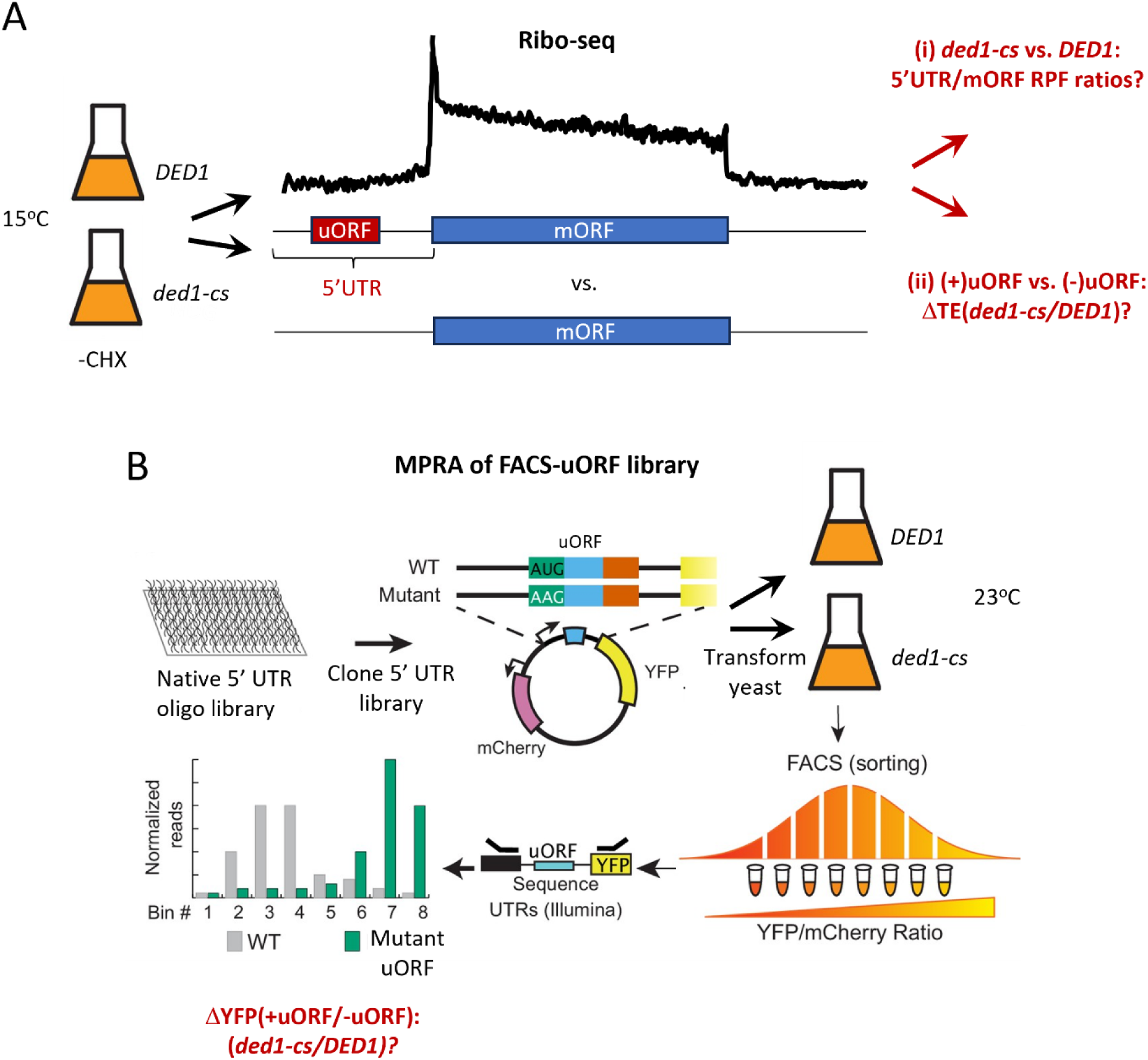
Workflow of orthogonal analyses of Ded1 regulation of uORF-mediated translational control. **(A)** Ribo-Seq analysis of *DED1* and *ded1-cs* cells cultured at the nonpermissive temperature of 15°C without treating cells with cycloheximide (-CHX) to determine (i) if mRNAs with uORFs exhibit increased densities of ribosome protected fragments (RPFs) in 5’UTRs coupled with decreased RPF densities in the corresponding mORFs in *ded1-cs* vs. *DED1* cells, and (ii) if mRNAs with translated uORFs exhibit larger reductions in translational efficiencies (TEs) in *ded1-cs* vs. *DED1* cells compared to mRNAs lacking uORFs, all as predicted by the Ded1-START model. **(B)** MPRA of a plasmid library of YFP reporters containing native 5’UTRs harboring translated uORFs and the corresponding mutant uORFs lacking start codons in *DED1* and *ded1-cs* cells cultured at the semi-permissive temperature of 23°C. Transformants of each strain are sorted *en masse* by fluorescence-activated cell sorting (FACS) into bins of different YFP expression levels and the reporter 5’UTR sequences present in each bin determined by deep-sequencing of DNA amplified from pooled plasmids. Idealized results are depicted for a repressive uORF that confers lower YFP expression in its presence vs. absence. Ratios of YFP expression (+uORF/-uORF) are compared in the two strains to determine if repressive uORFs are generally more inhibitory in *ded1-cs* vs. *DED1* cells, as predicted by the Ded1-START model.

## MATERIALS AND METHODS

### Yeast Strains

Yeast strains containing WT or mutant alleles of *DED1* isogenic to strain BY4741 were constructed previously (10) and are described in Table 1. The *ded1-120* allele, referred to below as *ded1-cs*, encodes amino acid substitutions of Gly108/Gly494 with Asp (G108D/G494D), whereas *ded1-952*, referred to as *ded1-ts,* encodes substitutions T408I/W253R. G494D likely impairs conserved subdomain VI function in ATP binding or hydrolysis, whereas T408I in subdomain IV probably affects RNA binding (21). (Our previous DNA sequence analysis (10) revealed that *ded1-952* differs from the *ded1-95* allele examined by Guenther et al. (2018) by encoding the W253R substitution in addition to T408I). The *ded1-952* mutation does not affect Ded1 protein abundance at the restrictive temperature (12), while it is unknown if the same holds for *ded1-120*.

**Table 1.** Yeast strains used in this study.

| Strain | Genotype | Source or reference |
| --- | --- | --- |
| NSY4 | *MATa his3Δ1 leu2Δ0 lys2Δ0 met15Δ0 ura3Δ0 ded1Δ::kanMX4*<br>p5467 (*DED1 LEU2*) | (10) |
| NSY5 | *MATa his3Δ1 leu2Δ0 lys2Δ0 met15Δ0 ura3Δ0 ded1Δ::kanMX4*<br>p5466 (*ded1-120 LEU2*) | (10) |
| NSY10 | *MATa his3Δ1 leu2Δ0 met15Δ0 ura3Δ0 ded1-952::kanMX4*<br>p5457 (*DED1 HIS3*) | (10) |
| NSY11 | *MATa his3Δ1 leu2Δ0 met15Δ0 ura3Δ0 ded1-952::kanMX4*<br>p1372 (*HIS3*) | (10) |

### Ribosome footprint profiling and RNA-Seq

Ribo-Seq was conducted essentially as described previously (5) as detailed below, on isogenic yeast strains NSY4 (*DED1*) and NSY5 (*ded1-120/ded1-cs* here); NSY10 (*DED1*) and NSY11 (*ded1-952/ded1-ts*) using two biological replicates for each genotype, except that cells were not treated with cycloheximide before harvesting, and cycloheximide was added to the lysis buffer at 5x the standard concentration, referred to below as “-CHX” conditions.

Yeast strains growing exponentially in synthetic complete (SC) medium at 30°C were harvested by centrifugation at room temperature and resuspended in SC medium at 37°C or 15°C (depending on which *ded1* allele was involved in the experiment). NSY4 (*DED1*) and NSY5 (*ded1-cs*) were grown in SC-Leu medium under conditions similar to those employed previously (10) including a temperature shift at 15°C for 10 min. Similarly, NSY10 (*DED1*) and NSY11 (*ded1-ts*) were grown in SC-His medium under conditions similar to those employed previously (10) including a temperature shift at 37°C for 2 h. Cells were harvested by vacuum filtration at room temperature, without prior treatment with cycloheximide, and quick-frozen in liquid nitrogen. Cells were lysed in a freezer mill with lysis buffer (20 mM Tris [pH 8.0], 140 mM KCl, 1.5 mM MgCl_2_, 1% Triton, 500 µg/mL cycloheximide). For ribosome footprint library preparation, 30 A_260_ units of extract were treated with 450 U of RNAse I (Ambion, #AM2295) at 25°C for 1 hr on a thermomixer at 700 rpm, and ribosomes were pelleted by centrifugation on a 1M sucrose cushion. Ribosome-protected mRNA fragments (RPFs) were purified using a miRNeasy Mini kit (Qiagen) per the vendor’s instructions and 25–34 nt RPFs were size-selected after electrophoresis through a 15% TBE-Urea gel. The purified RPFs were ligated to universal miRNA cloning linker (Synthesized by Integrated DNA Technologies) using truncated T4 RNA ligase 2 (New England Biolabs; M0242L) after a dephosphorylation reaction carried out with T4 Polynucleotide kinase (PNK; New England Biolabs, M0201L). The ligated RPF products were size-selected after electrophoresis on a 10% TBE-Urea gel, reverse-transcribed using Superscript III (Invitrogen; 18080044), size-selected again, and circularized using CircLigase ssDNA Ligase (Epicentre; CL4111K). Ribosomal RNA contamination was reduced by oligonucleotide subtraction hybridization. Each ‘subtracted’ library was amplified by PCR to add unique six nt indexes and common Illumina primer and flow cell binding regions. Quality of the libraries was assessed with a Bioanalyzer using the High Sensitivity DNA Kit (Agilent 5067–4626) and quantified by Qubit. Sequencing was done on an Illumina HiSeq system at the NHLBI DNA Sequencing and Genomics Core at NIH (Bethesda, MD). For RNA-seq library preparation, total RNA was purified using miRNeasy Mini kit (Qiagen) from aliquots of the same extracts used for RPF library preparation. Polyadenylated mRNA was isolated from total RNA using the Poly(A)Purist MAG Kit (Ambion, #AM1922) and randomly fragmented at 70°C for 8 min in fragmentation reagent (Ambion, #AM8740). Fragment size selection, library generation and sequencing were carried out as above. RPF and RNA-seq libraries were prepared and sequenced from 2 independent cultures (biological replicates) for each pair of *DED1* and *ded1* strains under comparison.

### Analysis of 5’ UTR translation using Ribo-Seq data

Linker sequences were trimmed from Illumina reads and the trimmed fasta sequences were aligned to the *S. cerevisiae* ribosomal database using Bowtie (22). The non-rRNA reads (unaligned reads) were then mapped to the *S. cerevisiae* genome using TopHat (23). Only uniquely mapped reads from the final genomic alignment were used for subsequent analyses. For uORF and 5’UTR TE calculations, we modified a BED file generated previously (24) that contained sequence coordinates of translated uORFs showing evidence of translation in Ribo-Seq data sets from isogenic WT *TIF11* or *tif11-R13P* strains and identified using two separate bioinformatics tools for discovering translated uORFs, both described previously (25,26), as well as sequence coordinates for 5’UTRs, mORFs, and 3’UTRs for every yeast gene, to which we added the coordinates of the ATIS uORFs identified by Guenther et al. (17) and the functional uORFs identified by FACS-uORF MPRA by May et al. (20). The uORFs that overlap between any two datasets are included only once in the final BED file, resulting in coordinate information for 3315 uORFs, which was used to obtain RPF counts for uORFs, 5’UTRs, mORFs and 3’UTRs in each Ribo-Seq dataset examined, excluding the first and last nucleotide triplets of 5’UTRs, and the first 20 codons of mORFs. (Note that the aforementioned uORF compilations used different uORF length criteria: no minimum uORF length (17), uORFs ≥2 codons excluding the stop codon (20), and uORFs ≥3 codons excluding the stop codon (24). Among all of the uORFs identified in the three compilations, only 68 were <3 codons long. The 5’UTR and uORF TEs were calculated as RPF counts in the 5’UTR or uORF normalized to mRNA read counts of the mORF. Statistical analysis of differences in RPF or mRNA read counts, or TE values, between *DED1* and *ded1* cells from Ribo-Seq data obtained for two biological replicates was conducted with DESeq2 (27). For mORF TE calculations, genes with less than 128 total mRNA reads in the 4 samples combined (two replicates of both *DED1* and *ded1* strains under comparison) were excluded from calculation of TE values. Because the vast majority of uORFs were >3 codons, we retained an RPF read-count minimum threshold of 8 for uORFs, and 128 total mRNA reads, in the 4 data sets (two replicates each from *DED1* and *ded1* cells), as we applied previously (24).

For the analysis of uORF containing mRNAs in Figs. S7A-C, we employed the functional uORF compilation from May et al. (20) mentioned above, our previous compilation of translated uORFs from Zhou et al. (28) that is highly similar in derivation and content to that described above from Martin-Marcos et al. (24), and a third compilation of uORFs conserved in sequence or position among different Saccharomyces species and showing evidence of translation in Ribo-Seq data sets (29).

Metagene analysis of RPFs in 5’UTRs, mORFs, and 3’UTRs across the translatome were determined using the R Bioconductor package riboSeqR (version 1.24.0) (http://www.bioconductor.org/packages/riboSeqR). Gene browser depictions of Ribo-Seq and RNA-seq data were obtained using the Integrated Genomics Viewer (IGV) (30). Notched box plots were constructed using a web-based tool at http://shiny.chemgrid.org/boxplotr/, in which the upper and lower boxes contain the second and third quartiles and the band gives the median. If the notches in two plots do not overlap, there is roughly 95% confidence that their medians are different. Scatterplots were created using the scatterplot function in Microsoft Excel. Venn diagrams were generated using the open source program Venn diagram plotter (https://pnnl-comp-mass-spec.github.io/Venn-Diagram-Plotter/) or the on-line tool at https://www.biovenn.nl/, UpSet plots were generated in R using the UpSetR package (31), and the significance of gene set overlaps was evaluated with the hypergeometric distribution using the web-based tool (https://systems.crump.ucla.edu/hypergeometric/index.php).

Center of ribosome density (CRD) was calculated as described previously (32) as the nucleotide position in each mRNA where 50% of footprint reads map upstream or downstream. The shift in CRD in *ded1* mutants was determined as Δ*CRD* = (Average of CRD values from biological replicates of *ded1* mutant − Average of CRD values from biological replicates of *DED1* strain). Structures downstream of uORFs were evaluated by compiling cumulative PARS scores of defined windows of 16-30, 16-45 or 16-60 nucleotides downstream of the uORF start codon for mRNAs containing a single uORF for which coordinates and read coverage are available in the PARS dataset (33).

### FACS-uORF analysis of *ded1-cs* and *DED1* strains

Large-scale yeast transformations were carried out (34) to generate two pools of ∼610,000 and ∼580,000 independent transformants of strains NSY4 (*DED1*) and NSY5 (*ded1-cs*), respectively, containing *YFP* reporter plasmids from the “long 5’UTR” reporter library, and two pools of ∼390,000 and ∼416,000 transformants, respectively, for the “short 5’UTR” library, constructed previously in plasmid *pGM-ENO2-YFP-mcherry* (20). Transformants were pooled, cultured overnight in 200 mL of SC medium lacking leucine and uracil (SC-L-U), harvested, and resuspended in twenty 1-mL aliquots of 15% glycerol and stored at -80°C. Three replicate cultures were prepared for each strain by inoculating 200 mL of SC-L-U with 500 µl each of the four glycerol stocks of library transformants and growing overnight at the permissive temperature of 30 °C. Three 50-mL cultures of SC-L-U were inoculated with 0.1 OD_600_ of the overnight cultures and incubated at the semi-permissive temperature of 23°C for 3 doublings (∼0.7-0.8 OD_600_). FACS was conducted as described previously (20) to sort cells into eight bins of increasing YFP/mCherry fluorescence. Cells from each bin were cultured in 5 ml of SC-L-U at 30°C overnight. To verify that the cells were sorted into the correct bins, the ratio of YFP and mCherry fluorescence was analyzed using a Tecan M1000 plate reader. DNA libraries were prepared for each bin essentially as described by May et al. (20). In brief, plasmids were isolated from the sorted yeast cells using Zymoprep yeast plasmid miniprep II kit and three rounds of PCR were used to amplify the library plasmids and add Illumina sequences and barcodes for DNA sequencing. For each uORF, mean YFP values from wildtype and mutant UTRs were compared using the Wilcoxon Rank-sum Test (WRT) in each replicate, and the Benjamini-Hochberg correction was applied to control the FDR at 5%. uORFs with significant (FDR <0.05) and consistent (same direction) effects on YFP levels were considered as significant regulators of downstream translation.

### Construction of YFP reporter plasmids and reporter assays in yeast

DNA fragments containing the appropriate 5’UTR sequences, either WT or with an uORF initiation codon altered to AAG, joined to the *ENO2* promoter sequence and containing Avr II and Bgl II restriction sites added to the 5’ and 3’ ends, respectively, were synthesized by Twist Biosciences and sequenced in their entirety. The synthesized fragments were digested with Avr II and Bgl II and inserted upstream of the *YFP* coding sequences of Avr II/Bgl II-digested plasmid *pGM-ENO2-YFP-mcherry* (20). Yeast strains NSY4 and NSY5 were transformed with the resulting plasmids and 3 independent transformants for each plasmid were grown overnight at 30 °C in SC-L-U media, after which 0.1 OD_600_ was used to inoculate 5 mL of SC-L-U and incubated at 23 °C for three doublings. YFP/mCherry fluorescence of whole cells was measured using Tecan M1000 PRO plate reader in top-read mode with optimal gain. Excitation/emission wavelengths were set to 512/532 nm for YFP and 586/608 nm for mCherry each with a 5 nm bandwidth.

## RESULTS

### Reduced translation of downstream mORFs is generally not accompanied by increased uORF translation in *ded1* mutants

A previous study led to the conclusion that Ded1 generally reduces translation initiation at uORFs in order to promote translation at mORF start codons. Key evidence supporting this uORF-dependent mode of Ded1 function—the Ded1-START model—came from Ribo-Seq analysis of the Ts^-^ *ded1-95* mutant in which elevated 5’UTR translation appeared to be associated with decreased translation of the downstream mORFs (17). An important caveat to these profiling data was the pre-treatment of cells with cycloheximide (CHX), rather than adding the inhibitor exclusively to the lysis buffer, to arrest translating ribosomes on the mRNA and enable the isolation of RPFs by RNase treatment. Adding CHX to cells can artifactually increase 80S ribosome occupancies near the start codons of mORFs and in 5’UTRs, particularly under stress conditions (19). We addressed whether this artifact influenced conclusions about the relationship between translation of 5’UTRs versus mORFs by repeating Ribo-Seq analysis of *ded1-cs* and *ded1-ts* mutants under the same growth conditions we used previously to examine these mutants (5) but omitting CHX treatment of cells and increasing by 5-fold the CHX concentration added to lysates to minimize the known artifacts associated with its use. As previously, we employed DESeq2 analysis of the biological replicates of the Ribo-Seq and RNA-Seq analyses to identify mRNAs exhibiting statistically significant changes in relative 80S ribosome densities, computed by normalizing the number of RPF reads to mRNA reads for each transcript, as a measure of relative TE. While TEs measured by RiboSeq in this way are relative to the average gene and not absolute values, we refer to them below simply as TEs.

Eliminating CHX treatment of cells (-CHX) had relatively little effect on the changes in TEs in *ded1* mutant vs. *DED1* cells determined for the majority of mORFs compared to our earlier studies where cells were pre-treated with the drug (+CHX) (Fig. S1A-B). We found highly similar groups of ∼1400-1500 mRNAs showing significant reductions in relative TE in the *ded1-cs* mutant vs. *DED1* strain (>1.5-fold at FDR<0.05) in the +CHX and -CHX datasets (Fig. S1C). We classified these as Ded1-hyperdependent (1.5-fold down in *ded1-cs*) and Ded1-hypodependent (1.5-fold up in *ded1-cs*). Although we identified roughly twice as many Ded1-hyperdependent mRNAs in the *ded1-ts* mutant in the new -CHX experiment, they included nearly all mRNAs identified in our previous +CHX experiment (Fig. S1D). The greater number of Ded1-hyperdependent mRNAs may reflect the ∼5-fold greater read-depth of the -CHX versus +CHX experiments for the *ded1-ts* mutant (Fig. S2A), increasing our ability to discern statistically significant TE changes across many mRNAs. In view of the strong overlap between the groups of Ded1-hyperdependent mRNAs identified in the -CHX and +CHX experiments, the new -CHX data can be interrogated to provide a more reliable evaluation of the effects of *ded1* mutations on 5’UTR translation. Unless mentioned otherwise, the -CHX data were used for all subsequent analyses.

In accordance with previous findings (19), eliminating CHX pre-treatment greatly decreased the occupancy of 80S ribosomes in 5’UTRs relative to the corresponding mORFs in both *DED1* and *ded1* strains (Fig. S2B, brown vs. grey). This outcome eliminated the increase in the proportion of 5’UTR RPFs that we found previously for the *ded1-ts* mutant vs. *DED1* strain in +CHX experiments (Fig. S2B, cols. 1-2, brown vs. grey). Nevertheless, in the new -CHX data we observed hundreds of mRNAs showing statistically significant >2-fold increases in relative TEs of their resident translated uORFs, calculated as the sum of RPFs in the uORFs normalized by the transcript abundance determined by RNA-Seq, in each of the *ded1* mutants compared to the *DED1* strain (Fig. S2C-D). For this last analysis, we interrogated a compilation of 3315 uORFs that either showed evidence of translation in previous Ribo-Seq data (24), were identified in our previous FACS-uORF analysis of functional uORFs, or were identified as Ded1-suppressed ATISs by Guenther et al. (17) (See Materials and Methods for additional details). We reasoned that interrogating a combination of different sets of uORFs identified by distinct criteria in this way would provide a comprehensive examination of the effect of inactivating Ded1 on uORF translation. We applied DESeq2 analysis of our RiboSeq data to assess statistically significant changes in TE of these uORFs and the downstream mORFs encoded on the same transcripts. For the 2661 uORFs with sufficient data coverage for analysis, 956 showed ≥2-fold increases in TE in *ded1-cs* cells versus *DED1* cells [uORF ΔTE*_ded1-cs_*_(-CHX)_] at FDR <0.01, and 555 uORFs showed increased TEs of this same magnitude in *ded1-ts* versus *DED1* cells, with a highly significant overlap of the two groups of uORFs (Fig. S2E). These results show that, although eliminating CHX pre-treatment reduces ribosome occupancy of 5’ UTRs, impairing *DED1* function increases the relative translational efficiencies of many uORFs.

We next asked whether these apparent increases in uORF translation on inactivating *DED1* are associated with decreased translation of the downstream mORFs. This would be expected if Ded1 predominantly enhances mORF translation by suppressing uORF translation, as proposed in the Ded1-START model. In this case, most genes whose translation is hyperdependent on Ded1 would be expected to show larger increases in uORF translation compared to Ded1-hypodependent genes in *ded1-cs* cells. At odds with this prediction, we found that Ded1-hyperdependent mRNAs exhibit smaller, rather than larger, increases in TE of their resident uORFs compared to the uORFs in Ded1-hypodependent mRNAs in *ded1-cs* vs. *DED1* cells (Fig. 2A, cols. 2-3). The 1074 uORFs present in the 410 Ded1-hyperdependent mRNAs also show smaller TE increases compared to all uORFs in *ded1-cs* cells (Fig. 2A, cols. 1-2). The same trends held after applying less stringent thresholds for defining Ded1-hyper/hypo-dependence in the *ded1-cs* data for the uORF-containing mRNAs (Fig. 2A, cols. 4-6). Expanding this last analysis, we divided Ded1 hyperdependent mRNAs containing uORFs into 11 bins according to their mORF TE changes conferred by *ded1-cs* (Fig 2B). The mRNAs with the greatest reductions in mORF TE (most Ded1-hyperdependent) contained in the first two bins showed the smallest median increases in uORF TEs among all bins (Fig. 2B)—opposite the expectation of the Ded1-START model. We also examined the two groups of mRNAs with the largest increases or decreases in uORF TEs in the *ded1-cs* mutant and compared their corresponding mORF TE changes. Consistent with results above, the mRNAs whose uORF TEs increased the most in the mutant have somewhat larger increases in mORF TE compared to all uORF-containing mRNAs (Fig. 2C, cols. 1 vs 2), while the group containing uORFs exhibiting the greatest TE reductions showed greater than average reductions in mORF TE (Fig. 2C, cols. 1 vs 3). These positive correlations between uORF and mORF TE changes conferred by *ded1-cs* were also observed when a relaxed FDR stringency for significant uORF TE changes was applied (Fig. 2C, cols. 4-6). Similar results were obtained by analyzing the *ded1-ts* mutant profiling data (cf. Figs. S3A-B vs. Figs. 2A & C). Finally, changes in both RPFs and TEs for uORFs are positively, rather than negatively, correlated with the corresponding changes for mORFs conferred by *ded1-ts* or *ded1-cs* for all expressed mRNAs (Fig. 2D). Overall, these results contradict the model that decreased translation of the mORF is generally linked to increased translation of uORFs on impairing Ded1 function. Our observations of a positive, rather than negative, correlation between uORF and mORF TE changes might reflect altered PIC attachment to the 5’UTR conferred by the *ded1* mutations, leading to changes in uORF and mORF translation in the same direction on the affected mRNAs. Indeed, we provided evidence from 40S profiling that Ded1 stimulates 43S PIC loading as well as subsequent 5’UTR scanning on many Ded1-hyperdependent mRNAs *in vivo* (5).

**Figure 2.**
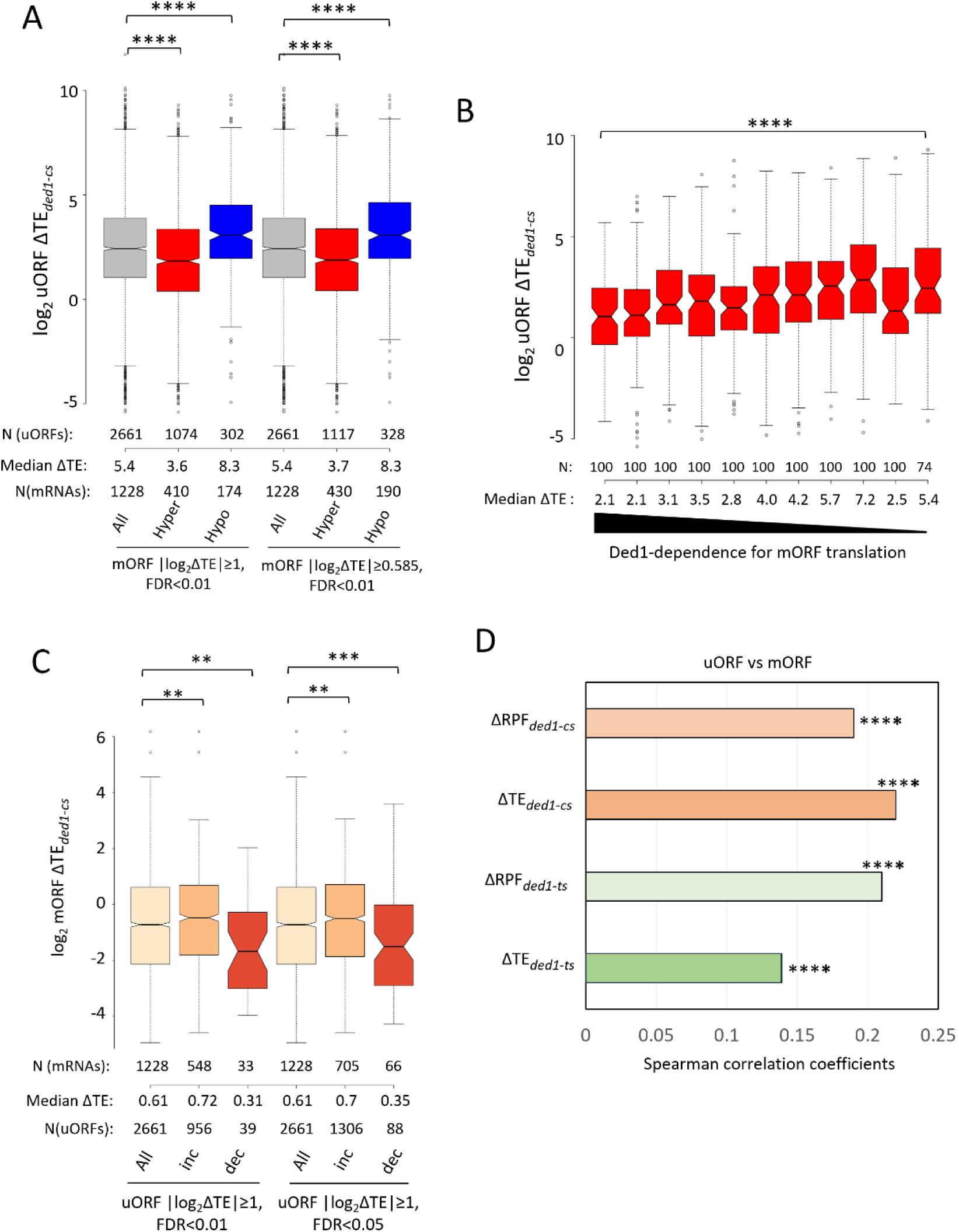
Increased uORF translation in *ded1* mutant cells is generally not accompanied by reduced translation of the downstream mORFs. **(A)** Notched box plots comparing the distributions of log_2_ fold-changes in uORF TE in *ded1-cs* vs. *DED1* cells (uORF ΔTE*_ded1-cs_*) from cultures untreated with CHX prior to Ribo-Seq analysis for all 1228 mRNAs containing uORFs (col. 1), the subset of 410 mRNAs exhibiting ≥2-fold decreases in mORF TE (Hyper, col. 2), and the subset of 174 mRNAs exhibiting ≥2-fold increases in mORF TE (Hypo, col. 3) in *ded1-cs* vs. *DED1* cells at FDR < 0.01, ie. with absolute values of log2ΔTE values ≥ unity (|log_2_ΔTE| ≥1). Results for mRNAs exhibiting ≥1.5-fold decreases (col. 5) or increases (col. 6) in mORF TEs (ie. |log_2_ΔTE| ≥0.585) in *ded1-cs* vs. *DED1* cells at FDR < 0.01 are presented similarly in cols. 4-6. Each box depicts the interquartile range containing 50% of the data, intersected by the median; the notch indicates a 95% confidence interval (CI) around the median. Numbers of uORFs in each mRNA group are shown above the x-axis and the unlogged median uORF TE changes and numbers of mRNAs in each group are shown below the x-axis. **(B)** Notched box plots showing distributions of log_2_ uORF TE changes in *ded1-cs* versus *DED1* cells in the -CHX condition for the 1074 uORFs present in the 410 Ded1-hyperdependent mRNAs (from panel A, col. 2) sorted into bins of 100 uORFs (except for the last bin with 74 uORFs) on the basis of TE reductions conferred by *ded1-cs* for the corresponding mORFs, arranged left to right from largest to smallest reductions. **(C)** Notched box plots comparing the distributions of log_2_ fold-changes in mORF TE observed in *ded1-cs* versus *DED1* cells from -CHX cultures (mORF ΔTE*_ded1-cs_*) for all uORFs (col. 1) or uORFs exhibiting ≥2-fold increases (inc, col. 2) or decreases (dec, col. 3) in TE (ie. |log_2_ΔTE| ≥1) in *ded1-cs* vs. *DED1* cells at FDR < 0.01. Results for uORFs exhibiting ≥2-fold increases or decreases in TE in *ded1-cs* cells at FDR < 0.05 are depicted similarly in cols. 4-6. Results of Mann-Whitney U-tests are summarized for panels A-C as: ****, P<0.0001; **, P<0.01. **(D)** Spearman coefficients for correlations between changes in uORFs versus the corresponding mORFs of either RPFs or TEs conferred by the indicated *ded1* mutation for all mRNAs with data available for both RPFs and TEs, including >2660 uORFs present in >1200 mRNAs, with P-values for the correlation analyses indicated (****, P<0.0001).

Previously, Guenther et al. provided evidence supporting widespread Ded1 suppression of uORF translation using the “center of ribosome density” (CRD) metric, which is the nucleotide position in each mRNA where 80S RPFs are divided equally between upstream and downstream locations (32). Increased 5’UTR translation in *ded1* cells should shift the CRD towards the mRNA 5’ end, and the difference between CRD position in *ded1* versus *DED1* cells, normalized to mRNA length, should yield negative ΔCRD values. Similar to the findings reported for the *ded1-95* mutant (17), we observed a modest positive correlation between TE changes for mORFs and ΔCRD values for all mRNAs in both *ded1-cs* and *ded1-ts* mutants using our previous +CHX data (Fig. S4A-B). However, an upstream shift in the CRD can result from an artifact of CHX treatment prior to cell lysis wherein 80S ribosomes accumulate at the beginnings of mORFs (19). Indeed, examining metagene plots of our Ribo-Seq data for all expressed genes reveals accumulation of RPFs at the start codon and 5’ ends of mORFs in the +CHX but not the -CHX datasets in both *ded1-ts* and *DED1* cells cultured at the restrictive growth temperature (Fig. S5A(i) vs. (ii)). The increases in RPF densities at the 5’ ends of mORFs in the +CHX conditions are much greater in magnitude than the total RPF densities in 5’UTRs, indicating that they will strongly influence calculations of CRD values. Similar findings were made for *ded1-cs* and *DED1* cells (Fig. S5B(i)-(ii)). Importantly, a positive correlation between TE changes for mORFs and ΔCRD values is absent for the *ded1-ts* mutant, and is considerably weaker for the *ded1-cs* strain, in the -CHX vs. +CHX conditions (Fig. S4, C-D vs. A-B). The weak correlation between TE changes and ΔCRD values still evident for the *ded1-cs* -CHX results (Fig. S4C) is inconsistent with the positive correlation between the TE changes of uORFs vs. mORFs observed for this same dataset (Fig. 2D, row 2). Because the CRD does not report exclusively on 5’UTR translation and can be influenced by CHX-induced artifactual RPF occupancies at the 5’ ends of mORFs (Fig. S5A-B), it is an unreliable indicator of altered uORF translation in +CHX Ribo-Seq data.

### AUG-initiated uORFs generally do not confer hyperdependence on Ded1 for translation of downstream mORFs

A second prediction of the Ded1-START model, that Ded1 mainly promotes translation initiation by suppressing initiation at inhibitory uORFs, is that the presence of uORFs should frequently exacerbate the requirement for Ded1 for efficient translation of the downstream mORFs. To test this prediction using our -CHX datasets, we examined whether Ded1-hyperdependent mRNAs are enriched for translated uORFs, and whether uORF-containing mRNAs show larger than average TE reductions in the *ded1* mutants. In addition to the two uORF compilations mentioned above identified functionally by FACS-uORF analysis or using evidence of translation from RiboSeq data, we examined a third group showing both evidence of translation in Ribo-Seq and evolutionarily sequence conservation among related yeasts (29). After omitting subsets of these mRNA groups that contain both AUG and NCC uORFs, we found that the resulting groups containing uniquely AUG or NCC uORFs display highly significant overlaps with each other, and with the Ded1-suppressed ATISs identified by Guenther et al. (Fig. S6A-B).

Interrogating the groups of Ded1-hyperdependent mRNAs identified in the -CHX datasets above, dubbed TE_down_*ded1-cs* and TE_down_*ded1-ts*, we found that neither group is enriched for the three sets of mRNAs harboring AUG-uORFs identified by different criteria (Fig. S7A & S7C((i)-(ii) & (v)-(vi)); whereas both Ded1-hyperdendent groups were significantly enriched for all three sets of mRNAs with NCC uORFs (Fig. S7B & S7C((iii)-(iv) & (vii)-(viii)). Despite the statistical enrichment for NCC uORFs, the overwhelming majority of Ded1-hyperdependent mRNAs (TE_dn_*ded1-cs* or TE_dn_*ded1-ts* transcripts) do not contain either AUG- or NCC-uORFs, ostensibly at odds with the proposition that Ded1 generally stimulates mORF translation by suppressing inhibitory uORFs. It should be noted however that, according to the Ded1-START model, some Ded1-hyperdependent mRNAs could harbor uORFs that are only translated at detectable levels in *ded1* mutant cells, in which Ded1-suppression of their translation would be impaired, and hence would be missing from our compilations of translated uORFs that were identified in *DED1* cells containing fully functional Ded1.

Consistent with the results above, none of the three sets of mRNAs with AUG uORFs, but all three sets with NCC uORFs, showed significant reductions in median TE in the *ded1-cs* mutant (Fig. S8A, cols. 2,4,6 vs. 1,3,5). After combining the three sets of uORF-containing mRNAs, we again found that the resulting group of 1047 NCC uORF-containing mRNAs, but not the 638 AUG uORF-containing mRNAs, shows a reduction in median TE in *ded1-cs* cells compared to all mRNAs (Fig. 3A, cols. 1-3). The same conclusion emerged for the group of 1374 Ded1-hyperdependent mRNAs (TE_down_*ded1-cs*) that contain either AUG or NCC uORFs, with only the latter showing decreased mORF TE in *ded1-cs* cells compared to all hyperdependent mRNAs (Fig. 3A, cols. 4-6). Very similar results were obtained on analyzing the *ded1-ts* data (cf. Figs. S8B vs. Fig. 3A). Together, these findings suggest that the presence of translated AUG uORFs generally does not exacerbate the effect of *ded1* mutations in reducing translation of the mORF—at odds with the Ded1-START model—whereas translated NCC uORFs might do so.

**Figure 3.**
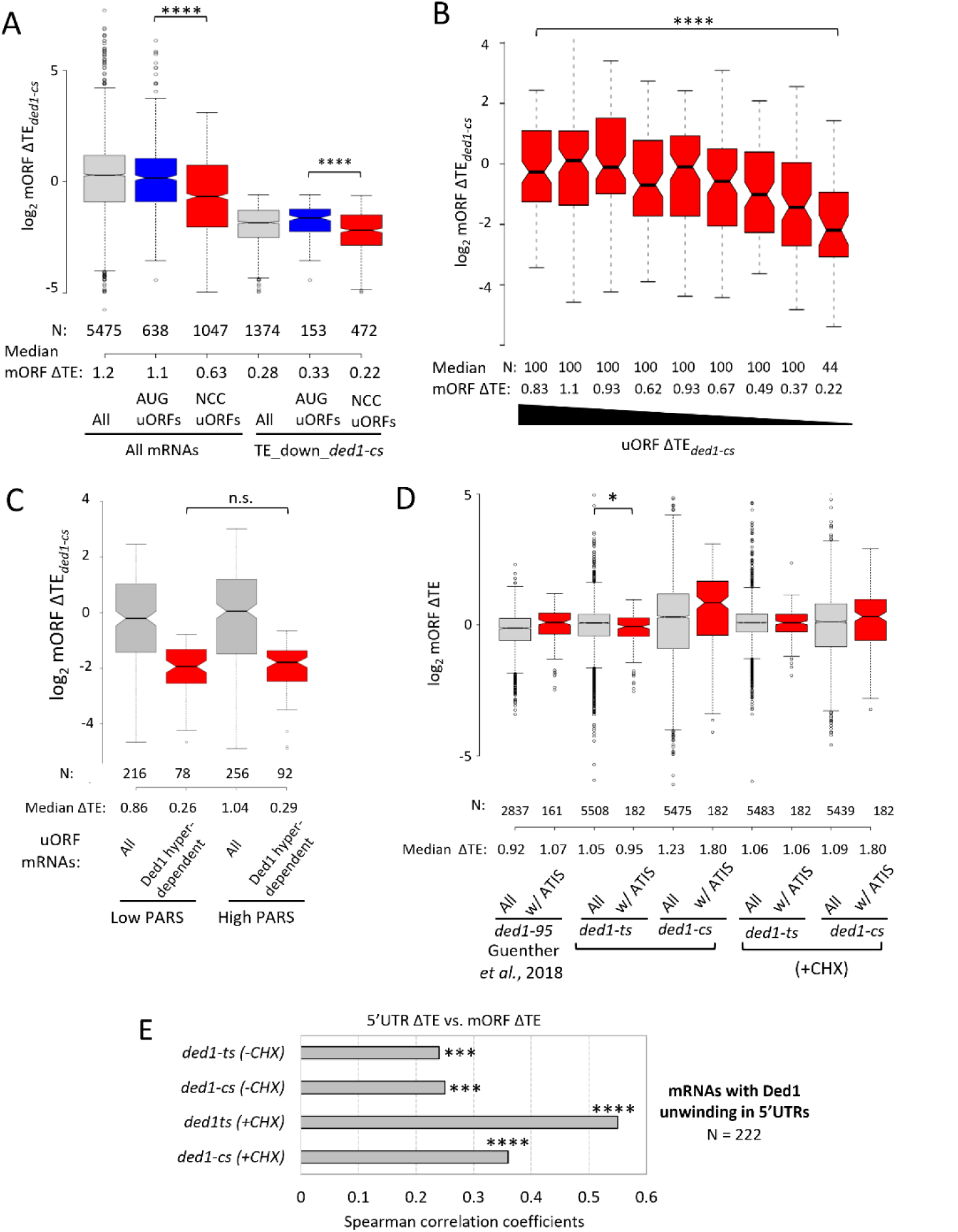
Ded1-hyperdependence conferred by *ded1* mutations is largely independent of increased AUG- or NCC-initiated uORF translation and secondary structures located 3’ of uORFs. **(A)** Notched box plot analysis of the log_2_ fold-changes in mORF TEs observed in *ded1-cs* versus *DED1* cells from -CHX cultures for all mRNAs, the sets of mRNAs containing unique AUG uORFs, NCC uORFs, or lacking (-) uORFs that were compiled from the datasets of May *et al.*, Spealmann *et al.* and Zhou *et al*. (as detailed in Results) for either all mRNAs (cols. 1-4), or from the 1374 TE_down_*ded1-cs*(-CHX) mRNAs. Median ratios are given below the x-axis, and numbers of mRNAs in each group are above the x-axis. (B) Notched box plots showing distributions of ΔTE*_ded1-cs_*_(-CHX)_ for the mRNAs containing unique NCC uORFs (from panel A, col. 3) sorted into bins of 100 mRNAs (except for the last bin with 44 mRNAs) on the basis of uORF TE changes under the same conditions. For each box the unlogged median ratios are given below the x-axis, and numbers of mRNAs in each group are shown above the x-axis. **(C)** Notched box plots comparing the distributions of log_2_ fold-changes in TE observed in *ded1-cs* versus *DED1* cells from -CHX cultures for all mRNAs containing NCC uORFs (grey boxes, cols. 1 & 3) or Ded1-hyperdependent mRNAs containing NCC uORFs belonging to the group TE_down_*ded1-cs*(-CHX) (red boxes, cols. 2 & 4). Two sets of mRNAs for each group with low or high total PARS scores for the interval 16-45 downstream of the +1 nucleotide of the NCC uORFs were analysed. **(D)** Notched box plot analysis comparing the distributions of TE changes in the indicated *ded1* mutant versus *DED1* cells from either -CHX or +CHX datasets for all mRNAs (light grey boxes) or mRNAs containing high stringency alternative initiation sites [ATISs (red boxes)] identified by Guenther *et al* (17). For cols. 1-2, published TE changes in *ded1-95* versus *DED1* cells in +CHX conditions were retrieved and plotted (17). Results of Mann-Whitney U-tests in panels A-D are summarized as: ****, P<0.0001; ***, P<0.001; n.s., not significant, P>0.05. **(E)** Spearman coefficients from correlations between the 5’UTR TE changes versus mORF ΔTE values conferred by the indicated *ded1* mutation versus *DED1* for cells treated (+CHX) or untreated with CHX (-CHX), for groups of mRNAs exhibiting the strongest evidence for 5’UTR unwinding by Ded1 as identified by MaPseq in Guenther et al (17), with P-values of the correlation analyses designated (****, P<0.0001; ***, P<0.001). Top 25% mRNAs with 5’UTR lengths >20nt and >30% MaPseq coverage of the 5’UTR were analyzed, N=222.

The possibility that NCC uORFs would differ from AUG uORFs and exacerbate the impact of impairing Ded1 on mORF translation seemed unlikely in view of previous observations that native NCC uORFs are generally much less effective than AUG uORFs in repressing mORF translation (20). Hence, we investigated further whether the reductions in mORF TEs conferred by *ded1-cs* for NCC uORF-containing transcripts are coupled to their increased 5’UTR translation, in the manner predicted by the Ded1-START model. At odds with this possibility, after binning the NCC uORF-containing mRNAs based on uORF TE changes, it emerged that the mRNAs with the largest increases in uORF TEs exhibit among the smallest, rather than largest, decreases in mORF TEs in *ded1-cs* cells (Fig. 3B, cf. bins 1-3 vs. bins 4-9). Similar results were obtained on examining the *ded1-ts* data (Fig. S8C). Thus, even for the NCC uORF-containing mRNAs, the evidence does not support the model that increased uORF translation is coupled with decreased mORF translation in *ded1* cells. The fact that both mORF and uORF median TEs are reduced the most by *ded1* mutations for mRNAs in the last two bins of Figs. 3B and S8C might be explained, as suggested above, by a decline in 43S PIC attachment on Ded1 inactivation that reduces all initiation events on these mRNAs.

We examined further the tendency for NCC uORF-containing mRNAs to be Ded1-hyperdependent by asking whether the presence of a stable secondary structure just downstream of the uORF start codon exacerbates the effect of *ded1-cs* on mORF translation in the manner expected from the Ded1-START model. To this end, we examined the PARS (parallel analysis of RNA structure) scores assigned to each nucleotide in 2652 native yeast mRNAs, with higher scores indicating more RNA structure (33). Previously, we found marked correlations between elevated PARS scores in 5’UTRs and reductions in mORF TEs conferred by *ded1-cs* and *ded1-ts* mutations (5,10). From the 1047 mRNAs containing one or more NCC uORFs, and the subset of 472 that are Ded1-hyperdependent in *ded1-cs* cells (examined in Fig. 3A, cols. 3 & 6)), we selected the subsets that contain only a single NCC uORF and calculated the cumulative PARS scores for nucleotides in different intervals beginning 16 nt downstream of the uORF start codon, the location where they are most likely to increase uORF translation (14). We divided mRNAs with single NCC uORFs and available PARS data into two groups of high or low cumulative PARS scores for the relevant intervals and compared the effects of *ded1-cs* on translation of the mORFs. Stronger predicted structure 16-45nt downstream of the NCC start codons had little influence on the magnitude of TE reductions conferred by *ded1-cs* (Fig. 3C). The same result was obtained for intervals of 16-30nt and 16-60nt 3’ of the NCC start codons (Figs. S8D-E). These results do not support the model that structure downstream of NCC-uORF start codons generally increases the dependence on Ded1 for mORF translation.

Finally, we examined whether the sets of mRNAs containing NCC uORFs have other features besides uORFs that render them Ded1-hyperdependent. Indeed, these mRNAs, but not those containing AUG uORFs, have longer than average 5’UTR lengths and greater than average PARS scores summed over the entire 5’UTR (Fig. S8F-G, red vs. blue/grey), which could be expected to confer heightened Ded1-dependence independently of the NCC uORFs. We conclude therefore that, unlike long structure-prone 5 ‘UTRs, the presence of AUG- or NCC-initiated uORFs generally does not confer Ded1-hyperdependence in vivo, at odds with the Ded1-START model.

#### uORFs with 3’-proximal Ded1-resolved structures generally do not confer heightened dependence on Ded1

By in vivo DMS mapping of mRNAs (DMS-MaPseq) in *DED1* and *ded1-95* cells at the restrictive temperature, Guenther *et al*. found that Ded1-suppressed ATIS-uORFs tend to contain sequences 10-15nt 3’ of their start codons that are unwound by Ded1, consistent with the model that increased ATIS-uORF translation is driven by downstream structures that remain unwound in *ded1* mutant cells and reduces mORF translation downstream. However, the 185 ATIS-containing mRNAs (harboring 274 ATISs) were not reported to show a greater reduction in mORF TEs compared to all mRNAs (as determined by Ribo-Seq) in *ded1*-*95* versus *DED1* cells (17). Indeed, we made similar observations for the 185 ATIS-containing mRNAs using our own profiling data for the *ded1-cs* and *ded1-ts* mutants, with only the *ded1-ts* (-CHX) data showing a slightly greater than average mORF TE reduction for the ATIS mRNAs (Fig. 3D). Thus, most mRNAs harboring translated uORFs positioned just upstream of Ded1-unwound structures do not have a heightened dependence on Ded1 for mORF translation. We further examined the effect of *ded1* mutations on the TEs of mRNAs with the greatest occurrence of nucleotides dependent on Ded1 for unwinding. Rather than exhibiting increased 5’UTR translation associated with decreased mORF translation in *ded1* cells, we observed significant positive correlations between the TE changes for 5’UTRs and mORFs in our -CHX profiling data for both *ded1-ts* and *ded1-cs* mutants (Fig. 3E). Overall, our Ribo-Seq data do not support the model that suppressing translation of uORFs by unwinding structures downstream of the uORF start codon is the primary mechanism used by Ded1 to stimulate mORF translation.

Guenther *et al*. demonstrated increased ribosome density in the 5’UTR in parallel with reduced mORF TE of the *PSA1* transcript in *ded1*-*95* versus *DED1* cells at the restrictive temperature and showed that mutationally eliminating one of three ATISs at *PSA1* (2^nd^ from the cap, ATIS2 below) diminished the reduction in mORF translation initiation conferred by inactivating the *ded1-95* product, supporting the Ded1-START model (17). Consistent with this, our Ribo-Seq results revealed elevated 5’UTR translation and increased TEs for all three ATISs associated with a reduction in mORF TE in *ded1-ts* vs. *DED1* cells in +CHX data (Fig. S9B). In our -CHX data, however, the 2^nd^ and 3^rd^ ATISs showed no significant TE changes (Fig. S9A), suggesting that the inverse relationship between ATIS2 and mORF TE in *ded1-ts* cells evident in the +CHX data reflects artifactual CHX-induced 5’UTR translation. In the *ded1-cs* mutant at the restrictive temperature (15°C), we found no significant TE increase for ATIS2 associated with reduced mORF TE for *PSA1* in either +CHX or -CHX datasets (Fig. S9C-D), inconsistent with an important role for ATIS2 in mediating Ded1-stimulation of *PSA1* translation by the Ded1-START mechanism.

#### FACS-uORF analysis of native 5’UTRs indicates that uORFs are generally not more inhibitory in *ded1* cells

To provide an orthologous examination of the Ded1-START model we used FACS-uORF, a MPRA system that compares YFP expression from thousands of reporters containing native 5’UTRs harboring uORFs to that of mutant reporters with the uORF start codons mutated to non-functional AAG (20) (Fig. 1B). All reporter genes are expressed from the yeast *ENO2* promoter, shown to be highly stringent and specify transcription start sites consistently across different reporters (20). Fluorescence-activated cell sorting was employed to sort cells into bins according to YFP levels normalized to mCherry levels expressed independently from the same reporter plasmids. The reporter genes in each expression bin were identified by sequencing DNA libraries prepared from the pooled cells. The FACS-uORF library included 21,620 transcript leaders <180 nt long containing at least one uAUG from *S. cerevisiae* (representing 1,524 uORFs) or the closely related species *S. paradoxus* (1,206 uORFs), or conserved *S. cerevisiae* NCC uORFs (29), totaling 3270 uORFs. Start codons were individually mutated in transcript leaders containing multiple uORFs, resulting in 6540 reporters with WT or mutated uORFs (20). We conducted FACS-uORF analysis in parallel in *DED1* and *ded1-cs* strains after shifting log-phase cultures from 30°C to a semi-permissive 23°C (10) for 3 doublings. A semi-permissive growth temperature was required to allow two-three cell doublings to occur following the temperature shift during which pre-existing YFP protein synthesized at the permissive temperature can turnover or be diluted by cell division to reveal reduced YFP expression from Ded1-dependent reporters at the restrictive temperature.

If Ded1 primarily reduces uORF translation to stimulate mORF translation in the manner predicted by the Ded1-START model, then eliminating repressive uORFs should strongly diminish the deleterious effect of the *ded1-cs* mutation on YFP expression (Fig. 4A, row (3), cols. (i)- (ii)). Moreover, the reduction in YFP expression conferred by uORFs should be intensified in the *ded1-cs* mutant, where the uORFs will be more inhibitory, in comparison to *DED1* cells (Fig. 4A(iii), rows 1-3). To test these predictions, we first identified the reporters containing repressive uORFs wherein the mutant uORF reporters lacking a start codon (Mut_uORF) showed consistently and significantly increased YFP expression compared to the corresponding WT_uORF reporters in all three FACS-uORF replicates. In the *DED1* strain, we obtained replicate data for both the WT_uORF and Mut_uORF reporters for 1993 uORFs, 1180 of which were consistently and significantly repressive (CSR). In *ded1-cs* cells, 607 of all 1085 uORFs with data for both WT_uORF and Mut_uORF reporters were CSR, for a total of 1435 CSR uORFs identified in one or the other strain (Fig. S10A). Combined, we obtained replicate YFP expression data for 510 CSR uORFs in both *DED1* and *ded1-cs* cells, allowing us to evaluate whether uORF inhibition is increased when Ded1 function is impaired for a large set of repressive uORFs (Fig. S10B).

**Figure 4.**
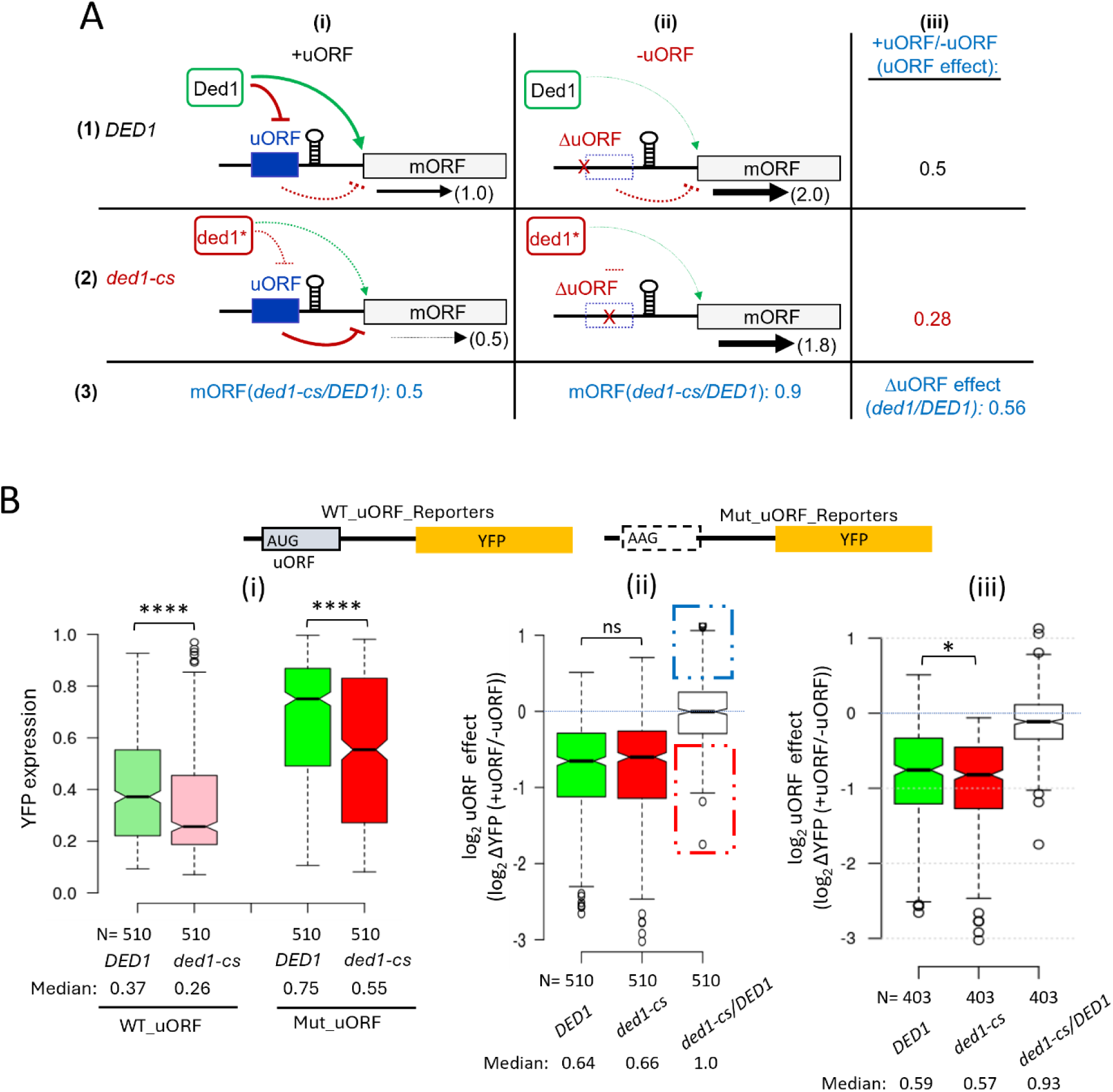
MPRA of the FACS-uORF library indicates that most repressive uORFs are not more inhibitory in *ded1-cs* vs. *DED1* cells. **(A)** Schema summarizing predictions of the Ded1-START model for mRNAs containing repressive uORFs. (i) (+uORF), *DED1:* WT Ded1 stimulates translation of the mORF indirectly by damping inhibition by the uORF whose recognition by scanning PICs is enforced by a stem-loop downstream of the uORF start codons that can be unwound by Ded1. The amount of mORF translation is assigned a value of 1.0; (ii) (-uORF), *DED1*: Removal of the uORF start codon increases the proportion of scanning PICs that reach the mORF and increases mORF translation 2-fold, yielding a +uORF/-uORF expression ratio (uORF effect) of 0.5 as listed in panel (iii); (i) (+uORF), *ded1-cs:* Impairment of Ded1 increases the proportion of scanning PICs that pause at the stem-loop and translate the uORF versus the mORF, intensifying repression by the uORF to decrease mORF translation by a factor of 0.5 compared to *DED1* cells, as indicated on row 3; (ii) -uORF, *ded1-cs*: In the absence of the uORF, impairing Ded1 produces only a small 0.9-fold reduction in mORF translation (shown on row 3) owing to the increased barrier to scanning PICs posed by the unwound stem-loop. Because the uORF is more inhibitory in *ded1-cs* cells, the derepressing effect of its removal is greater than in *DED1* cells, with a uORF effect of 0.28. The resulting *ded1-cs/DED1* ratio of ORF effects is 0.56 (row 3). **(B) (i)** Notched box plot of normalized YFP expression for reporters containing the native uORFs (WT_uORF) or with AAG substitutions of the uORF start codons (Mut_uORF), as depicted in the schematics above, that were detected by MPRA screening of the FACS-uORF library in *DED1* strain NSY4 or *ded1-cs* mutant NSY5 cells cultured in SC-L-U medium for ∼3 doublings at 23°C for the 510 uORFs depicted in Fig. S10B that were judged to be CSR in either *ded1-cs* or *DED1* cells and also contain replicate reporter data for both Mut_uORF and matched WT_uORF reporters in both *ded1-cs* and *DED1* cells. Median YFP expression is given below the genotype in each column. **(ii)** Notched box plot of log_2_ uORF effects (log2 ratios of YFP expression for the WT_uORF vs. corresponding Mut_uORF reporters) measured in *DED1* or *ded1-cs* cells and the log_2_ of *ded1-cs/DED1* ratios of uORF effects (col. 3) for the 510 uORFs analyzed in (i). The un-logged median values are given below the genotypes in each column. **(iii)** Analysis identical to that in panel (ii) but for the 403 uORFs belonging to the group of 607 uORFs judged to be CSR in *ded1-cs* cells, with replicate reporter data for both the Mut_uORF and matched WT_uORF reporters in both *ded1-cs* and *DED1* cells, as depicted in Fig. S10C. Results of Mann-Whitney U tests are summarized as: ****, P<0.0001; *, P<0.05; n.s., not significant.

Analyzing this group of 510 CSR uORFs revealed that the median YFP expression decreased 30% in *ded1-cs* mutant vs. *DED1* cells, (p < 0.0001; Fig. 4B(i), cols. 1-2), in keeping with the established role of Ded1 in promoting translation of most yeast mRNAs. The corresponding group of 510 Mut_uORF reporters were expressed ∼2-fold higher than WT_uORF reporters in *DED1* cells (0.75/0.37, Fig. 4B(i), cols. 3 vs. 1), as expected for the elimination of repressive uORFs. Importantly, the Mut_uORF reporters were also expressed 27% lower in *ded1-cs* vs. *DED1* cells, (p < 0.0001; Fig. 4B(i), cols. 3-4), which is similar to the 30% decrease in WT uORF reporter expression on Ded1 impairment shown in cols. 1-2. This last comparison was the first indication that the effect of impairing Ded1 in reducing reporter expression is generally unaffected by the presence of the uORFs assayed by FACS-uORF.

To examine this inference more rigorously, we compared the magnitudes of YFP repression for each uORF (“the uORF effect”) as the ratio of expression in the presence vs. absence of each uORF determined individually in *DED1* and *ded1-cs* strains (as explained in Fig. 4A(iii)) for all 510 repressive uORFs. The median uORF effects (+uORF/-uORF) did not differ significantly between the two strains (0.64 vs 0.66; Fig. 4B(ii), cols. 1-2). Moreover, the ratios of the uORF effects calculated for each of the 510 CSR uORFs in *ded1-cs* vs. *DED1* cells have a median value of 1.00 (Fig. 4B(ii), col. 3), indicating that most uORFs are similarly repressive in *DED1* and *ded1-cs* cells. Overall, these results indicate that the repressive effects of the majority of the 510 CSR uORFs on translation of downstream mORFs are not mitigated by Ded1, implying in turn that the Ded1-START model may apply to only a small subset of all repressive uORFs.

We considered the possibility that the presence of multiple uORFs in the 5’UTRs of reporters containing more than one repressive uORF could diminish the consequences of impairing Ded1 on the inhibitory effect of the particular uORF that was mutated. To address this, we repeated the analysis in Fig. 4B(ii) for the subset of 374 uORFs found in reporters with no other AUG-initiated uORFs in the 5’UTRs. The results in Fig. S10D again showed essentially no difference in the inhibitory effects of the uORFs between *DED1* and *ded1-cs* cells, with a median ratio of uORF effects in *ded1-cs* vs. *DED1* cells close to unity (0.98).

It could be argued that the uORFs most likely to conform to the Ded1-START model will be significantly repressive only in the *ded1-cs* mutant, where their proximal structures will remain unwound to enhance uORF translation. Accordingly, we repeated the analysis described in Fig. 4B(ii) for all 607 uORFs found to be CSR in the *ded1-cs* strain and for which data were also obtained in *DED1* cells regardless of whether the uORF was significantly repressive in the *DED1* strain (Fig. S10C). As shown in Fig. 4B(iii), these 403 uORFs were slightly more inhibitory in the *ded1-cs* vs. *DED1* strain, with a median ratio of uORF effects in *ded1-*cs/*DED1* cells of 0.93, suggesting that a fraction of these uORFs do conform to the Ded1-START model. Consistent with this, a subset of the 510 CSR uORFs analyzed above exhibit ratios of uORF effects in *ded1-cs* vs. *DED1* cells well below unity (Fig. 4B(ii), col. 3, dotted red box). Hence, we selected for further examination the subset of uORFs with calculated uORF effects >1.33-fold smaller in *ded1-cs* vs *DED1* cells (those in the dotted red box in Fig. 4B(ii)), which exhibit the greatest Ded1-suppression of uORF inhibitory effects among all 510 uORFs. We similarly identified a subset of uORFs whose inhibitory effects are >1.33-fold larger in *ded1* vs. *DED1* cells (in the dotted blue box of Fig. 4B(ii)), which appear to be the most dependent on Ded1 for the ability to repress translation of the downstream mORFs. This latter behavior is not predicted by the Ded1-START model, as it implies that Ded1 enhances rather than suppresses uORF translation, possibly by resolving a structure that includes the uORF initiation region. Among the 46 presumptive Ded1-suppressed uORFs falling within the red dotted box in Fig. 4B(ii), 16 reside in genes we identified as being Ded1-hyperdependent in our Ribo-Seq data of the *ded1-cs* mutant described above and that also contain uORFs found in both *S. cerevisiae* vs. *S. paradoxus* 5’UTRs where they could potentially mediate the reduced translational efficiencies of the corresponding transcripts at reduced Ded1 function.

To pursue this last possibility, we attempted to confirm Ded1-suppression of the repressive effects of 10 of the aforementioned 16 uORFs showing the strongest evidence for the Ded1-START model in our FACS-uORF data by constructing the WT_uORF and Mut_uORF reporters, designed identically as in the FACS-uORF library under the *ENO2* promoter (WT and Mut reporters 1-10 in Table S1). We similarly constructed WT_uORF and Mut_uORF reporters for 4 uORFs whose inhibitory effects appeared to be enhanced rather than suppressed by Ded1 in the FACS-uORF analysis (WT and Mut reporters 11-14 in Table S1). Expression of all 14 pairs of WT_uORF and Mut_uORF reporters was measured individually in transformants of the *ded1-cs* and *DED1* strains cultured under the same conditions employed in the FACS-uORF experiments above.

Assaying both classes of WT_uORF reporters revealed lower median expression compared to the corresponding Mut_uORF reporters in both *DED1* and *ded1-cs* cells, as expected for repressive uORFs (Figs. 5A(i)-(ii), cf. cols. 1-2 & 3-4). In addition, the median expression of both WT and mutant uORF reporters was lower in *ded1-cs* compared to *DED1* cells for the 10 reporters with presumptive Ded1-suppressed uORFs (Figs. 5A(i), cf. cols. 1 & 3 and 2 & 4). The fact that removing the repressive uORFs generally did not eliminate diminished reporter expression in *ded1-cs* cells departs from the Ded1-START model and implies that Ded1 overcomes one or more inhibitory elements in the 5’UTRs besides repressive uORFs. The same conclusion applies to the four reporters with presumptive Ded1-enhanced inhibitory uORFs, whose median expression was again lower in *ded1-cs* vs. *DED1* cells for the mutant uORF reporters (Figs. 5A(ii), cf. cols. 2 & 4). The median expression of the WT uORF reporters was also lower in *ded1-cs* vs. *DED1* cells but the difference is not statistically significant (Figs. 5A(ii), cf. cols. 1 & 3), possibly owing to the small number (four) of reporters under comparison.

**Figure 5.**
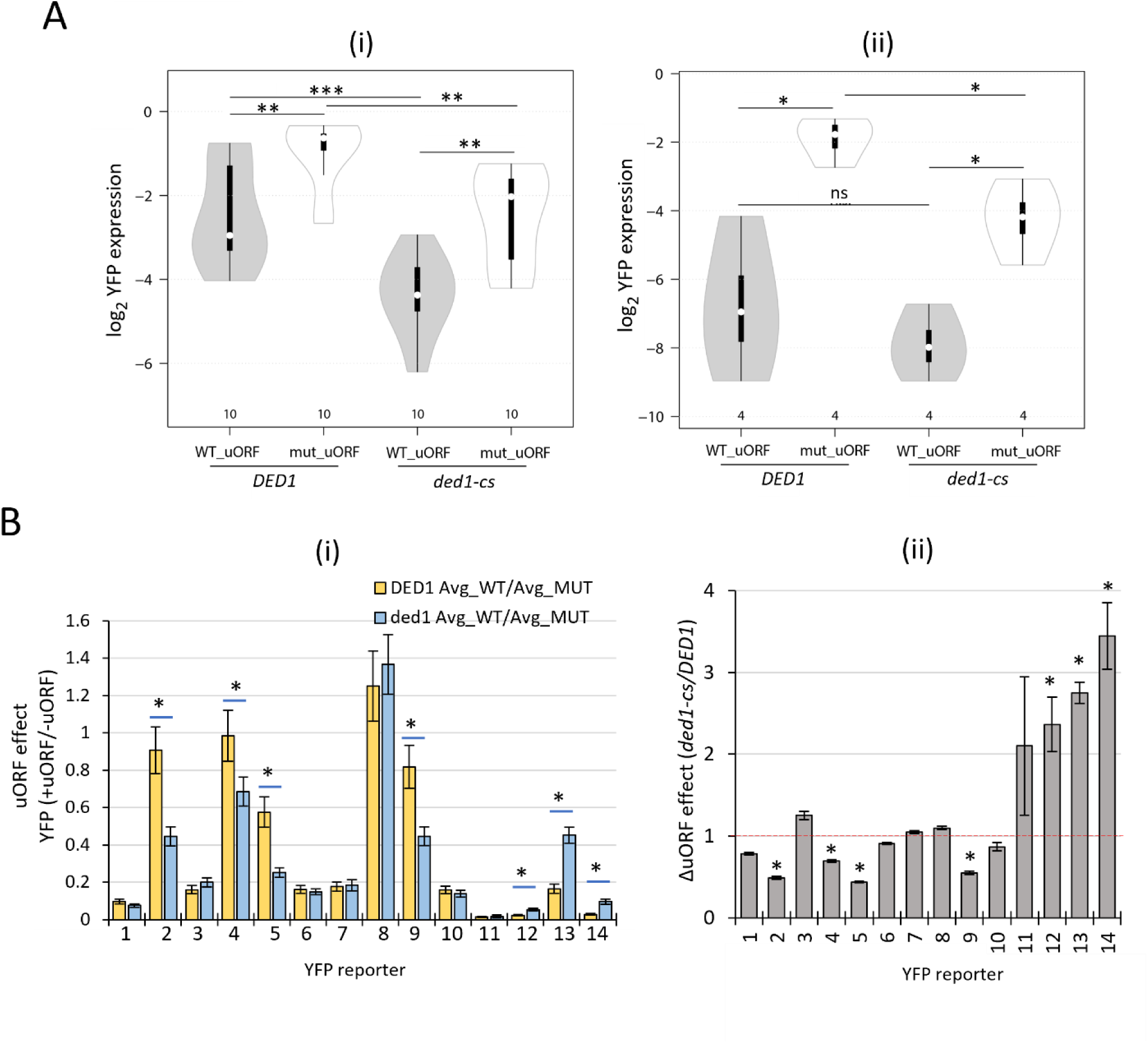
Assays of individual WT_uORF and Mut_uORF reporters for genes exhibiting evidence from the FACS-uORF library screening for Ded1 modulation of uORF effects. **(A)-(i)** Violin plots of log_2_ YFP expression for 10 uORF reporters whose MPRA results adhere to the predictions of the Ded1-START model in showing enhanced uORF repression of YFP expression in *ded1-cs* vs. *DED1* cells. Three independent transformants of the WT_uORF and corresponding Mut_uORF *YFP* reporters in strains NSY4 or NSY5 were cultured in SC-L-U medium for ∼3 doublings at 23°C and the YFP/mCherry fluorescence ratio was measured in whole cells. Results of Mann-Whitney U tests are indicated as follows: ***, P<0.001; **, P<0.01; *, P<0.05; n.s., P>0.05. **A-(ii)** Analysis conducted exactly as in (i) for 4 uORF reporters, whose MPRA results depart from predictions of the Ded1-START model in showing diminished uORF repression of YFP expression in *ded1-cs* vs. *DED1* cells, summarizing data from 3 biological replicates for each reporter/strain combination. **(B)-(i)** Data from (A) were analyzed by plotting ratios of YFP expression for the +uORF vs. -uORF reporters (uORF effects), with error bars depicting the S.E.M.s calculated for the ratios determined in *DED1* (cyan) vs. *ded1-cs* (gold) transformants with results from student’s t-tests indicated (*, P<0.05) only for those comparisons where the differences between mean values from *DED1* vs. *ded1* cells are statistically significant. **(B)-(ii)** ratios of the mean uORF effects in *ded1-cs* vs. *DED1* cells from (B)-(i) for the indicated uORF reporters. Results from student’s t-tests are indicated as in (B)-(i).

We next examined the reporters individually to determine which ones show evidence of Ded1 regulation of uORF effects in the manner predicted by the FACS-uORF data. In the *ded1-cs* mutant, 9 of the 10 reporters containing presumptive Ded1-suppressed ORFs (with the exclusion of reporter 8) showed significantly lower expression in the presence vs. absence of the uORF, leading to +uORF/-uORF expression ratios (uORF effects) of <1, in the manner expected for inhibitory uORFs (Fig. 5B(i), cyan bars). In *DED1* cells, by contrast, only 6 of 10 reporters showed uORF effects <1, as removing the uORFs for reporters 2, 4, 8, and 9 did not confer a significant difference in expression (Fig. 5B(i), gold bars). As a result, reporters 2, 4, and 9 showed ratios of uORF effects in *ded1-cs* vs. *DED1* cells of <1 (Fig. 5B(ii)). This was also true of reporter 5 even though its uORF is inhibitory in *DED1* cells, because the uORF is significantly more inhibitory in the *ded1-cs* strain (Fig. 5B(i), cyan vs gold). Thus, the reporter data confirm our FACS-uORF results indicating Ded1-mediated suppression of uORFs 2, 4, 5, and 9, that reside, respectively, at genes *YDR072C, YDR186C, YJR054W*, and *YGR010W*. Our results also confirmed the FACS-uORF data for three of the four Ded1-enhanced uORFs, showing larger uORF effects in the *ded1-cs* vs. *DED1* strain (Fig. 5B(i), cyan vs. gold, reporters 12-14). Together, these results substantially validate our FACS-uORF analysis and confirm that uORF repression at certain genes can be either increased or decreased by reducing Ded1 function.

The Ded1-START model posits that Ded1 suppresses inhibitory uORFs by unwinding secondary structures located downstream of their start codons that enhance initiation at the uORFs and prevent leaky-scanning to the downstream mORFs. To examine whether Ded1-resolved structures are located downstream of the uORFs in the aforementioned reporters 2, 4, 5, and 9, whose expression conforms with the Ded1-START model, we interrogated the DMS-MaPseq data obtained for *ded1* mutant and *DED1* cells utilized by Guenther et al. to identify the Ded1-suppressed ATISs mentioned above. For the uORFs in reporters 5 and 9, the DMS-MaPseq data coverage was inadequate to evaluate the presence of a Ded1-resolved structure downstream of their start sites. Although the DMS-MaPseq coverage was sufficient for reporters 2 and 4, a Ded1-resolved structure was not indicated within 8-30nt downstream of the uORF start codons. Thus, we were unable to ascribe the increased repressive effects in *ded1* cells of the uORFs in reporters 2, 4, 5, and 9 to a Ded1-resolved structure positioned downstream of the uORF start codons.

We next interrogated the published DMS-MaPseq data to identify other Ded1-suppressed uORFs situated just upstream of Ded1-resolved structures. We found seven CSR uORFs identified in the *ded1*-cs strain whose WT 5’ UTRs drive lower YFP expression in *ded1-cs* vs. *DED1* cells and show evidence for Ded1-resolved structures downstream of the uORF start codons (Fig. S11A(i)-(vii)). Four different YFP reporters were generated for each uORF containing the complete 5’UTR sequence of the corresponding gene and harboring either (i) WT uORF and WT downstream structure (WT-WT), (ii) Mut uORF and WT structure (Mut-WT), (iii) WT uORF and mutated structure (WT-Mut), and (iv) Mut uORF and mutated structure (Mut-Mut), substituting residues involved in Ded1-suppressed structures in the 7-30 nt intervals downstream of the uORF start codons with CAA nucleotides to destabilize the structures (6), as summarized schematically in Figs. 6A and S11B). All seven WT-WT reporters showed significantly lower YFP expression ratios in *ded1-cs* vs. *DED1* cells, as expected from the FACS-uORF data (Fig. 6B). However, only three of the seven uORFs, those present at *YIL09C, YBL032W, YDR481C*, conformed to all predictions of the Ded1-START model. Thus, these uORFs were more repressive in *ded1-cs* cells when the downstream structure was intact (Fig. 6C, cols. a vs b, reporters 1-3) and they were relatively less repressive in *ded1-cs* cells in the reporters with disrupted downstream structures (Fig. 6C, cols b vs. d, reporters 1-3). As a result, the ratios of uORF effects in *ded1-cs* vs. *DED1* cells had significantly smaller values in the presence vs. absence of the structures (Fig. 6D, cols. 1 vs. 2, 3 vs. 4, and 5 vs. 6). We conclude that these three uORFs are strong candidates for exerting Ded1-suppressed translational repression of the downstream mORFs according to the Ded1-START model.

**Figure 6.**
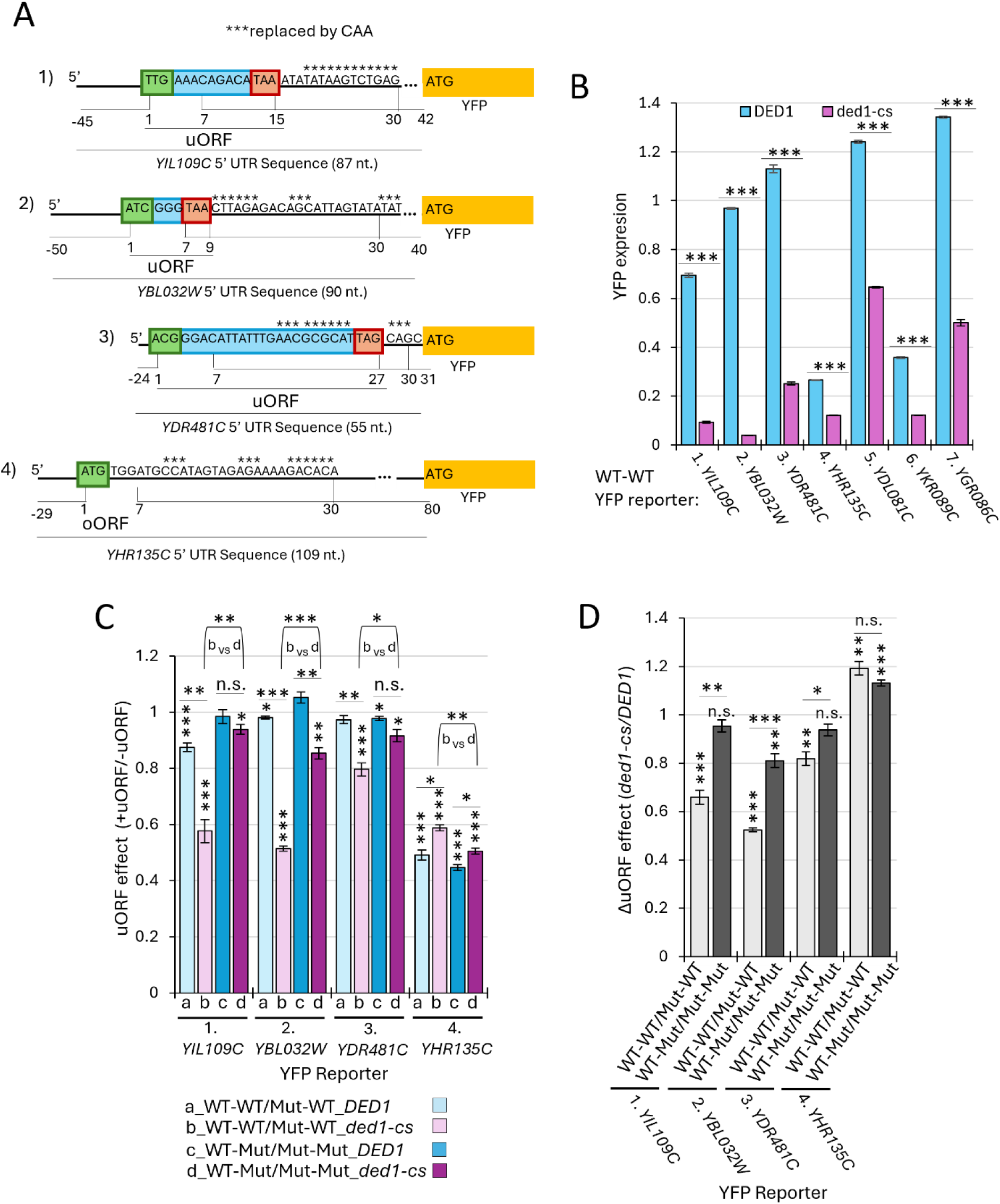
Reporter analysis identifies three repressive uORFs conforming to the Ded1-START model. **(A)** Schematics depicting the 5’ UTRs (solid line from 5’ to ATG of YFP), uORFs (depicting the start codon, coding sequence and stop codon in green, cyan, and pink boxes, respectively) that were eliminated in the Mut-WT and Mut-Mut reporters by replacing the start codons with AAG, and downstream sequences with evidence from DMS-MapSeq for Ded1-resolved structures (labeled with asterisks) that were replaced by CAA repeats to destabilize the structures in the WT-Mut and Mut-Mut reporters. **(B)** YFP expression was determined for three independent transformants of strains NSY4 (*DED1)* or NSY5 (*ded1-cs)* harboring the indicated WT-WT reporters as described in panel A, with results of student’s t-tests indicated as: ***, P<0.001. **(C)** YFP expression was determined for three independent transformants of strains NSY4 (*DED1*) or NSY5 (*ded1-cs*) for the panel of four WT-WT, Mut-WT, WT-Mut, and Mut-Mut reporters for each of the four repressive uORFs depicted in (A) and the ratios of YFP expression in the presence vs. absence of the uORF (uORF effects) were determined for reporters containing either WT downstream structures (WT-WT/Mut-WT) or those with CAA-substituted structures (WT-Mut/Mut-Mut) in both *DED1* and *ded1-cs* cells and plotted as indicated in the key. The results of t-tests comparing mean YFP expression in the presence vs. absence of the uORF are indicated as: ***, P<0.001; **, P<0.01; *, P<0.05; n.s., P>0.05; as are results of t-tests comparing the uORF effects between columns a vs. b, c vs. d, or b vs. d for each reporter. **(D)** Ratios of uORF effects determined in (C) between *ded1-cs* and *DED1* cells for the indicated reporters. Results of t-tests comparing uORF effects between *ded1-cs* and *DED1* cells as determined in panel C (cols a vs. b or c vs. d)) are replotted above each bar; as are results of t-tests comparing the ΔuORF effects between adjacent columns plotted here.

The reporter data for *YHR135C,* by contrast, depart from the Ded1-START model in revealing a strongly repressive uORF that is somewhat less, rather than more, inhibitory in *ded1-cs* vs. *DED1* cells, and is slightly more, rather than less, inhibitory in the absence of the downstream structure (Fig. 6C, cols. a vs b and b vs d, reporter 4), all suggesting that repression by this uORF is exerted independently of both Ded1 and the downstream structure. Interestingly, this last uORF was classified as an ATIS by Guenther et al. and it also showed a significantly greater uORF effect in *ded1-cs* vs. *DED1* cells in our FACS-uORF analysis in the manner expected for a Ded1-suppressed uORF. It is unclear why our reporter analysis did not confirm dependence on Ded1 for its repressive effect on mORF translation. As described in detail in Fig. S11B-D, the reporter data we obtained for the three remaining uORFs we interrogated, at genes *YDL081C, YGR086C,* and *YKR089C,* also depart from one or more predictions of the Ded1-START model, preventing us from confirming their adherence to the model.

In summary, the results of our FACS-uORF analysis in *ded1-cs* vs. *DED1* cells suggests that the inhibitory effects of most uORFs are not increased in the *ded1-cs* mutant in the manner predicted by the Ded1-START model among a large group of 510 uORFs that could be fully interrogated for compliance with the predictions of the model. By analyzing individual reporters for a group of 7 uORFs whose FACS-uORF data conform to the Ded1-START model and also show evidence for a proximal Ded1-resolved structure, we obtained data confirming that the uORFs at three genes, *YIL09C, YBL032W,* and *YDR481C,* do repress translation of the downstream main ORFs in a manner exacerbated by downstream structures and diminished by Ded1 function. Overall, it appears that only a small fraction of repressive uORFs are regulated by Ded1 according to the Ded1-START model.

### The presence of longer, more structured 5’UTRs is associated with greater reductions in YFP reporter expression in *ded1-cs* cells

We asked next whether the effects of the *ded1-cs* mutation on the complete set of YFP reporters harboring WT 5’UTRs are consistent with our Ribo-Seq data of the *ded1-cs* mutant. To this end, we analyzed the subset of all 5185 reporters with WT 5’UTRs from *S. cerevisiae* with data in both *ded1-cs* and *DED1* cells (numbering 2656), of which 913 contain one or more uORFs that were mutagenized in the Mut_uORF library, as discussed above. Comparing the changes in YFP reporter expression to the changes in TE calculated in *ded1-cs* vs. *DED1* cells using our (+CHX) Ribo-Seq data reported previously (5) revealed a highly significant correlation between the two data sets, with a Spearman correlation coefficient (ρ) of 0.31 and P-value of <0.00001 (Fig. 7A). Essentially identical correlation results were obtained using the (-CHX) Ribo-Seq data we obtained here (ρ = 0.30, P <0.00001). We then identified the two sets of mRNAs showing the largest reductions or increases in relative TE conferred by *ded1-cs* in the Ribo-Seq experiments (>1.41-fold change, FDR<0.1), dubbed TE_down and TE_up, respectively. We found that the 671 YFP reporters representing the TE_dn group of genes display a reduction in median YFP expression in *ded1-cs* vs. *DED1* cells greater than that seen for all 2009 genes with YFP reporter data, which was not observed for the 600 reporters representing the TE_up genes (Fig. 7B). Thus, we observed a significant tendency for reporters to show greater than average reductions in YFP expression in the *ded1-cs* mutant if they correspond to mRNAs judged to be Ded1-hyperdependent by Ribo-Seq analysis. These correlations are especially notable considering that the YFP reporter library does not include the ∼10% of all yeast genes with 5’UTRs longer than 180 nt, which tend to be the most dependent on Ded1 for efficient translation both in vivo (5) and in the yeast reconstituted system (18). In addition, the FACS-uORF measurements were conducted at the semi-permissive growth temperature of 23°C to allow for YFP synthesis in the *ded1-cs* cells whereas the Ribo-Seq experiments were carried out at the non-permissive temperature of 15°C where the catalytic activity of the *ded1-cs* product and bulk translation is more severely impaired and 5’UTR structures are relatively more stable.

**Figure 7.**
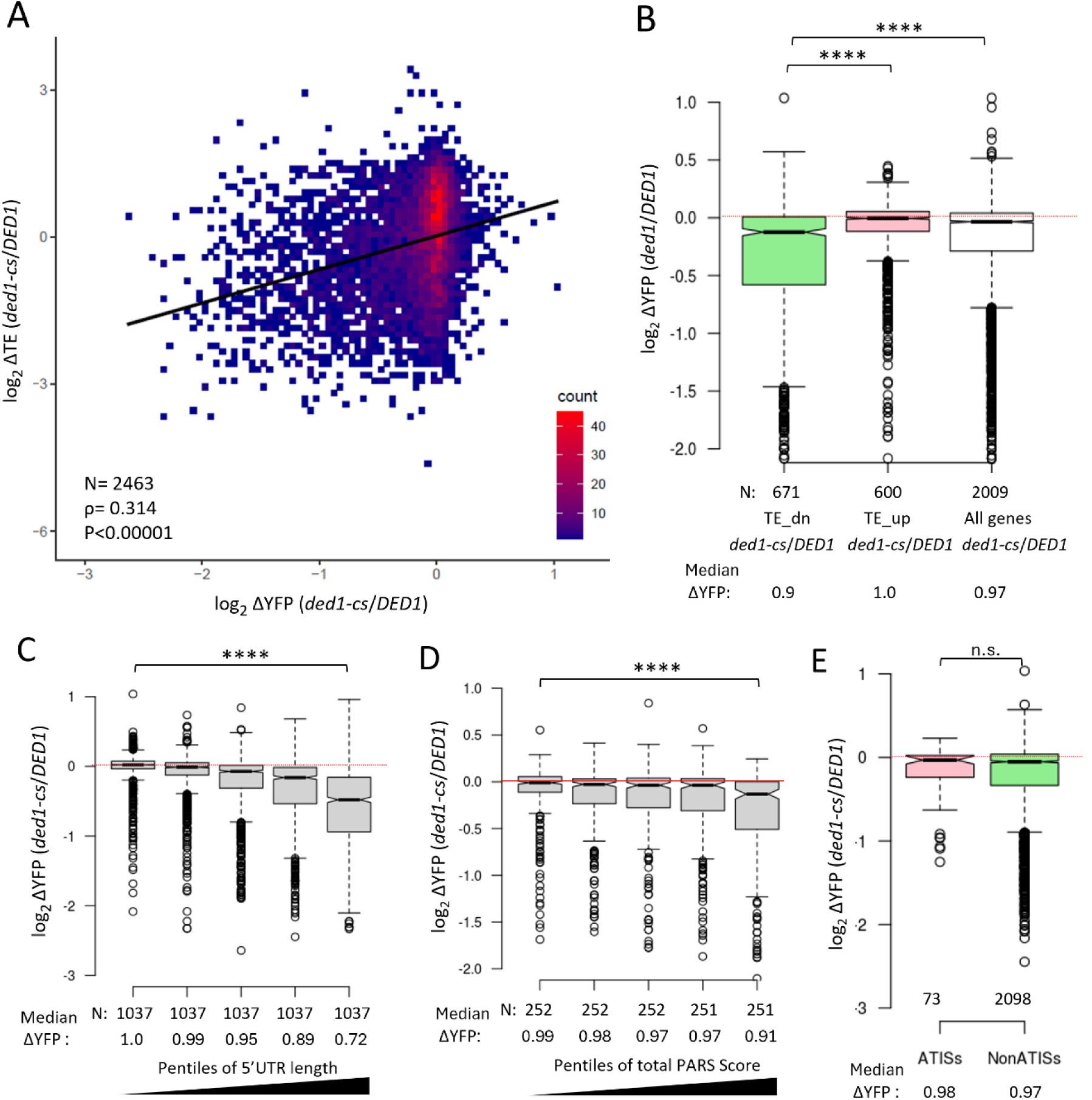
Correlations between effects of *ded1-cs* on expression of WT YFP reporters and TEs of the native genes, 5’UTR lengths, predicted 5’UTR structures and presence of ATISs. **(A)** Intensity scatterplot of log_2_ changes in TE between *ded1-cs* and *DED1* cells cultured at 15°C previously determined by Ribo-Seq (5) and the log_2_ changes in YFP between *ded1-cs* and *DED1* cells determined at 23°C by MPRA of all WT YFP reporters in the FACS-uORF library for the 2463 genes with data available for both measurements, with the Spearman ρ and P-value for the correlation indicted. Colors indicate numbers of gene counts with the same x and y coordinates as indicated in the color key. The greater range of TE changes seen on the y-axis for the sizeable group of YFP reporters showing little change in YFP expression (centered around 0 on the x-axis) might reflect the greater impairment of the *ded1-cs* product at 15°C (in Ribo-Seq) vs. 23°C (in MPRA). **(B)** Notched boxplot of the log_2_ changes in reporter expression between *ded1-cs* and *DED1* cells for the subset of 671 and 600 WT YFP reporters whose corresponding native genes showed significant reductions or increases in TE (>1.41-fold change, FDR<0.1) between *ded1-cs* and *DED1* cells in Ribo-Seq, dubbed TE_down and TE_up, respectively. **(C)** Notched boxplot of the log_2_ changes in reporter expression between *ded1-cs* and *DED1* cells for the indicated pentiles of 5185 WT YFP reporters binned according to the 5’UTR lengths of the reporter constructs for all 5185 reporters analyzed by MPRA. **(D)** Similar analysis as in (C) but binned according to the total PARS scores summed across all 5’UTR nucleotides for the subset of 1258 WT YFP reporters with available PARS data. 5’UTR PARS scores were computed for the single 5’UTR length for each gene referenced in Kertesz et al. (2010) and may not match the 5’UTR length that we determined previously and was employed for YFP reporter design (20); nor does it consider differences in total PARS scores for reporters of differing 5’UTR lengths constructed for the same gene. **(E)** Similar analysis as in (C) but binned according to presence of ATISs for the 2171 YFP reporters containing WT *S. cerevisiae* 5’UTRs for which only a single reporter represents each gene in the FACS-uORF library. Results of Mann-Whitney U tests are indicated (****, P<0.0001; n.s., P>0.05).

**Figure 8.**
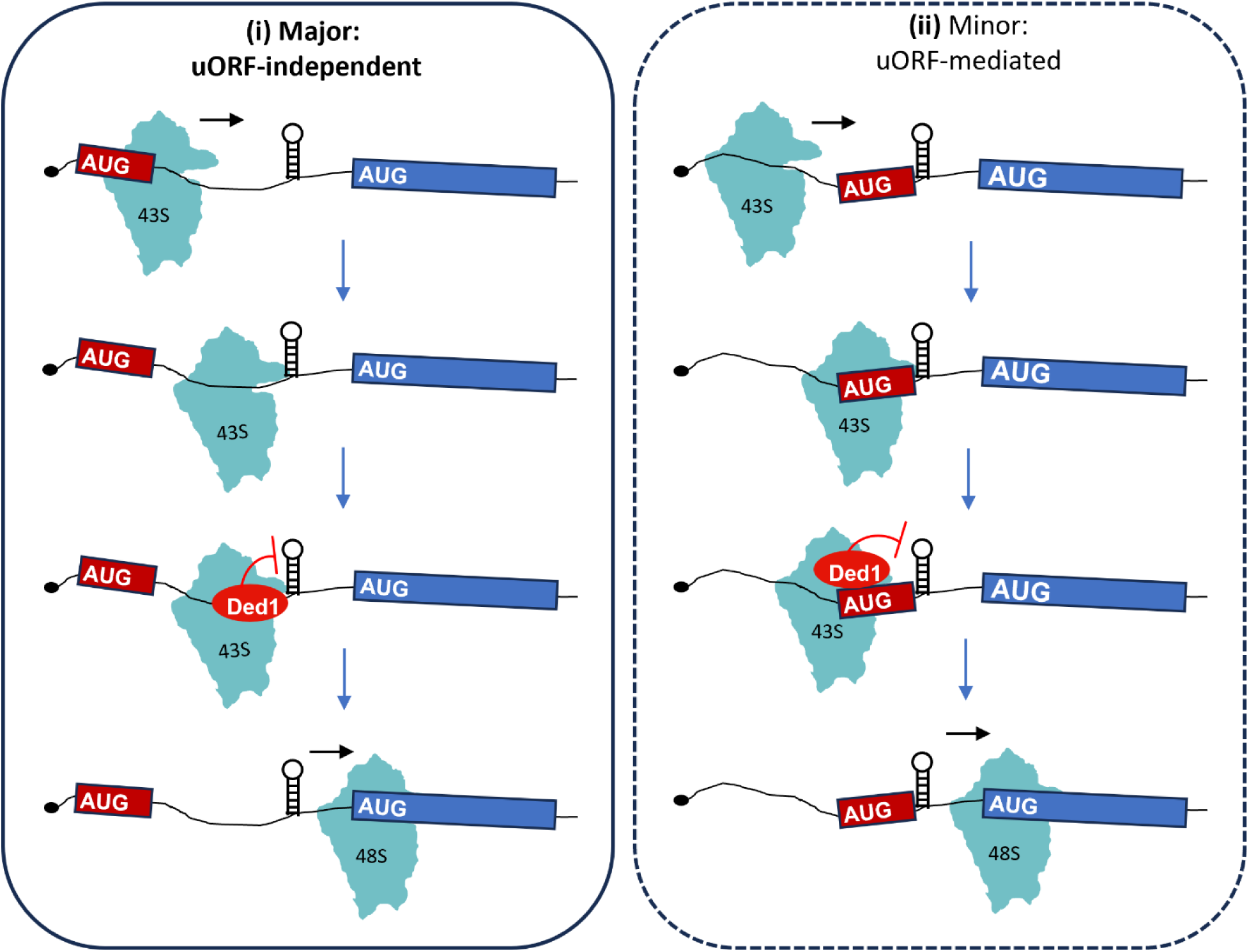
Model: Ded1 generally stimulates initiation on mRNAs burdened with secondary structure without modulating translational repression by uORFs. **(i)** For the great majority of mRNAs, Ded1 stimulates the traversal of 5’UTRs by scanning 43S PICs and subsequent initiation at the mORF start codon (blue box with AUG) by resolving secondary structures in the 5’UTRs regardless of the presence or absence of translated uORFs (red box with AUG). While not depicted, the same applies to uORFs with NCC initiation sites. **(ii)** For a small number of mRNAs, Ded1 stimulates traversal of 5’UTRs by suppressing initiation at a repressive uORF by resolving a proximal structure that pauses the 43S PIC and increases its probability of initiating at the uORF start codon rather than scanning downstream to the mORF AUG codon, as proposed by the Ded1-START model.

Our Ribo-Seq analysis of the *ded1-cs* mutant indicated that Ded1-hyperdependent mRNAs have 5’UTRs that are longer and more structure-prone than the average mRNA (10). Consistent with these findings, we found that the reduction in YFP reporter expression conferred by *ded1-cs* generally increased with 5’UTR length and potential for secondary structure. The set of all 5185 reporters was divided among five pentiles of increasing 5’UTR length, with median lengths of 21 nt for pentile 1 and 112 nt for pentile 5. As shown in Fig. 7C, the *ded1-cs* mutation conferred progressively greater reductions in YFP expression across pentiles 2-5 compared to the first pentile, ranging from 0.99 to 0.72. We then sorted the reporters differently into bins of increasing total PARS scores for the entire 5’UTR sequences of reporter genes with available PARS data (33). As shown in Fig. 7D, the pentile of highest total PARS scores showed a greater reduction in YFP expression conferred by *ded1-cs* compared to the other four pentiles. Because the total PARS score reflects both 5’UTR structure and length, we conducted a separate analysis of PARS scores in a sliding 30 nt window across the 5’UTR to identify the interval with the maximum summed PARS score (Max30), which should reflect the presence of local RNA structures independently of 5’UTR length. As shown in Fig. S12A, we observed a significant tendency for mRNAs with the highest Max30 PARS scores to show the greatest reductions in YFP reporter expression in *ded1-cs* vs. *DED1* cells. As a complementary approach, we used RNAfold to estimate the change in free energy (ΔΔG) required to unfold the entire 5’UTR (35) and sorted the reporters into pentiles of increasing ΔΔG. As shown in Fig. S12B, increasing estimated 5’UTR unfolding energy is associated with greater reductions in YFP expression conferred by *ded1-cs* across all five pentiles.

Studies on the mammalian ortholog Ddx3 provide evidence that it binds preferentially to 5’UTR sequences capable of forming G-quadruplexes (hereafter “G-quads”), structures produced by stacking of G-quartets comprised of four hydrogen-bonded guanines (36,37). Evidence indicates that Ddx3 can destabilize particular G-quads in vitro (38) and it unwinds such structures in stimulating the translation of particular Ddx3-target mRNAs in cells (37,38). Ded1 was also shown to bind and unwind a particular G-quad present in a native yeast mRNA (39). Interestingly, we found that YFP reporters identified previously as harboring a predicted G-quad structure (35) showed larger median reductions in YFP expression in *ded1-cs* vs. *DED1* cells compared to all reporters lacking them (Fig. S12C). Together, these results indicate that longer, more structured 5’UTRs, including those capable of forming Gquad structures, are associated with heightened dependence on Ded1 for *YFP* reporter expression in yeast cells.

Sorting the 5185 WT reporters according to number of upstream AUGs (uAUGs) in the 5’UTRs revealed greater reductions in YFP expression in *ded1-cs* vs. *DED1* cells for reporters with one or more uAUGs compared to those lacking uAUGs (Fig. S12D). Except for the fact that the reductions were similar in magnitude between one and several AUGs, this finding is ostensibly consistent with the Ded1-START model. However, we found that the presence of greater numbers of uAUGs is associated with increasing 5’UTR lengths for the WT YFP reporters in the library (Fig. S12E), making it unclear whether the presence of uAUGs or 5’UTR length is the more important determinant of Ded1 dependence. To help resolve this issue, we examined the effect of 5’UTR length and PARS scores for the majority class of 4272 reporters lacking uAUGs. As shown in Fig. S12F-G, the reduction in YFP expression conferred by *ded1* once again scaled with increasing 5’ UTR length and total PARS score in 5’UTRs. These findings suggest that Ded1 generally stimulates translation of mRNAs with long or structured 5’UTRs even when they lack AUG-initiated uORFs.

To extend our analysis of whether uORFs confer a greater dependence on Ded1 for efficient translation, we determined the effect of the *ded1-cs* mutation on expression of the set of 73 WT YFP reporters containing one or more of the 255 high-confidence Ded1-suppressed ATISs identified by Guenther et al. As shown in Fig. 7E, we found no significant difference in the median change in reporter expression conferred by the *ded1-cs* mutation between the 73 ATIS-containing reporters and the 2098 reporters lacking ATISs found among the subset of 2171 reporters with *S. cerevisiae* WT 5’UTRs and only a single 5’UTR representing each gene in the library. Thus, the presence of an ATIS generally did not confer heightened dependence on Ded1 for efficient translation of the corresponding YFP reporter mRNAs, at odds with the Ded1-START model.

## DISCUSSION

It has been proposed that Ded1 enhances translation of protein coding genes primarily by unwinding RNA structures to prevent premature initiation at upstream ORFs that would preclude translation of the main ORFs (mORFs) downstream (17). We used a combination of Ribo-Seq, MPRA, and genetic analysis of reporter genes to examine the applicability of this Ded1-START mechanism in vivo for the large cohort of mRNAs showing heightened dependence on Ded1 for efficient translation. Our findings indicate that Ded1 generally stimulates translation of mORFs by overcoming the inhibitory effects of long, structured 5’UTRs independently of its influence on uORF translation, and that the Ded1-START mechanism applies to only a small subset of Ded1-hyperdependent transcripts.

We conducted new Ribo-Seq experiments on *ded1* mutants omitting treatment of cells with CHX to avoid artifactually increasing 80S occupancies in 5’UTRs (19) and examined whether decreased mORF translation in *ded1* mutants was associated with elevated 5’UTR translation, as predicted by the Ded1-START model. This inverse correlation was not observed in Ribo-seq data obtained with or without CHX pretreatment. Moreover, the presence of translated AUG-initiated uORFs in *DED1* cells is not significantly associated with Ded1-hyperdependence for mORF translation determined by Ribo-Seq. These conclusions also apply to mRNAs identified by Guenther et al. containing uORFs, dubbed ATISs, whose translation appeared to be increased in a *ded1* mutant and were enriched for RNA structures downstream of the uORFs that appear to require Ded1 for unwinding in vivo. The group of ATIS-containing mRNAs does not show a greater than average reduction in TE in any of the Ribo-Seq experiments conducted on *ded1* mutants carried out either here or by Guenther et al. We did observe a modest enrichment of Ded1-hyperdependent mRNAs for NCC uORFs; however, the mORF TE reductions for this group were associated with decreased, rather than increased, 5’UTR translation. Moreover, we found no evidence that these NCC uORFs have proximal downstream structures. Thus, the presence of NCC uORFs in this group of mRNAs seems to be largely incidental to their heightened Ded1-dependence, which might be attributed instead to their greater than average mORF lengths and enrichment for structured 5’UTRs.

We conducted a direct, orthologous examination of Ded1 suppression of repressive uORFs using FACS-uORF analysis of >500 native yeast 5’UTRs containing one or more inhibitory uORFs. Comparing expression of the WT and mutated uORF reporters in *ded1-cs* and *DED1* cells revealed that uORF inhibitory effects generally were not increased in the *ded1-cs* mutant in the manner expected if Ded1 stimulates mORF translation by enabling scanning PICs to bypass repressive uORFs. Consistent with this, YFP reporters harboring defined ATISs did not show a greater reduction in expression compared to reporters lacking ATISs in the *ded1-cs* mutant. These last results are especially noteworthy considering that we analyzed the *ded1-cs* mutant at 23°C, where the ATIS-associated structures should be more stable compared to the 37°C growth temperature employed by Guenther et al. to impair the thermosensitive *ded1-95* gene product, and thus should confer an even greater requirement for Ded1 to bypass ATISs and translate the mORFs in our study. Our FACS-uORF results further suggest that Ded1 inactivation did not induce nonsense-mediated mRNA decay (NMD) on ATIS-uORF containing reporters in the manner expected from increased translation of the uORFs predicted by the Ded1-START model. Importantly, we found that among >5000 5’UTRs analyzed by FACS-uORF, the presence of long, structure-prone 5’UTRs is associated with heightened dependence on Ded1 for reporter expression, including the subset devoid of uORFs. Thus, it appears that dependence on Ded1 is dictated primarily by long or structured 5’UTRs, rather than by inhibitory uORFs whose translation is enhanced by proximal downstream structures.

A small group of YFP reporters showed FACS-uORF data conforming to the Ded1-START model in exhibiting enhanced repression by the resident uORFs in *ded1-cs* vs. *DED1* cells. We pursued a subset of these exceptional mRNAs by assaying the WT and Mut_uORF reporters individually in *ded1-cs* and *DED1* cells, confirming that four harbor Ded1-suppressed uORFs. However, none of those four showed evidence of a Ded1-resolved structure downstream of the uORF in the published DMS-MaPseq data, making it unclear whether they conform to the Ded1-START model. Subsequently, we analyzed seven additional uORFs that were found to be repressive by our FACS-uORF analysis at least in *ded1-cs* cells and do harbor a Ded1-resolved structure 3’-proximal to the uORF start codon. By assaying a panel of four reporters for each uORF either containing or lacking the uORF and either with or without substitutions of uORF-proximal structurogenic nucleotides, we identified three repressive uORFs, in genes *YIL109C, YBL032W*, and *YDR481C,* whose inhibition of downstream mORF translation appears to be enhanced by downstream structures and attenuation of Ded1 function in accordance with the Ded1-START model.

Previously, we obtained evidence from profiling of scanning 40S subunits in *ded1-cs* cells that mRNAs judged to Ded1-hyperdependent on the basis of 80S profiling data frequently showed evidence for impaired scanning of 5’UTRs or reduced attachment to mRNAs by 43S PICs in vivo (5,10). We also reconstituted the strong stimulatory effects of Ded1, and its paralog Dbp1, in accelerating 48S PIC assembly on particular Ded1-hyperdependent mRNAs in a purified translation initiation system, finding that heightened dependence on Ded1 was conferred by stable stem-loops in the 5’UTRs of certain Ded1-hyperdependent mRNAs. Recently, we conducted massively parallel analysis of 48S PIC assembly (Rec-seq) on all native yeast mRNAs in the purified initiation system and found that Ded1 preferentially stimulated PIC assembly at the mORFs of mRNAs harboring long, structure-prone 5’ UTRs, including the majority of Ded1-hyperdependent mRNAs identified by 80S profiling of *ded1* mutants. Although Ded1 suppressed PIC assembly at numerous uORFs in the Rec-seq analysis, possibly by unwinding structures downstream of their start codons, the magnitude of PIC reduction at these uORFs was far too small to account for the increased PIC assembly at the corresponding mORF start codons conferred by Ded1 (18).

There is no doubt that proximal downstream structures can stimulate uORF recognition by scanning PICs (14) (15,16), and the mechanism of uORF-dependent translational control by Ded1 does appear to operate on a small number of uORFs equipped with Ded1-resolved proximal downstream structures. However, our results do not support the view that such structure-associated uORFs generally underlay the regulation of translation initiation by Ded1, at least when cells are growing vegetatively in rich medium. Instead, our work shows that, under these conditions, Ded1 primarily functions by unwinding structures that impede PIC attachment, 5’UTR scanning or recognition of the mORF start codon independently of the presence of translated uORFs. It is possible however that Ded1 acts more broadly according to the Ded1-START model during the starvation conditions that induce meiosis and sporulation, where it was found that Ded1 protein expression is greatly reduced and increased ATIS translation in *DED1* cells was reported (17).

## DATA AVAILABILITY

All primary data obtained from RNA-Seq and Ribo-Seq are deposited at the Gene Expression Omnibus data repository under a pending accession number. All processed data used to generate the figures are provided in Supporting Files S1-S9.

## ACKNOWLEDGEMENTS

We are grateful to all members of our laboratories for helpful comments and suggestions. We thank the NHLBI DNA Sequencing and Genomics Core and NIH HPC Biowulf cluster for computational support. This research was supported [in part] by the Intramural Research Program of the National Institutes of Health (NIH). The contributions of the NIH author(s) are considered Works of the United States Government. The findings and conclusions presented in this paper are those of the author(s) and do not necessarily reflect the views of the NIH or the U.S. Department of Health and Human Services. This work was also supported in part by an Extramural research grant from NIGMS (R35GM145317 to CJM) and by the Department of Biotechnology, India (BT/RLF/Re-entry/55/2017 to NDS).

## Author contributions

R.K.: Conceptualization, Investigation, Formal analysis, Methodology, Validation, Visualization, Writing—original draft, Writing—review & editing, Data Curation; G.M: Investigation, Formal analysis, Methodology, Data Curation; N.S.: Conceptualization, Investigation, Formal analysis, Methodology, Validation, Visualization, Writing—original draft, Writing—review & editing, Data Curation; J.M.: Conceptualization, Formal analysis, Writing—original draft, Writing—review & editing, Funding acquisition, Supervision; A.G.H.: Conceptualization, Formal analysis, Writing—original draft, Writing—review & editing, Funding acquisition, Supervision.

## SUPPLEMENTARY MATERIAL

## SUPPLEMENTARY FIGURES

**Figure S1.**
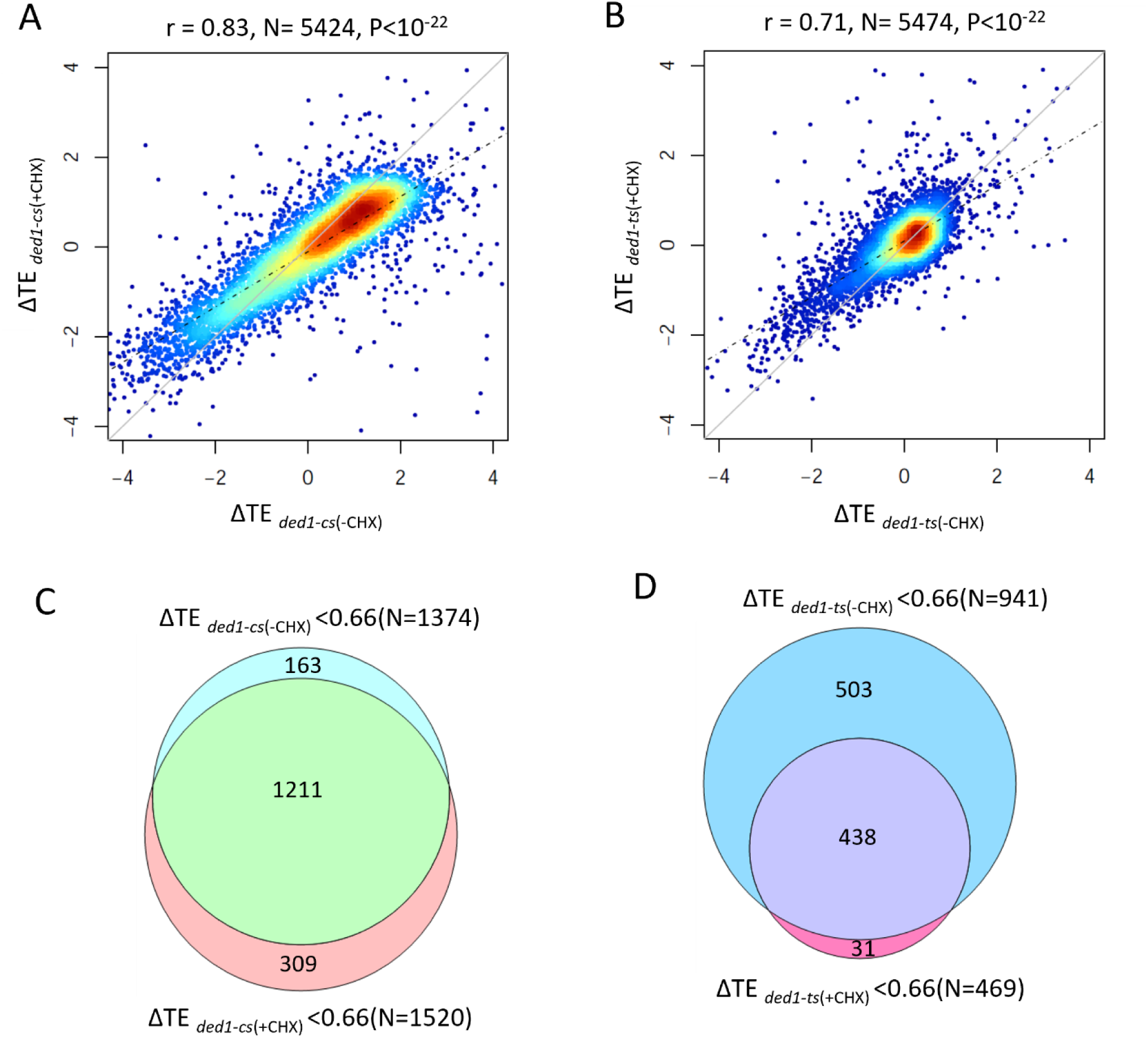
Genome-wide translational changes of mORFs conferred by *ded1-cs* and *ded1-ts* mutations are influenced minimally by treatment of cells with cycloheximide. **(A)** Density plot of log_2_ fold-changes in TE for mORFs conferred by *ded1-cs* versus *DED1* for cells treated (+CHX) or untreated with CHX (-CHX), for 5,424 expressed genes, excluding from the plot (but not the analysis) genes with −6 > log_2_ΔTE >6 values to expand the axes. The dotted line is the determined regression line; grey solid line is the theoretical regression line for identical changes in ΔTE values. Individual genes are shown by blue filled circles; the coloring indicates increased density of genes (red is maximum). TE change values of +CHX samples were taken from (1). Pearson correlation coefficients (r) and associated P-values are indicated. **(B)** Similar to (A), but for 5,474 expressed genes quantified in the *ded1-ts*(+CHX) versus *ded1-ts*(-CHX) experiments. **(C)** Overlap between mRNAs exhibiting ≥1.5-fold decreases in TE at FDR<0.05 in *ded1-cs* versus *DED1* cells under CHX -treated or -untreated conditions. **(D)** Similar to (C), but for mRNAs exhibiting significant TE reductions in response to the *ded1-ts* mutation.

**Figure S2.**
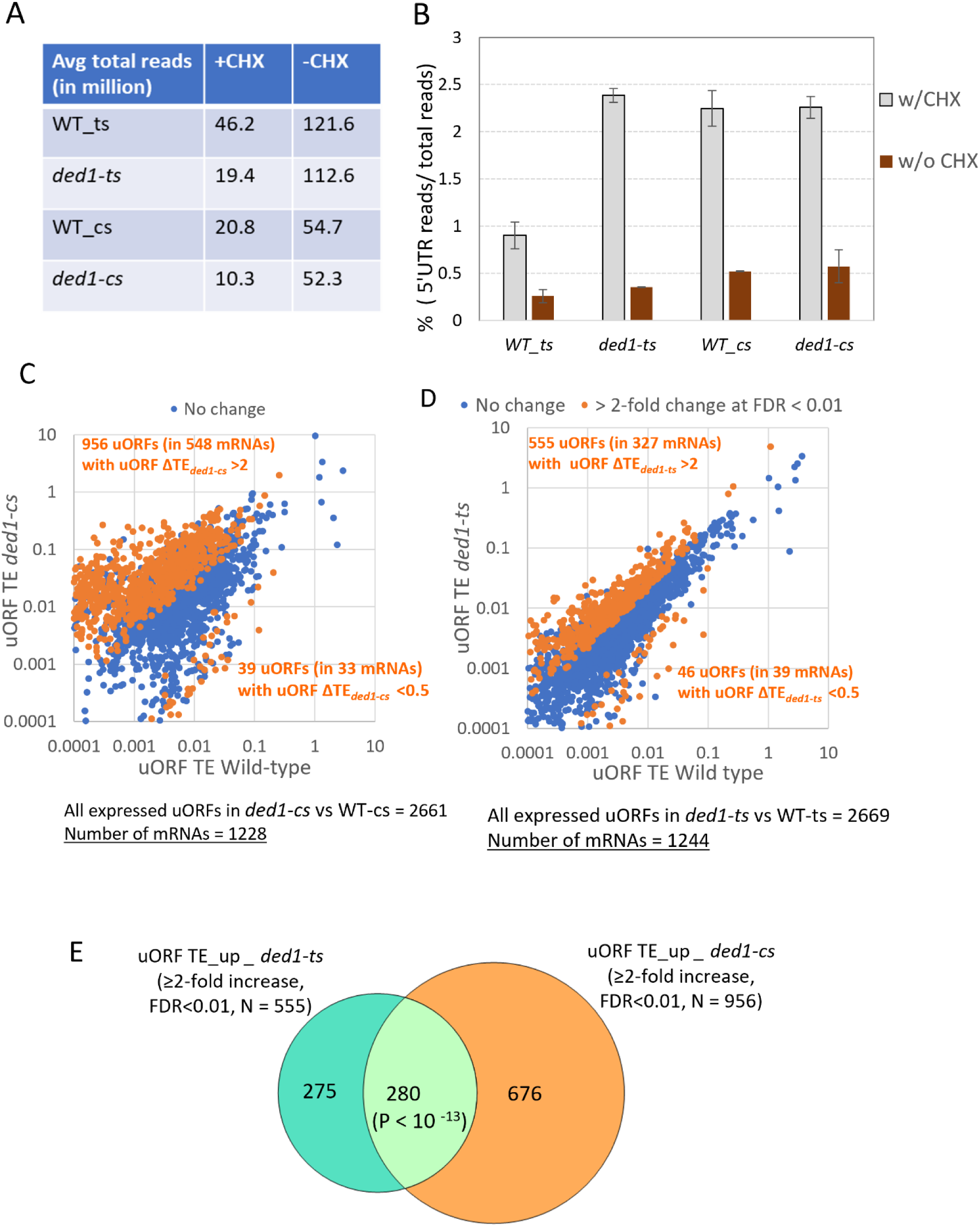
Despite reduced overall RPF occupancies in 5’UTRs, a significant number of mRNAs show evidence of increased uORF translation in *ded1* cells in -CHX vs. +CHX experiments. **(A)** Average number of reads mapped to the yeast transcriptome from the two replicates for the indicated *ded1* mutants and the isogenic *DED1* strains from -CHX and +CHX experiments. **(B)** Fraction of total 80S RPFs mapped to 5’UTRs in the *ded1-ts, ded1-cs,* and respective *DED1* samples from cultures treated (grey) or untreated (maroon) with CHX, plotting data from two biological replicates for each strain. **(C-D)** Scatterplots of uORF TEs in the indicated mutant versus corresponding *DED1* strain from -CHX experiments for (C) *ded1-cs*, N=2661, or (C) *ded1-ts*, N=2669, with N equaling the total number of expressed uORFs analyzed in each comparison. uORF TE values were determined by DESeq2 using biological replicates of each strain for genes having ≥ 128 mRNA and ≥ 8 RPF reads in uORFs in the four samples (two replicates each of *DED1* and *ded1* mutant). Genes exhibiting ≥2-fold changes in uORF TE in *ded1* mutant versus *DED1* cells at FDR *<* 0.01 are highlighted in orange and the number of uORFs and mRNAs exhibiting a substantial increase or decrease in uORF TE are indicated in orange type. **(E)** Venn diagram of overlap between uORFs exhibiting ≥2-fold increases in TE in *ded1-cs* or *ded1-ts* vs. *DED1* cells., with the P value for significance of overlap based on the hypergeometric distribution.

**Figure S3.**
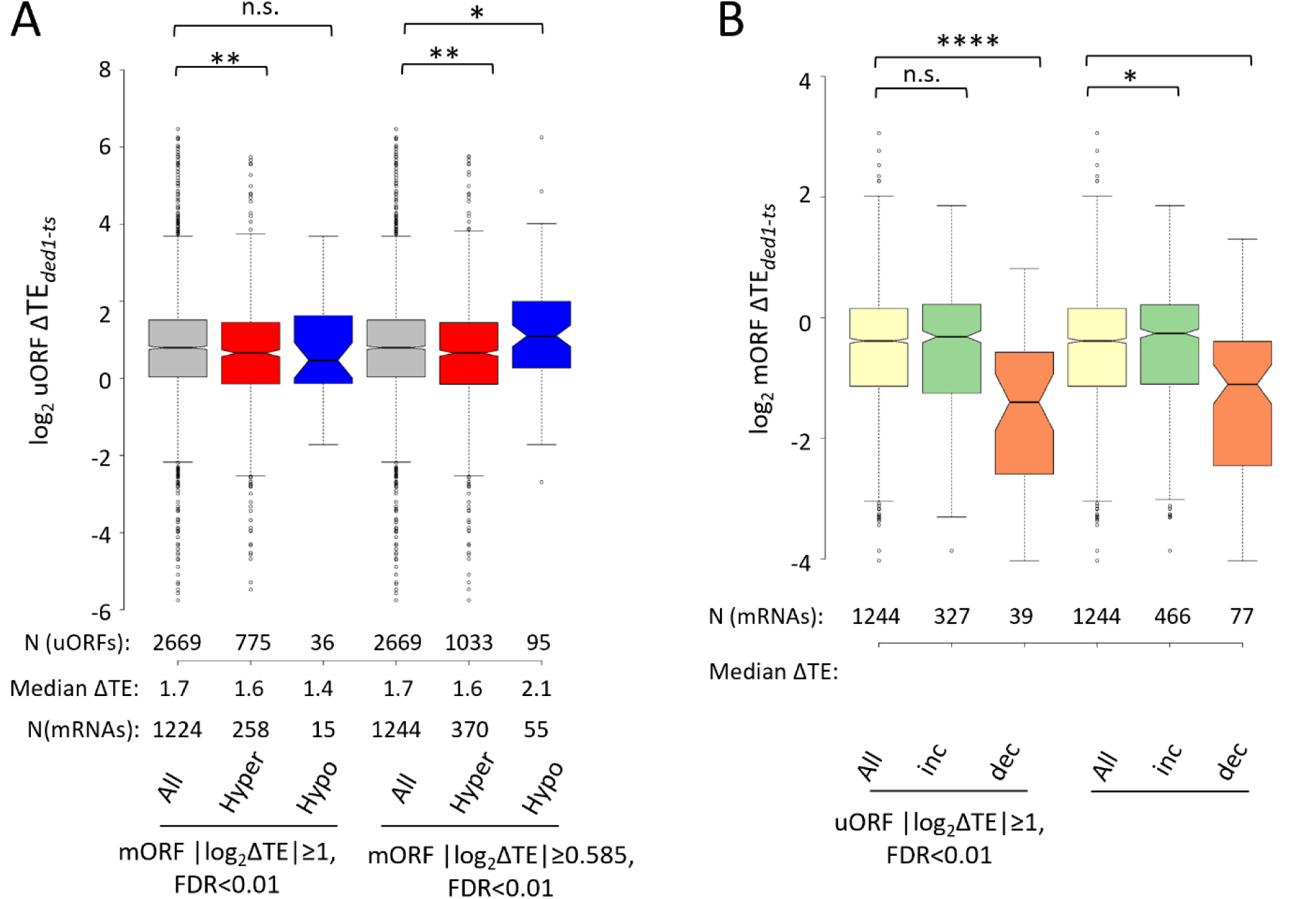
Increased uORF translation in *ded1-ts* cells is generally not accompanied by reduced translation of the downstream mORFs. (A)-(B) Analysis identical to Figs. 2A & C but conducted using *ded1-ts* (-CHX) Ribo-Seq data. Results of Mann-Whitney U-tests are summarized as: ****, P<0.0001; **, P<0.01; *, P<0.05; n.s., not significant, P>0.05.

**Figure S4.**
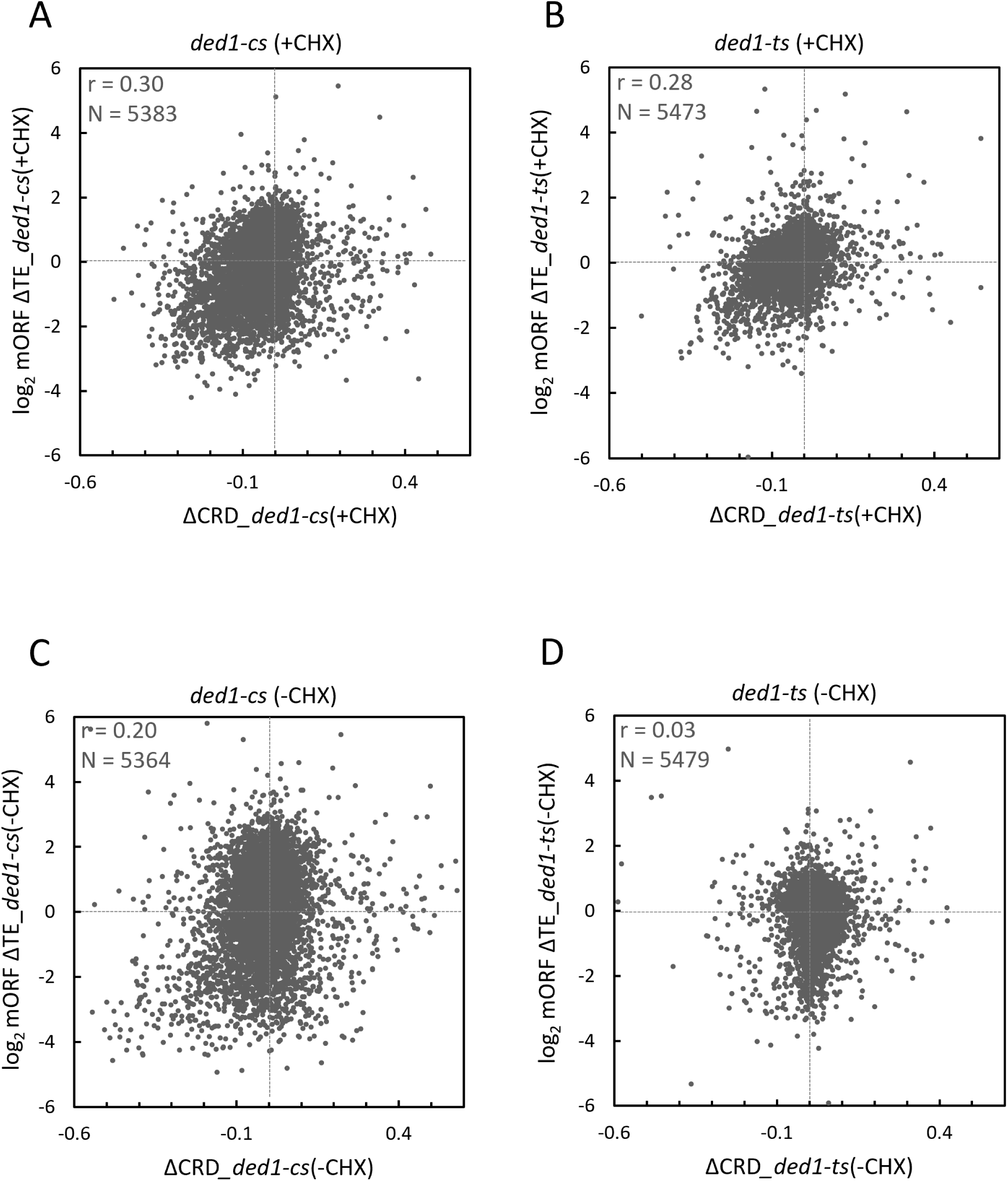
Cycloheximide pre-treatment of cells increases positive correlations between changes in TE and changes in center of ribosome density in *ded1* mutants vs. *DED1* cells. (A-D) Scatterplots comparing changes in TE and changes in center of ribosome density (CRD) for all expressed genes in *ded1* mutant versus *DED1* cells for (A) *ded1-cs*(+CHX) (B) *ded1-ts*(+CHX) (C) *ded1-cs*(-CHX) and (D) *ded1-ts*(-CHX) experiments. Total number of genes in each comparison (N) and Pearson correlation coefficients (r) values are shown.

**Figure S5.**
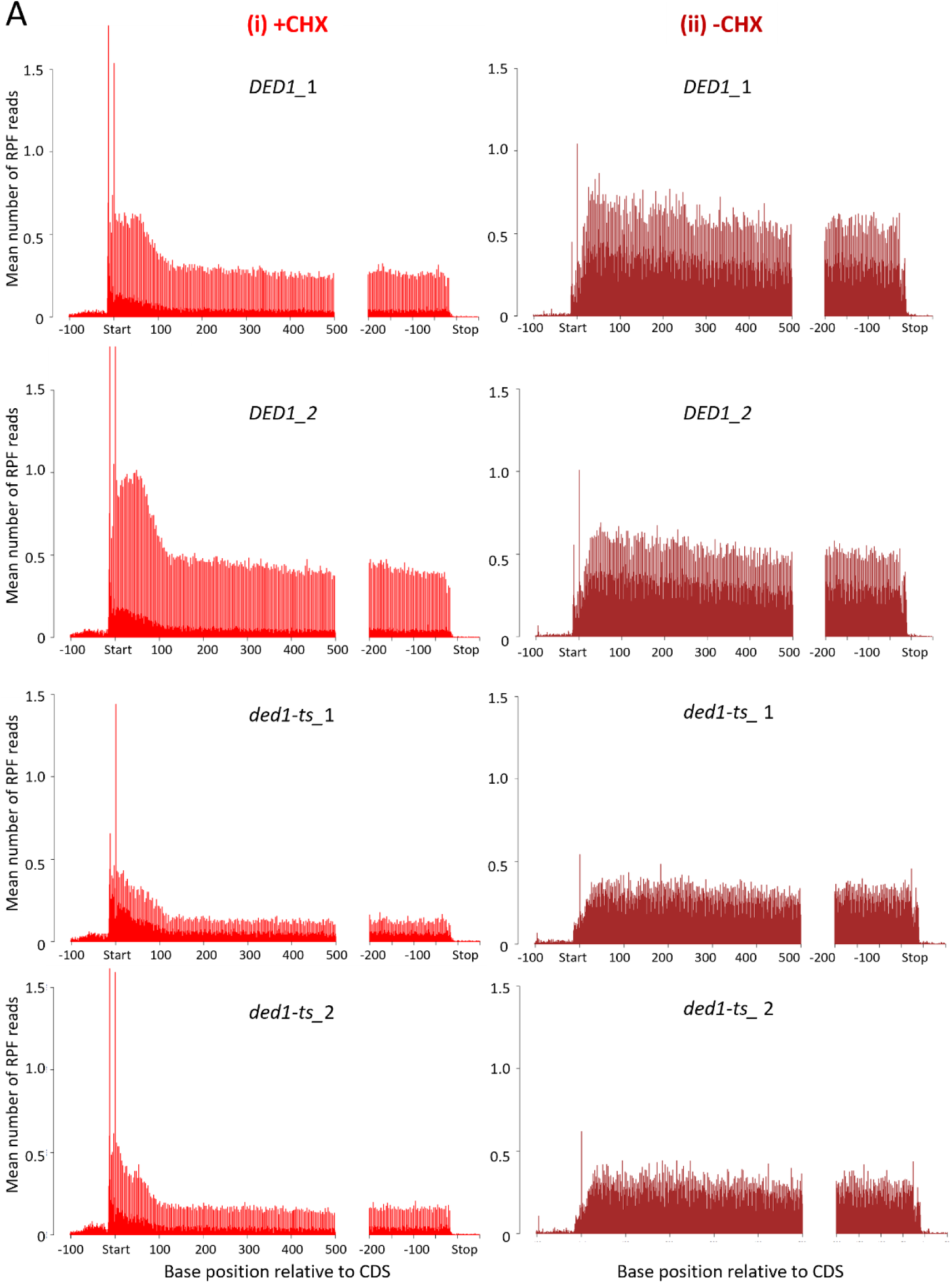
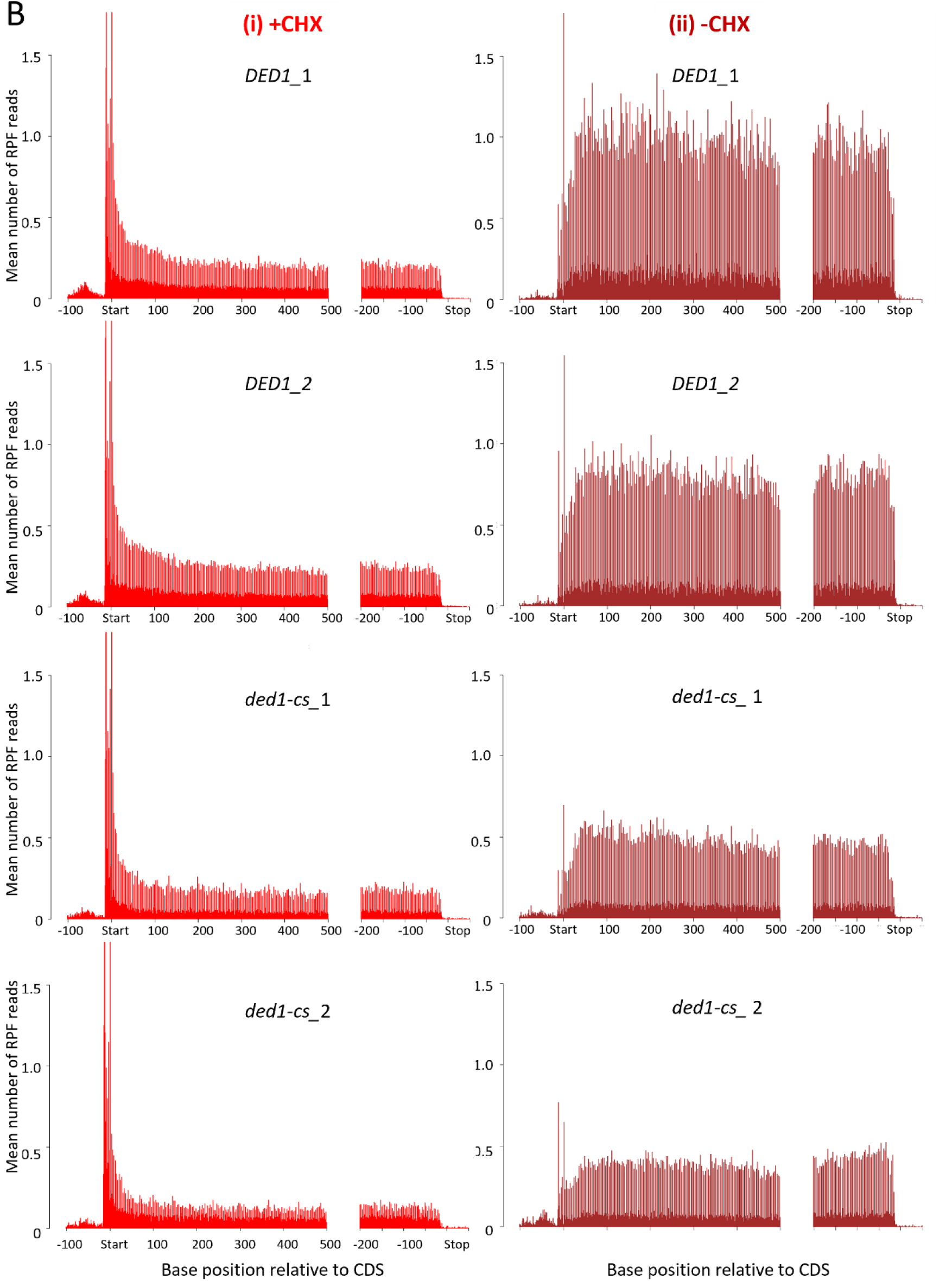
Pre-treating cells with cycloheximide increases RPF densities in the beginnings of mORFs throughout the translatome. Metagene plots of the mean number of RPF reads averaged over all expressed genes at each nucleotide from bases -100 to -1 and 1 to 500 relative to the ATG start codon (position 1, “Start”) and from bases -200 to -1 and 50 bases downstream of the stop codon (position 1, “Stop”) calculated from Ribo-Seq data for two biological replicates (_1, _2) of *DED1* and *ded1-ts* strains (A) or *DED1* and *ded1-cs* strains (B) cultured at the non-permissive temperature for the corresponding *ded1* mutant, with cells pre-treated with CHX ((i), +CHX) or with CHX added only to cell lysates ((ii), -CHX).

**Figure S6.**
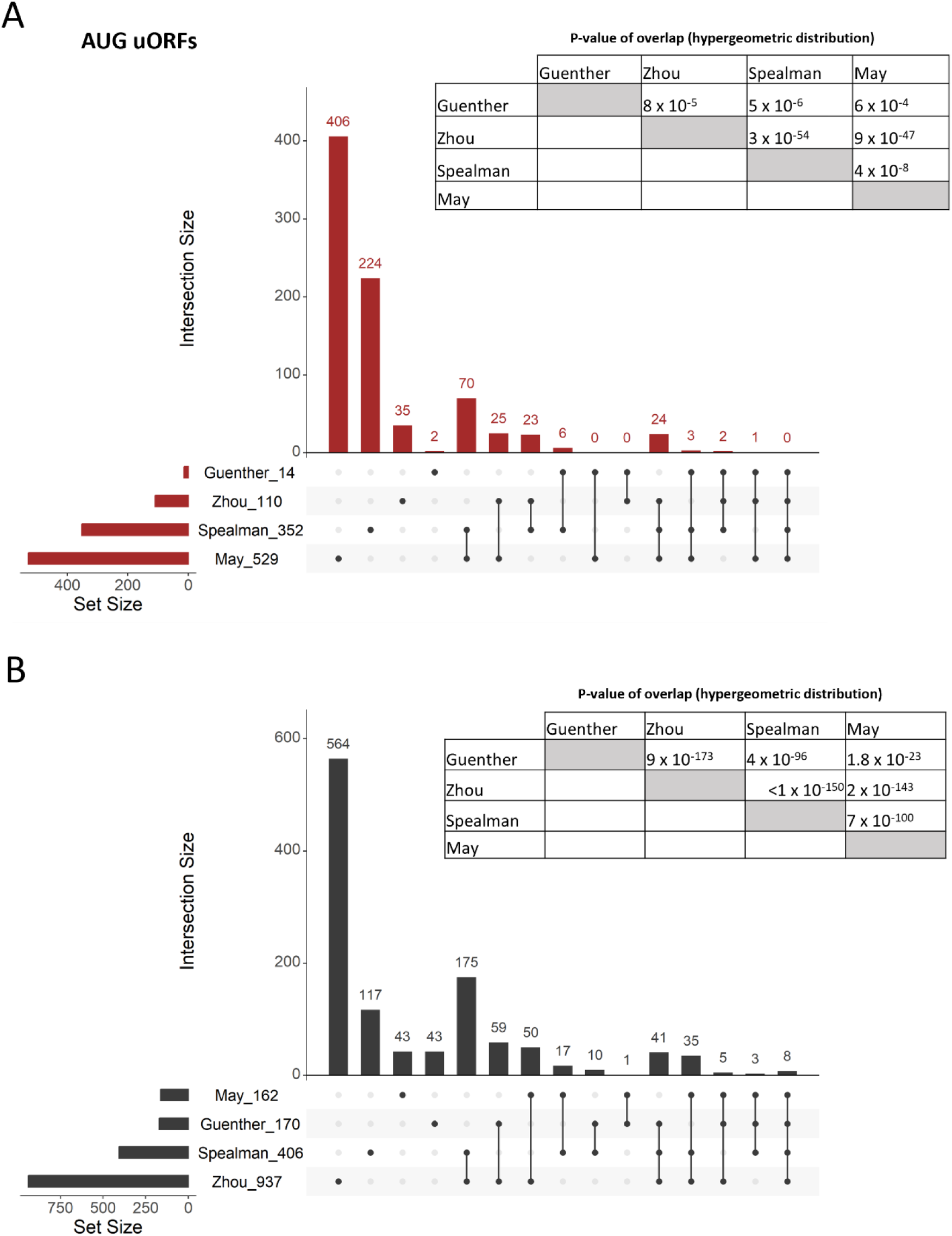
Highly significant overlaps among independent compilations of uORFs whose translation has been evaluated by Ribo-Seq analysis of *ded1* mutants. **(A-B)** Upset plots depicting overlaps between sets of 14, 110, 352, and 529 AUG-initiated uORFs (A) or 162, 170, 406, and 937 NCC uORFs (B) identified by Guenther et al. (2), Zhou et al. (3), Spealman et al. (4), and May et al. (5), respectively. The first four columns list the numbers of uORFs unique to each set, while the last 11 columns depict the numbers of uORFs shared among 2, 3, or all 4 sets connected by lines. The significance of overlaps is indicated by the insets tabulating the P-values determined for all six pairwise comparisons using the hypergeometric distribution, based on 3156 total AUG uORFs and 74353 NCC uORFs identified in the sequences of annotated 5’UTRs for all yeast genes.

**Figure S7.**
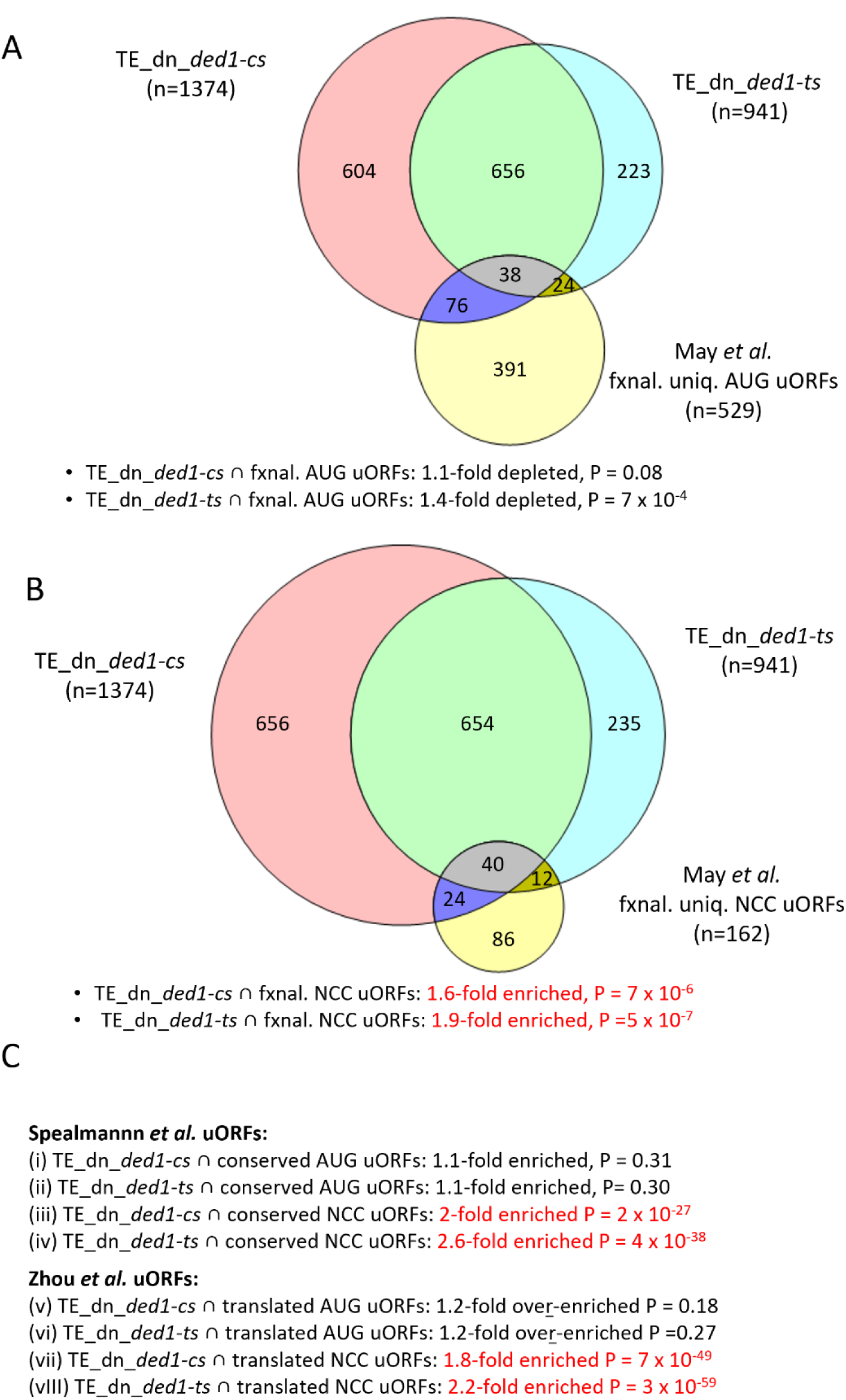
Ded1-hyperdependent mRNAs are not significantly enriched for mRNAs with AUG-initiated uORFs. **(A-B)** Venn diagram showing overlap between Ded1-hyperdependent mRNAs identified in *ded1-cs* or *ded1-ts* cells in -CHX experiments (mRNAs exhibiting ≥1.5-fold decrease in TE at FDR<0.05 in *ded1* mutant versus *DED1* cells, reported in Fig. S1C-D) and either 529 mRNAs containing functional AUG uORFs (A) or 162 mRNAs containing functional NCC uORFs (B) that were compiled by May et al. (5). These mRNA groups omit mRNAs that contain both AUG and NCC uORFs but include mRNAs that contain multiple AUG uORFs or multiple NCC uORFs, and thus contain mRNAs that uniquely harbor AUG or NCC functional uORFs (fxnal. uniq. AUG or NCC uORFs). **(C)** Same as (A-B) except analyzing evolutionarily conserved uniquely AUG (n=352) or uniquely NCC (n=406) uORFs identified by Spealmann et al. (4), or analyzing translated uniquely AUG (n=110) or uniquely NCC (n=937) uORFs identified by Zhou et al. (3).

**Figure S8.**
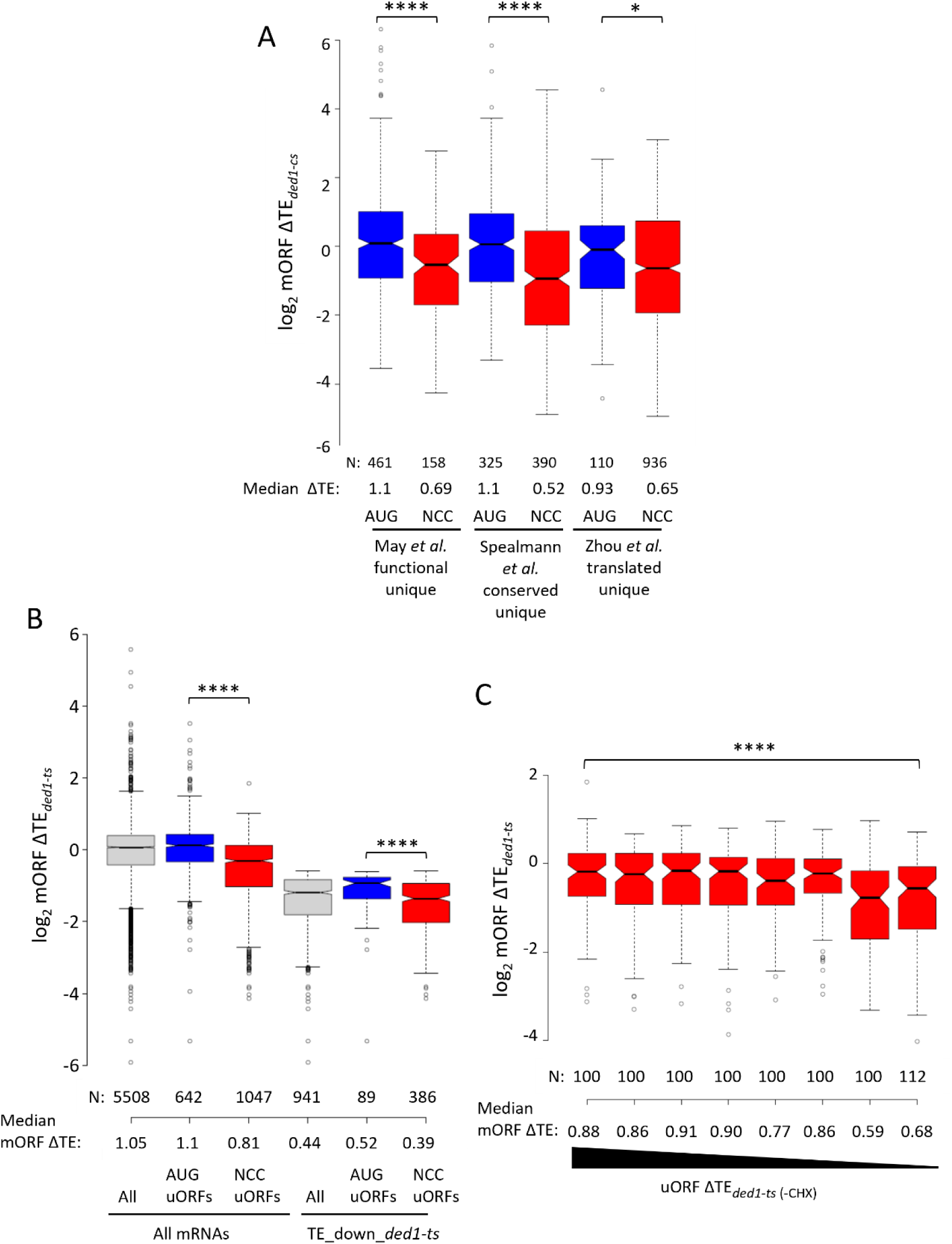
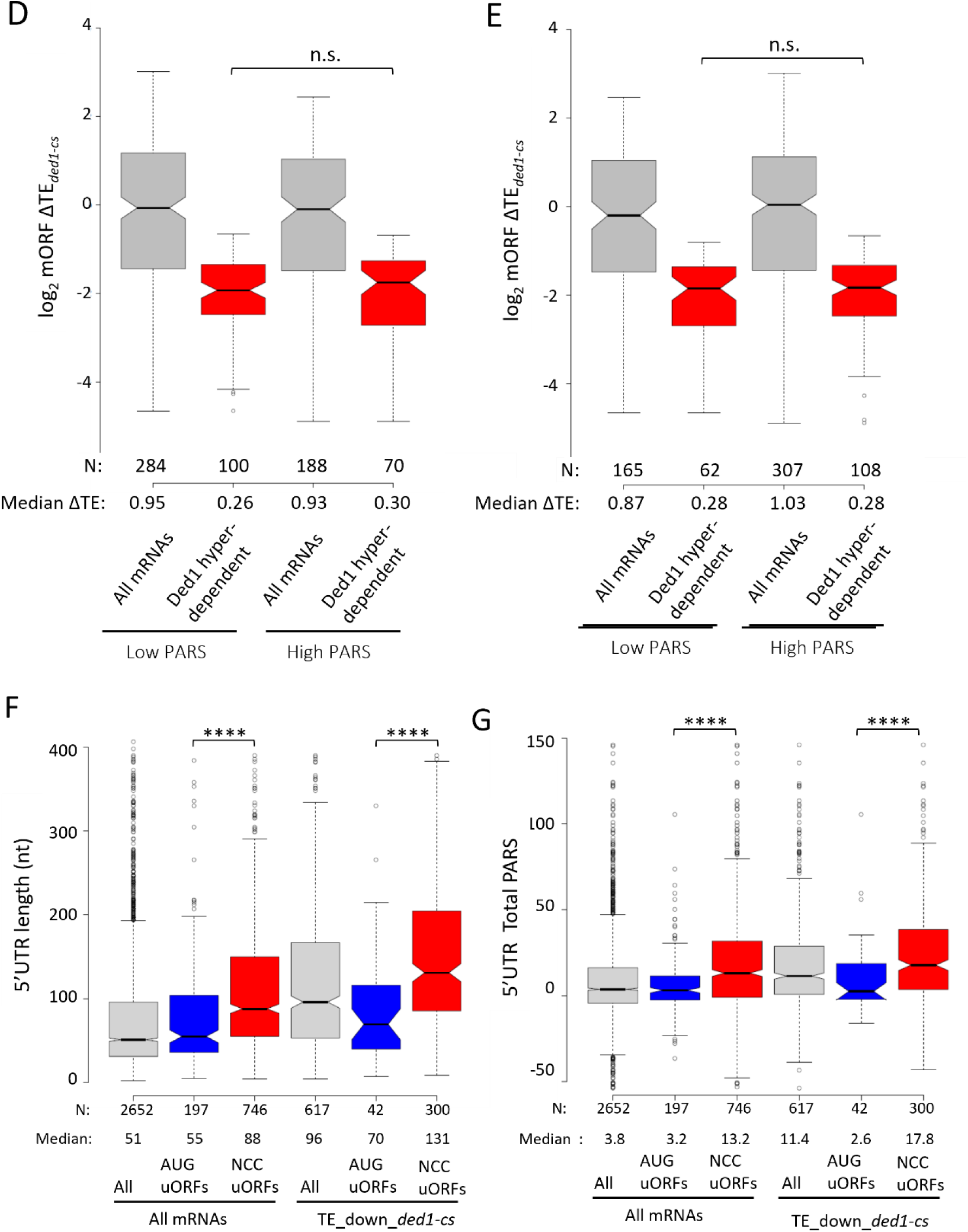
Presence of NCC-initiated uORFs is associated with, but not functionally linked to, heightened dependence on Ded1 for translation of mORFs. **(A)** Notched box plots comparing log_2_ fold-changes in mORF TE observed in *ded1-cs* versus *DED1* cells from -CHX cultures for mRNAs containing either unique AUG uORFs (cols. 1,3 & 5) or NCC uORFs (cols. 2,4 & 6) compiled by May et al. (5), (cols. 1-2), Spealmann et al. (4) (cols. 3-4), or Zhou et al. (3) (cols. 5-6) and described in Fig. S5A-C. (B-C) Ded1-hyperdependence conferred by the *ded1-ts* mutation is largely independent of increased AUG- or NCC-initiated uORF translation. Analysis identical to Fig. 3A-B conducted using *ded1-ts* (-CHX) Ribo-Seq data. **(D-E) Higher propensity for secondary structures downstream of NCC uORFs does not confer heightened dependence on Ded1 for translation of mORFs.** Similar to Fig. 3C but summing up the PARS scores for nucleotides 16-30 (A) or 16-60 (B) downstream of the start codon of the NCC uORFs. **(F-G) Longer and structure prone 5’UTRs of mRNAs containing NCC uORFs confer heightened dependence on Ded1 for translation of mORFs.** Notched block blots of the distributions of 5’UTR lengths (A) or Total PARS scores (B) for the same groups of mRNAs analyzed in Fig. 3A for which data was reported by Kertesz et al. (6). Results of Mann-Whitney U-tests for all panels are summarized as: ****, P<0.0001; ***, P<0.001; n.s., not significant, P>0.05.

**Figure S9.**
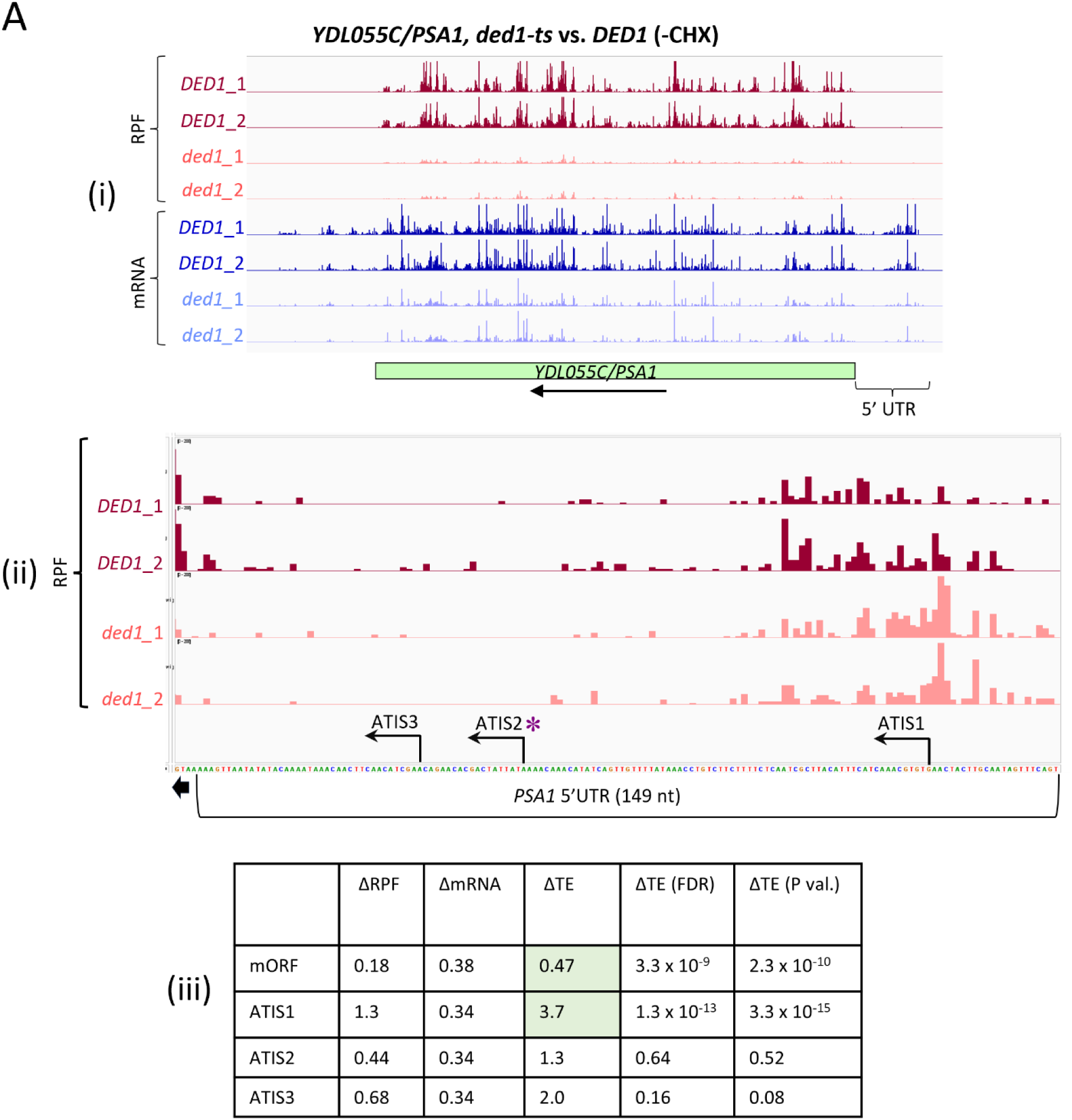
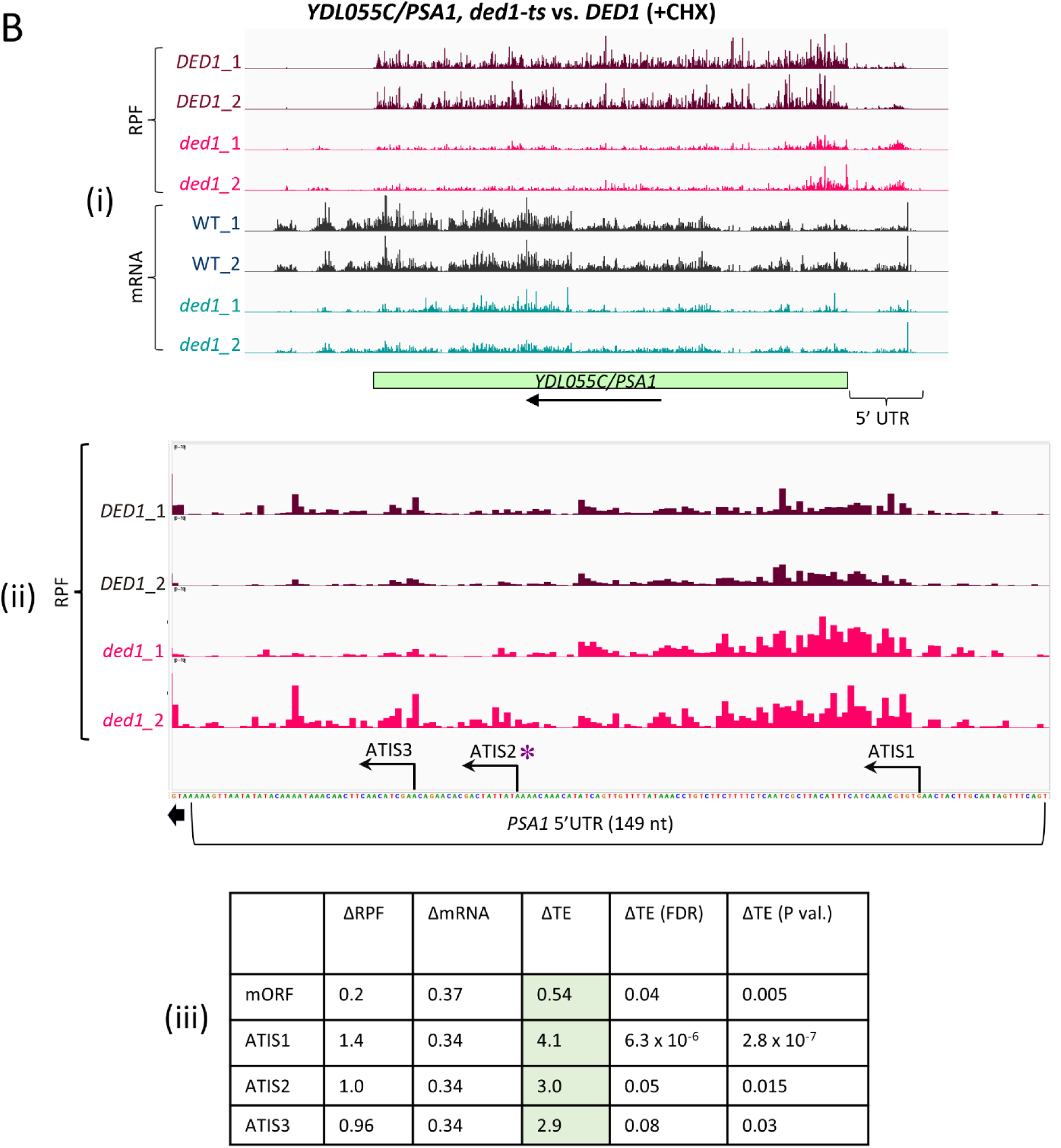
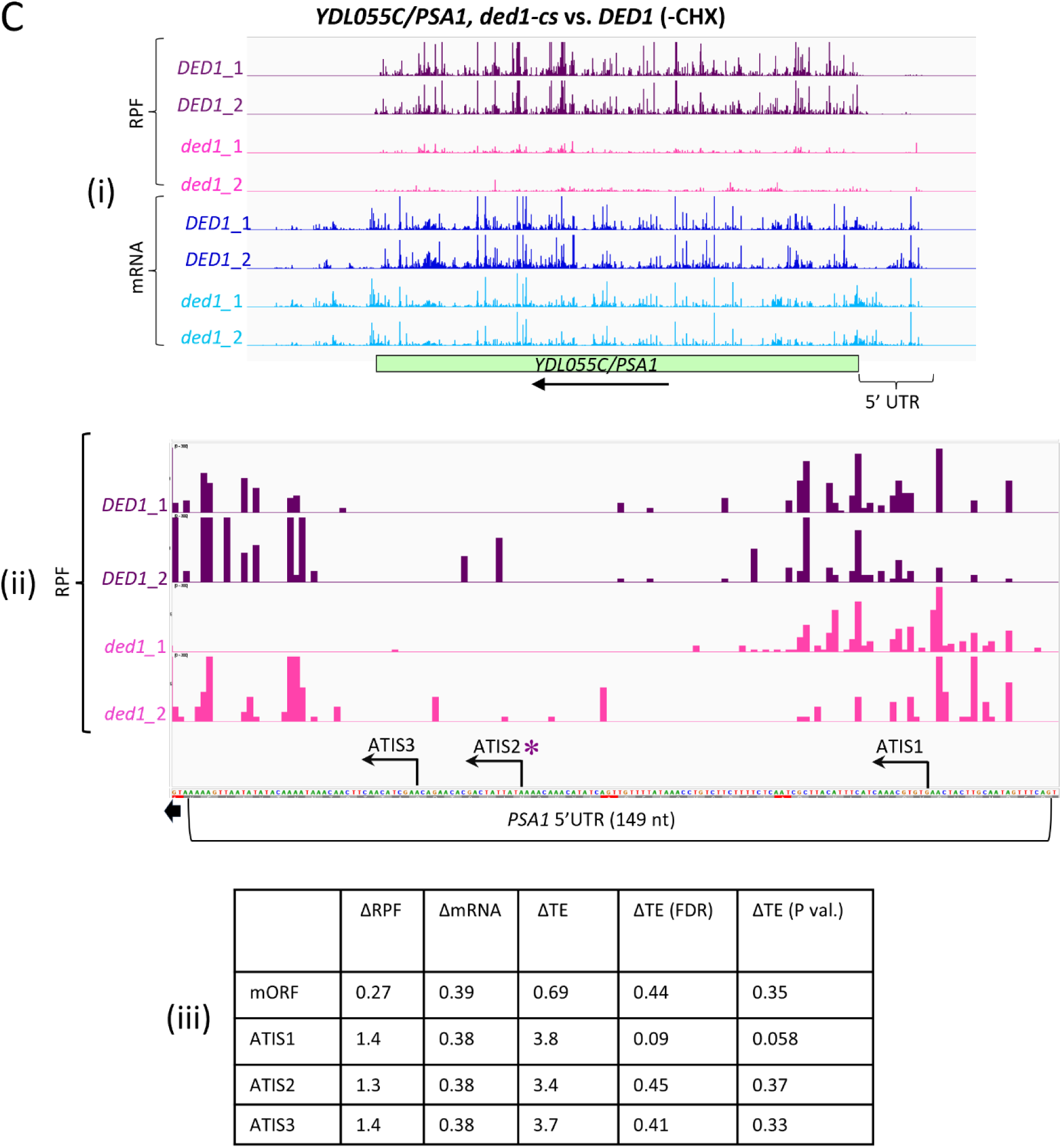
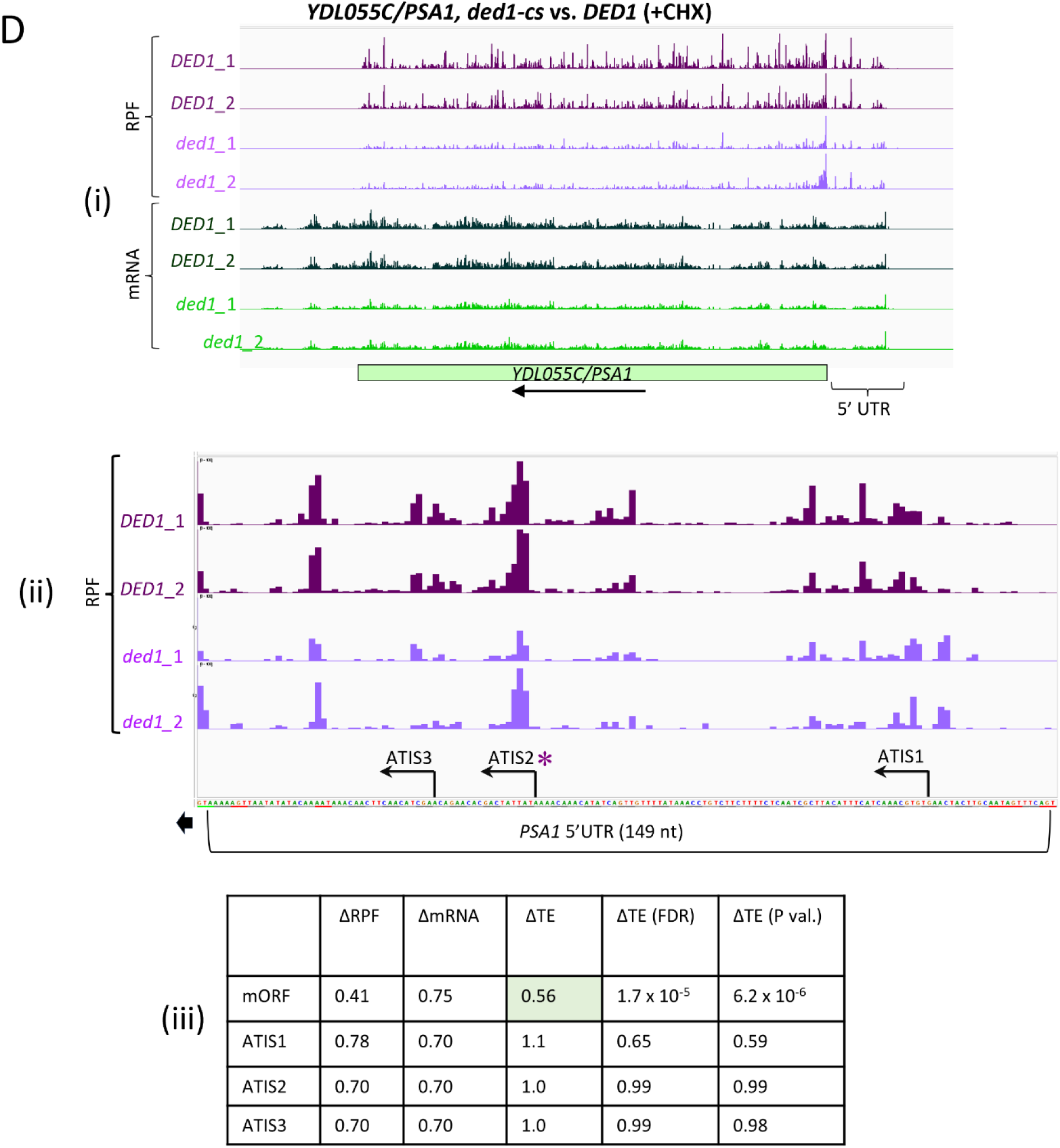
*ded1* mutations do not confer increased translation of ATIS2 associated with decreased translation of the *PSA1* mORF in Ribo-Seq conducted without CHX-treatment of cells. **(A)-(D). Panels (i)-(ii)** Gene-browser depictions of RPF and mRNA counts in the 5’UTR and mORF (i) or only the 5’UTR (ii) of *PSA1* determined by Ribo-Seq analysis of two replicates (_1, _2) of *DED1* and *ded1-ts* cells (A-B) or *DED1* and *ded1-cs* cells (C-D) prepared with CHX treatment of either lysates (-CHX, panels A & C) or cells (+CHX, panels B & D). The GUG, AUA, and AAG start codons of three NCC ATISs (dubbed 1, 2, and 3) are indicated above the 149 nt 5’UTR sequence of *PSA1* in (ii), with the asterisk indicating the importance of ATIS2 in Ded1-mediated enhanced translation of the *PSA1* mORF, as reported by Guenther et al. (2). All eight tracks in panel (i) or four tracks in panel (ii) have the same ranges of read counts. **(iii)** Tabulated changes in RPFs, mRNA, and TEs, with FDR and P-values for the ΔTE values, determined by DESeq2 analysis, conferred by the *ded1* mutation and condition of Ribo-Seq analysis (-CHX vs. +CHX) indicated at the top of the figure. Statistically significant TE changes (P < 0.05) are indicated with light green shading of the relevant cells. Whereas the mORF TE is reduced by the *ded1* mutation in three of the four datasets (panels A, B & D), an inversely correlated TE increase for ATIS2, in the manner predicted by the START model, was observed only for the *ded1-ts* mutation in the +CHX dataset (panel B). Slightly different ΔmRNA values for mORFs versus ATISs 1-3 in each panel result from differences in the number of entries in the fpcount input files subjected to two separate DESeq2 analyses of TE changes for uORFs or mORFs. For the uORF TE analyses, the fpcount file had 8962 entries comprised of 3315 uORFs and 5647 CDSs, whereas only the 5647 CDS entries comprised the fpcount file used for mORF TE analysis. We included the CDS entries with the uORF entries for the uORF TE analysis to reduce imprecision resulting from generally low read counts for the short uORFs compared to the much higher read counts for the longer mORFs encoded within the same transcripts, as done previously (7).

**Figure S10.**
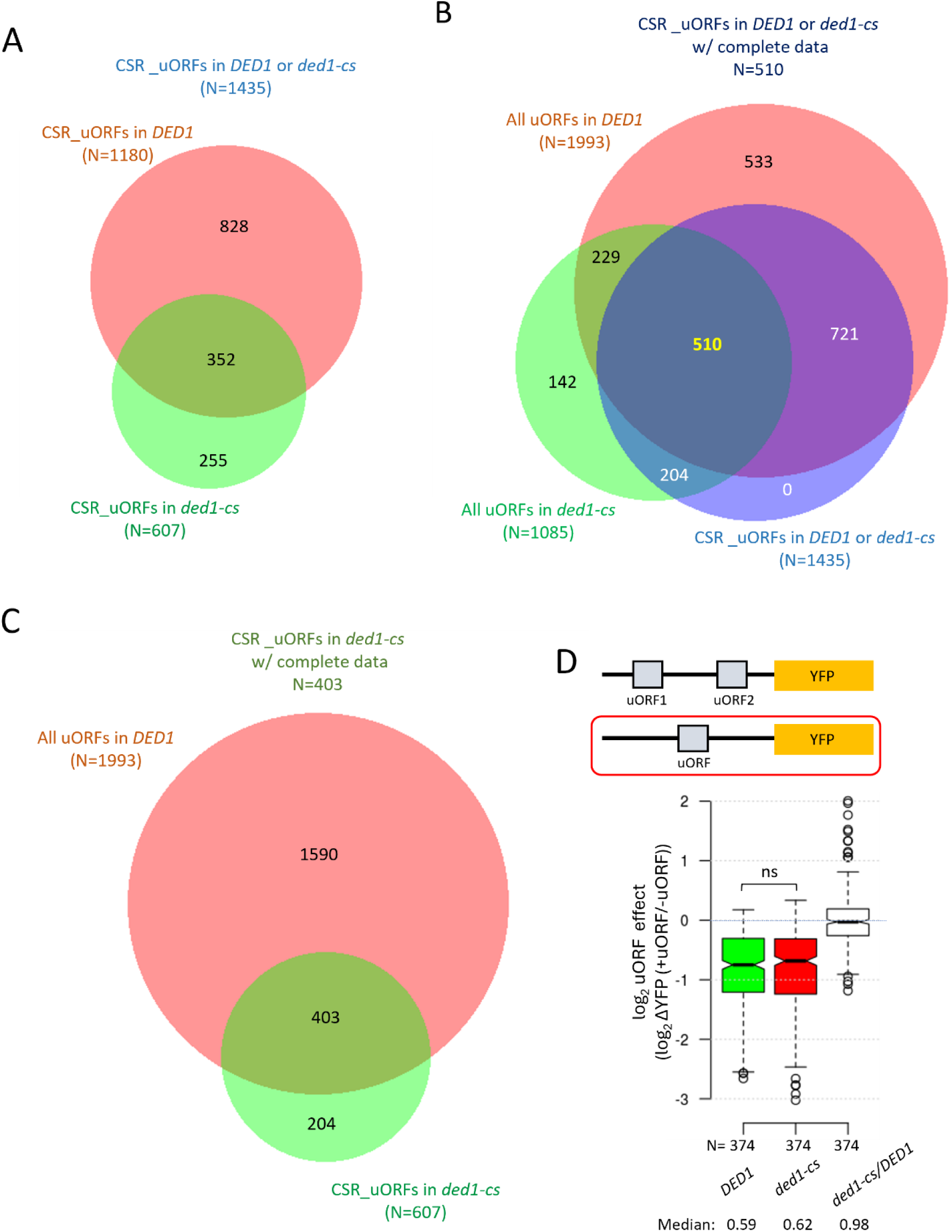
Compilation of results from MPRA of the FACS-uORF library in *ded1-cs* vs. *DED1* cells. **(A)** Venn diagram depicting overlap between the two sets of YFP reporters whose uORFs were judged to CSR by comparing expression of replicates for the matched WT_uORF and Mut_uORF in the *DED1* strain (N=1180) or *ded1-cs* mutant (N=607). **(B)** Venn diagram depicting overlap between the set of 1435 reporters whose uORFs were judged to be CSR in either *DED1* or *ded1-cs* cells (union of two sets in (A), blue), the 1993 reporters for which replicate data were obtained for both WT_uORF and Mut_uORF reporters in *DED1* cells (red), and the 1085 reporters with replicate data for both WT_uORF and Mut_uORF reporters in *ded1-cs* cells (green). **(C)** Venn diagram depicting overlap between the set of 1993 reporters for which replicate data was obtained for both WT_uORF and Mut_uORF reporters in *DED1* cells (also analyzed in (B), red) and the 607 reporters whose uORFs were judged to CSR in *ded1-cs* cells (also analyzed in (A), green). **(D)** Analysis identical to that in Fig. 4B(ii) but for the subset of 374 CSR uORFs from the group of 510 described in panel C that contain only a single uORF that was mutated in the FACS-uORF library, as depicted with red outline in the schematic above.

**Figure S11.**
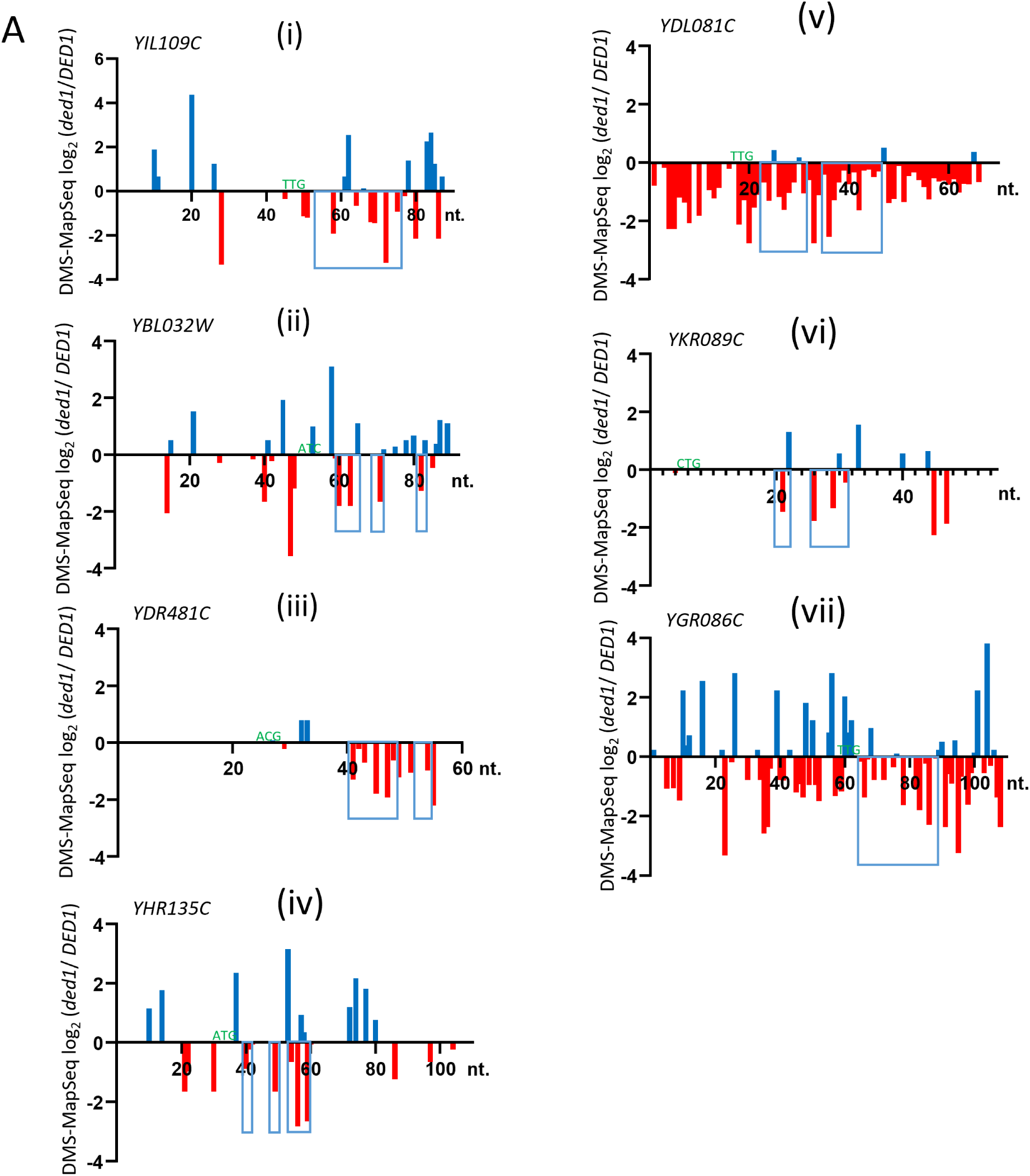
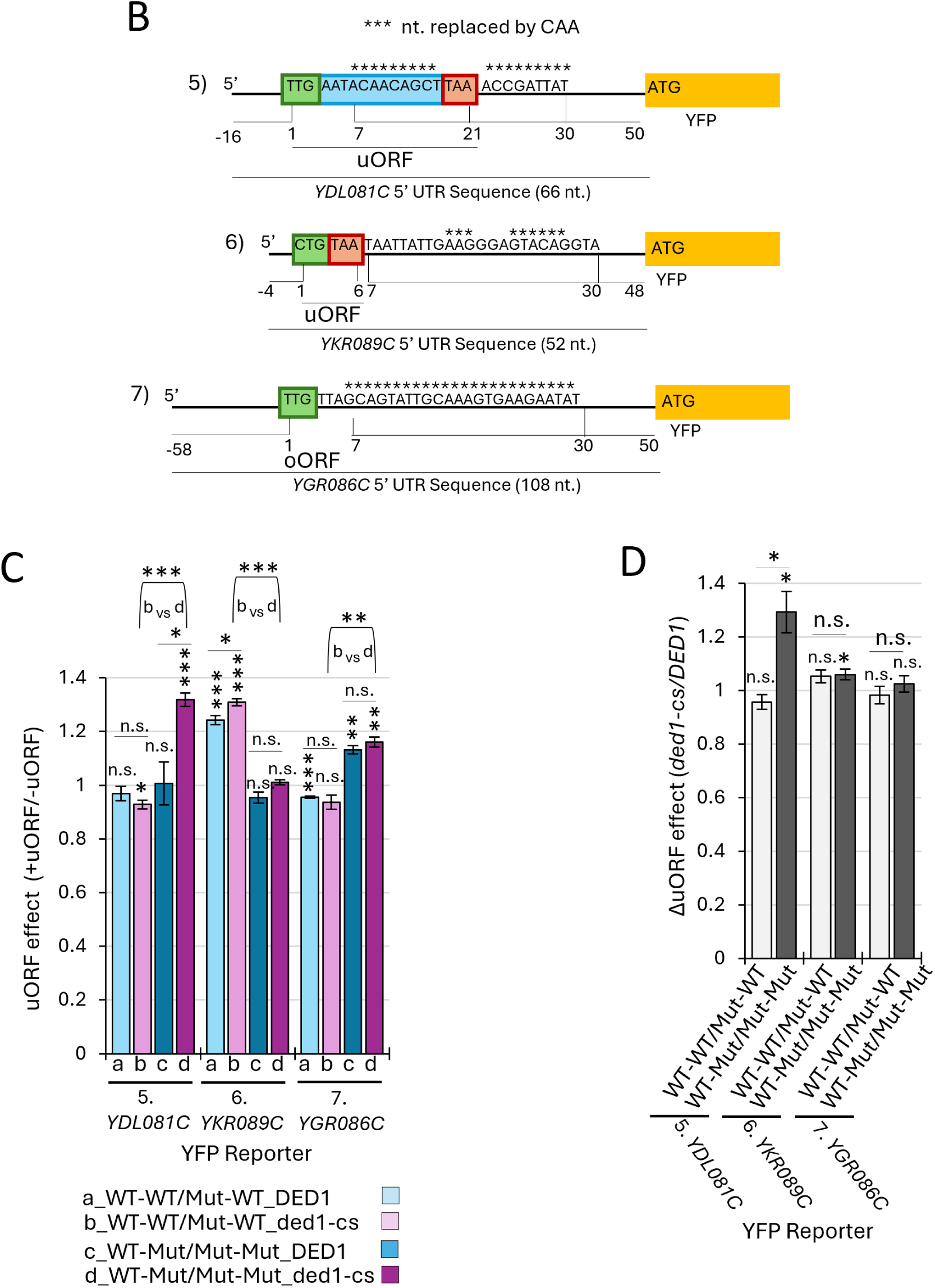
Reporter analyses indicating that uORFs at *YDL081C, YKR089C,* and *YGR086C* do not conform to the Ded1-START model. **(A)** Compilations of the published DMS-MapSeq data from Guenther et al. (2) presented as log_2_ ratios of read counts (normalized reverse transcription stops) plotted against nucleotide position (nt) in the 5’UTR (5’ to 3’ from the cap) obtained from *ded1-ts* mutant vs. *DED1* cells after 5 min at the non-permissive growth temperature for (i)-(iv) the four genes subjected to mutational analysis of uORFs and proximal downstream structures with results presented in Fig. 6, and (v)-(vii) the three genes similarly analyzed in panels B-D below. Negative values on y-axes (red bars) signify nucleotides more unwound (and DMS-modified) in *DED1* vs. *ded1-ts* cells. Violet boxes delineate regions downstream of the uORF start codons (green type on or above x-axis, not exactly to scale) that were subjected to CAA replacements of nucleotides unwound by Ded1 to disrupt structures 3’-proximal to the uORF start codons that are unwound by Ded1, which were separately substituted with AAG or AAA triplets to mutate the uORF start sites in the presence or absence of CAA substitutions. **(B)-(D)** Depiction of *YFP* reporter constructs (B) and analyses of reporter expression in *DED1* and *ded1-cs* cells for three uORFs at genes *YDL081C, YGR086C,* and *YKR089C* (C-D) presented exactly as described above for the four reporters examined in Figs. 6A, C & D, summarizing data from 3 biological replicates for each reporter/strain combination. Results for all three gene/uORFs analyzed here depart from one or more predictions of the Ded1-START model, as follows. The data in (C) for *YDL081C* and *YGR086C* reveal that their uORFs are not significantly more inhibitory in *ded1-cs* vs. *DED01* cells, exhibiting essentially the same uORF effects in the two strains, when the WT downstream structures are present (reporters 5 & 7, col. a vs. b); and eliminating the downstream structures unexpectedly renders the uORFs stimulatory, now exhibiting uORF effects >1, in both *DED1* and *ded1-cs* cells for *YGR086C* (reporter 7, col. c vs. a and col. d vs. b) or only in *ded1-cs* cells for *YDL081C* (reporter 5, col. d vs. b). For *YKR089C,* the uORF appears to be stimulatory rather than inhibitory in both strains, conferring uORF effects >1 in the presence of the structures (reporter 6, col. a-b); and removing the downstream structure renders the uORF ineffectual, now exhibiting a uORF effect of ∼1.0 (reporter 6, col. c vs. a and d vs. b). Consequently, the results in (D) show that none of the three reporters exhibit ΔuORF effect (*ded1-cs/DED1*) ratios significantly < 1.0 for the reporters with WT structures (light grey bars) but not those with mutated structures (adjacent black bars).

**Fig. S12.**
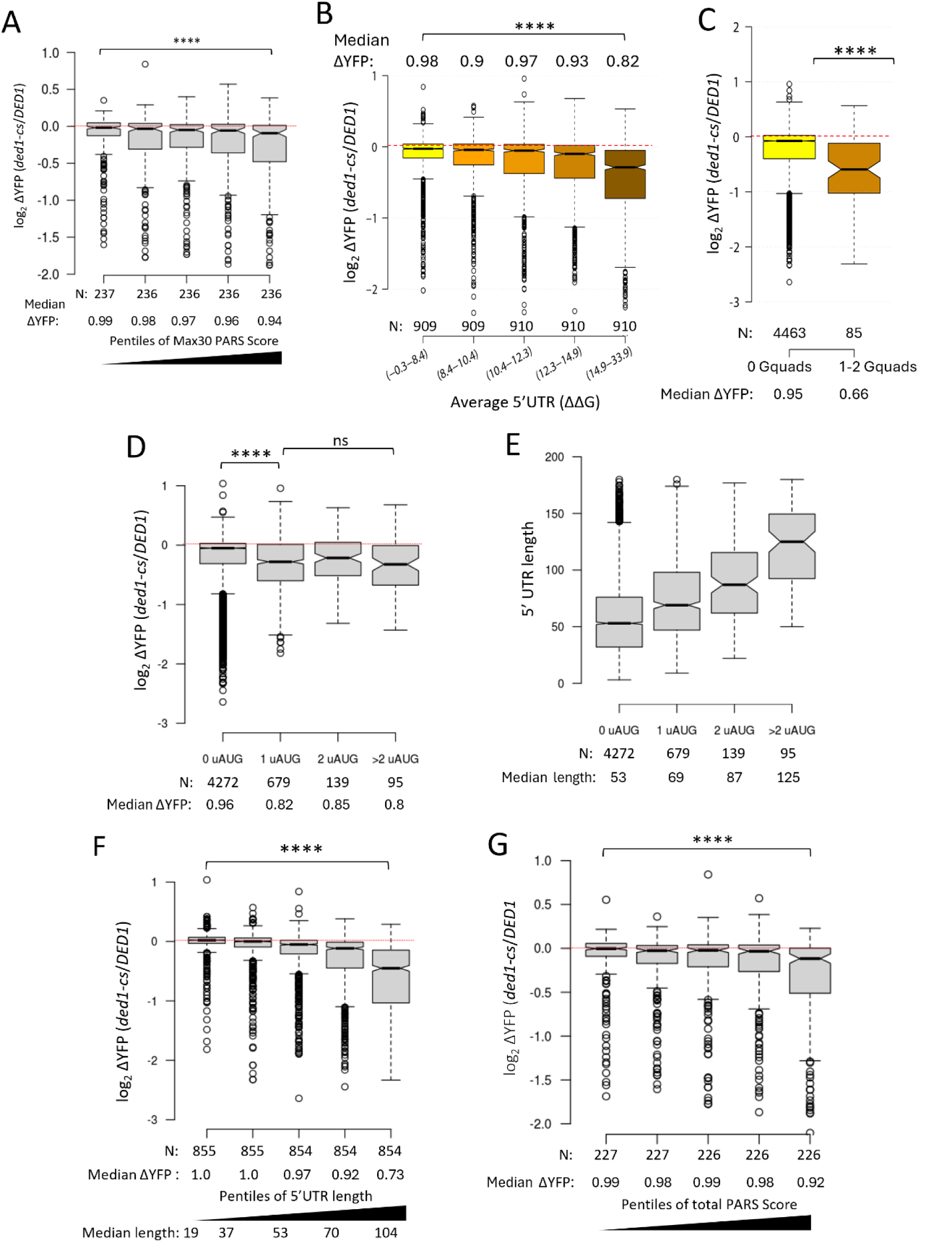
Evidence that 5’UTR length and secondary structure rather than the presence of upstream AUGs dictate Ded1-dependence of YFP reporter expression in MPRA of most WT reporters in the FACS-uORF library. **(A)** Similar analysis as in Fig. 7D but binned according to the 5’UTR Max30 PARS scores for the subset of 1258 WT YFP reporters with available PARS data. **(B-C)** Notched boxplot of the log_2_ changes in reporter expression between *ded1-cs* and *DED1* cells for the subsets of WT YFP reporters binned according to the estimated ΔΔG required to unfold the entire 5’UTR computed with RNAfold (A) or by the number of predicted g-quadruplexes in the 5’UTRs (B) for 4548 of the WT YFP reporters. **(D)** Notched boxplot of the log_2_ changes in reporter expression between *ded1-cs* and *DED1* cells for the subsets of WT YFP reporters binned according to the number of AUG codons in the 5’UTRs for which Mut_uORF reporters are represented in the FACS-uORF library. **(E)** Notched boxplot of 5’UTR length for the same bins of WT YFP reporters in (C). **(F-G)** Notched boxplots of the log_2_ changes in reporter expression between *ded1-cs* and *DED1* cells for the subset of 4272 WT YFP reporters lacking an AUG codon in the 5’UTR represented by a Mut_uORF reporter in the FACS-uORF library and binned according to reporter 5’UTR length (E), total 5’UTR PARS score (F), or Max30 PARS score. Results of Mann-Whitney U-tests for all panels are summarized as: ****, P<0.0001; n.s., not significant, P>0.05.

**Supplementary Table S1.**
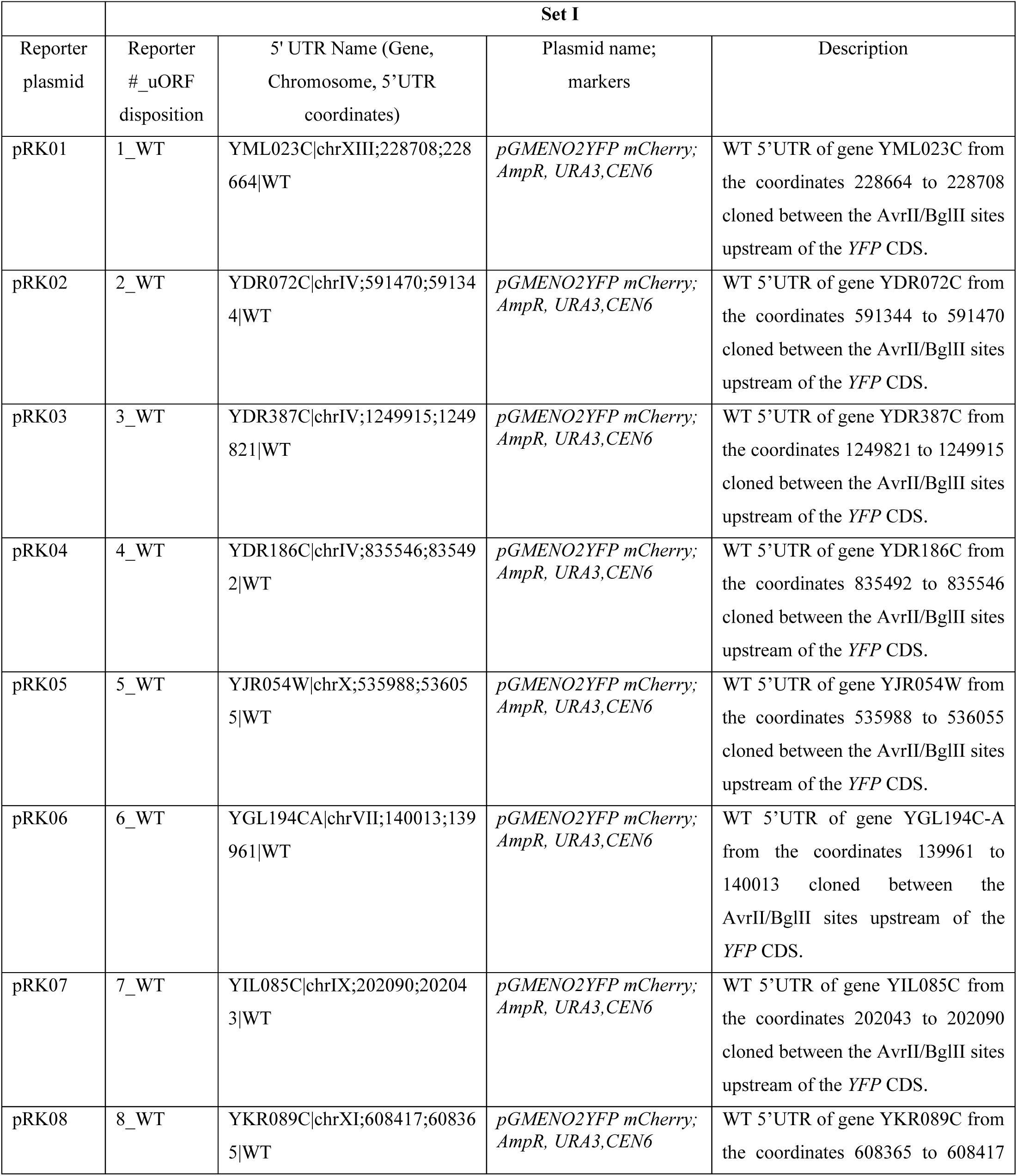
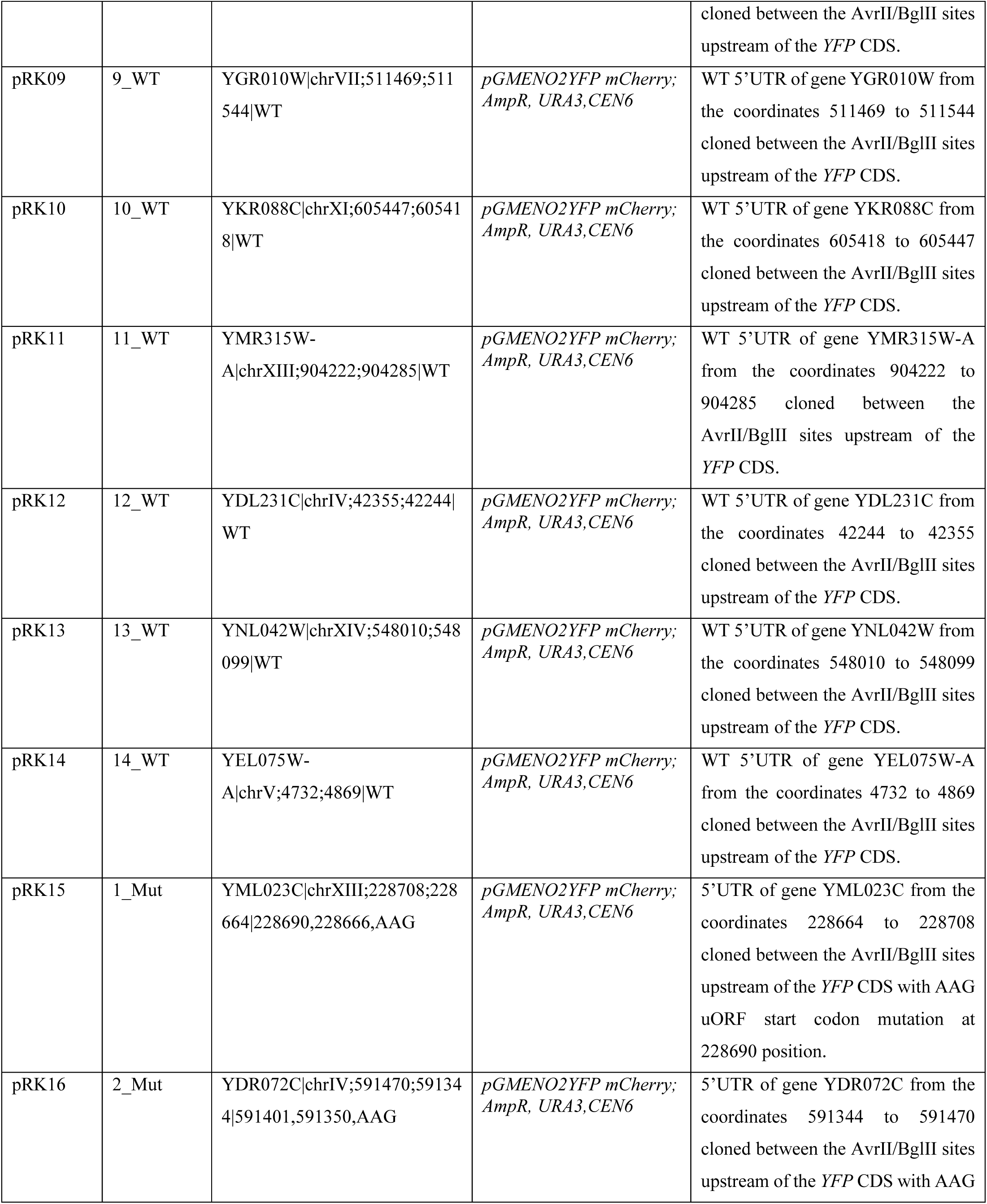
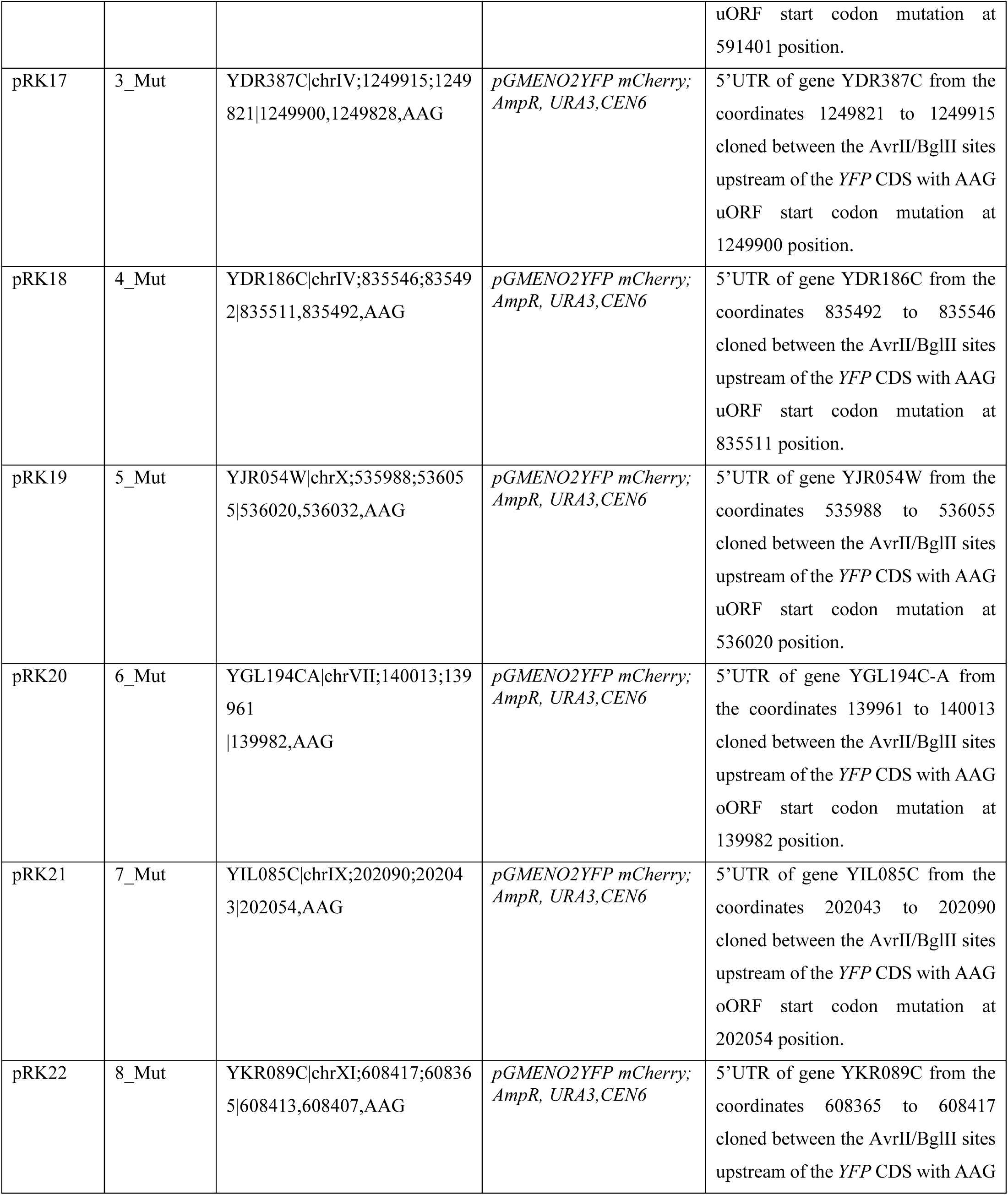
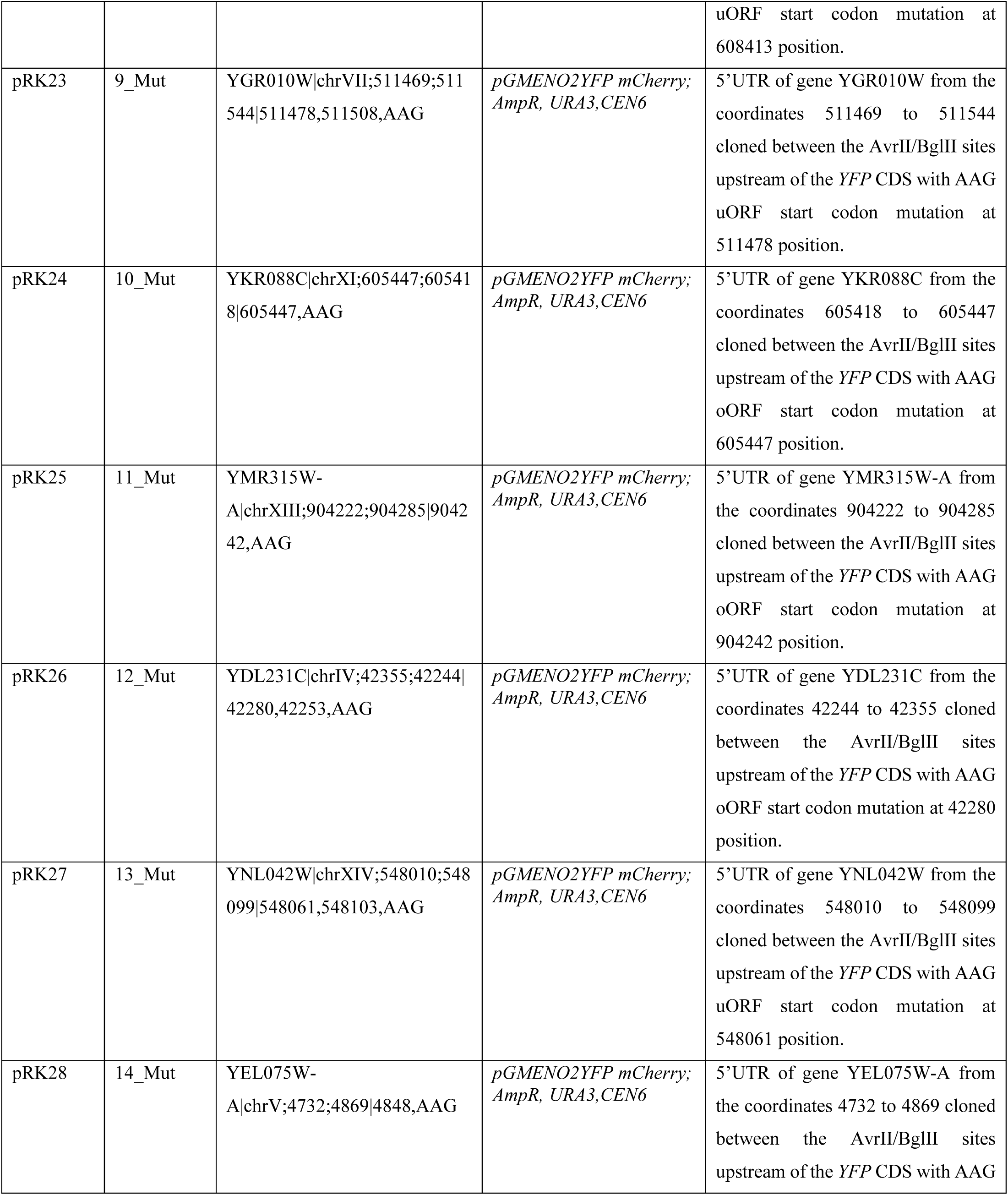
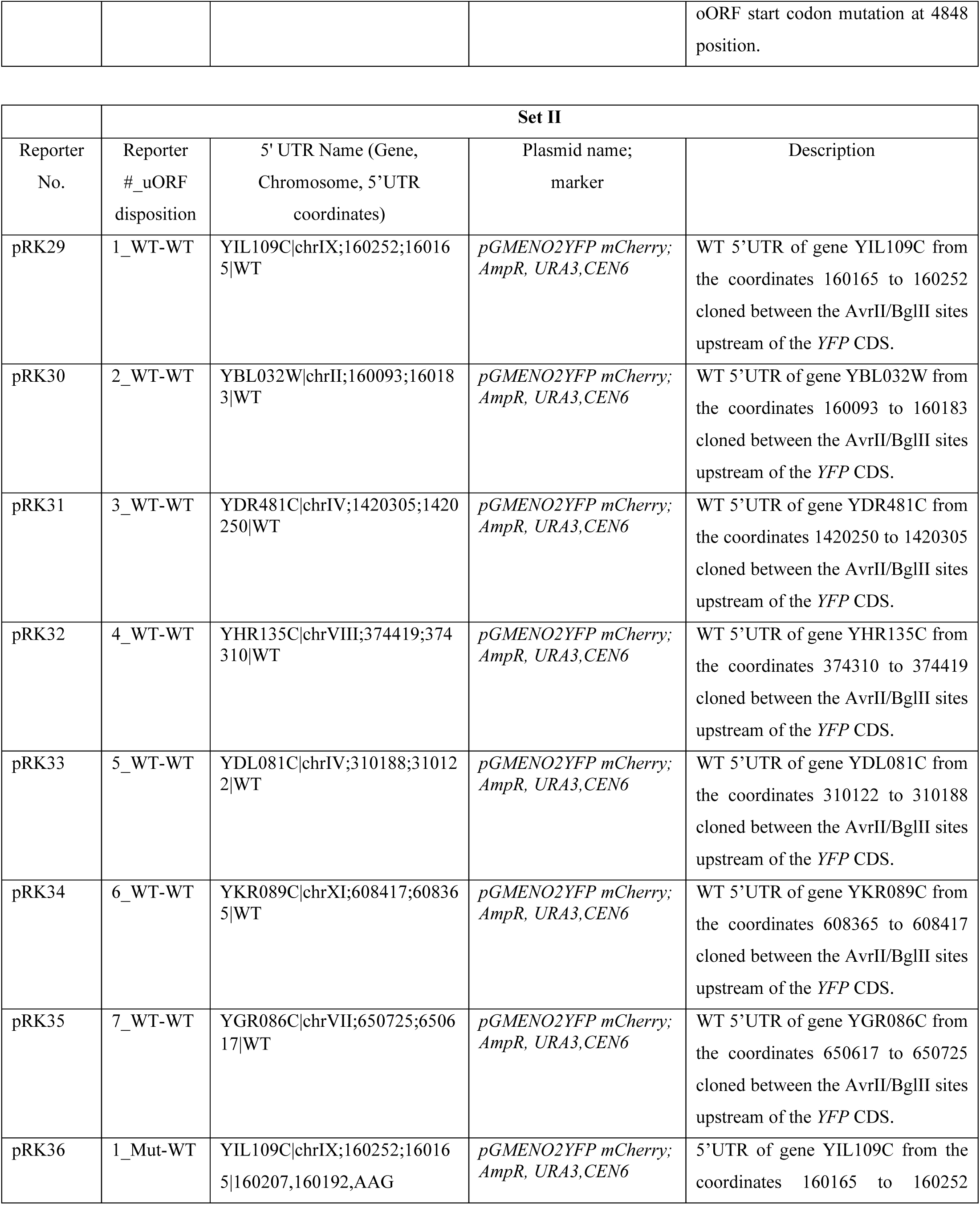
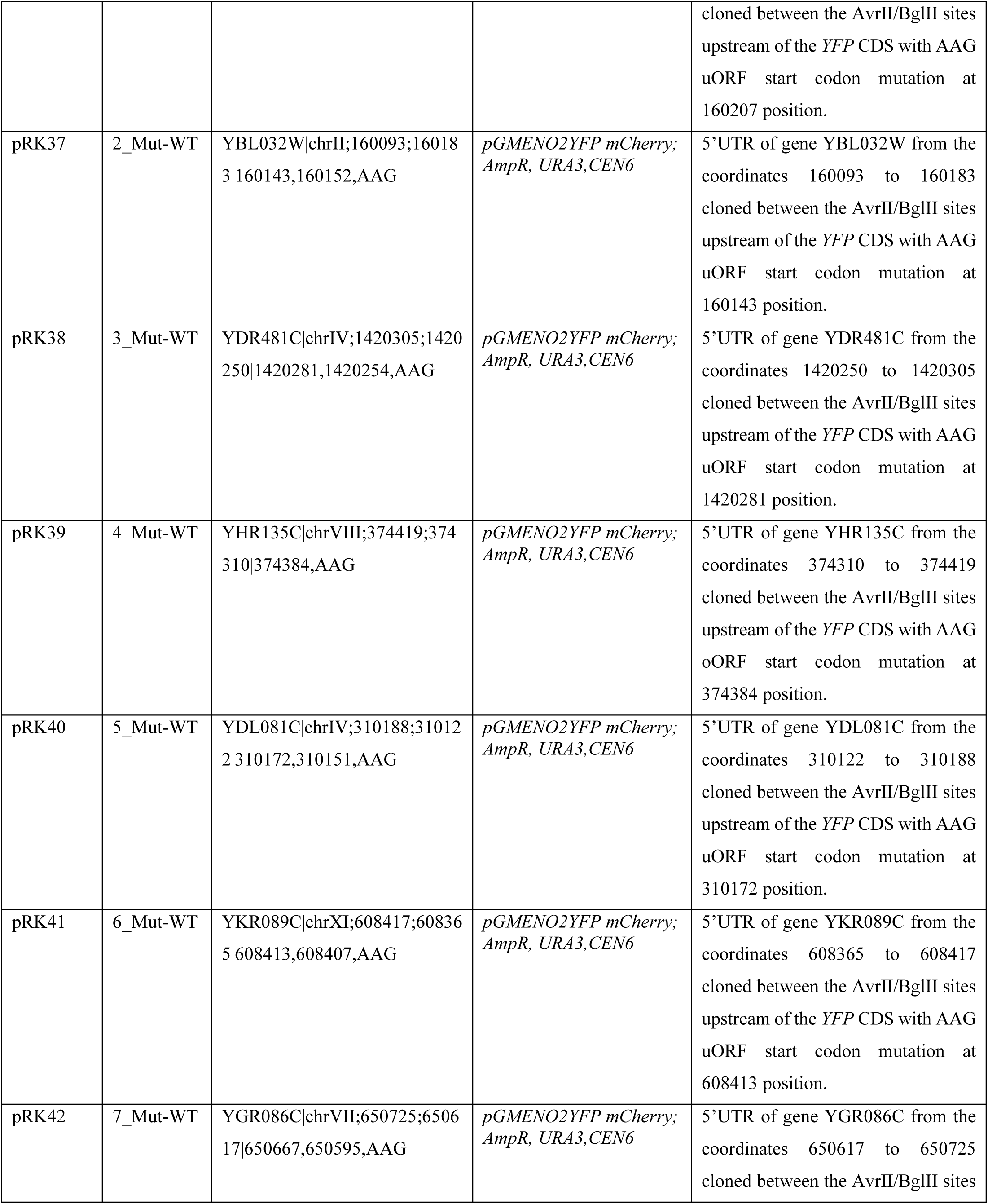
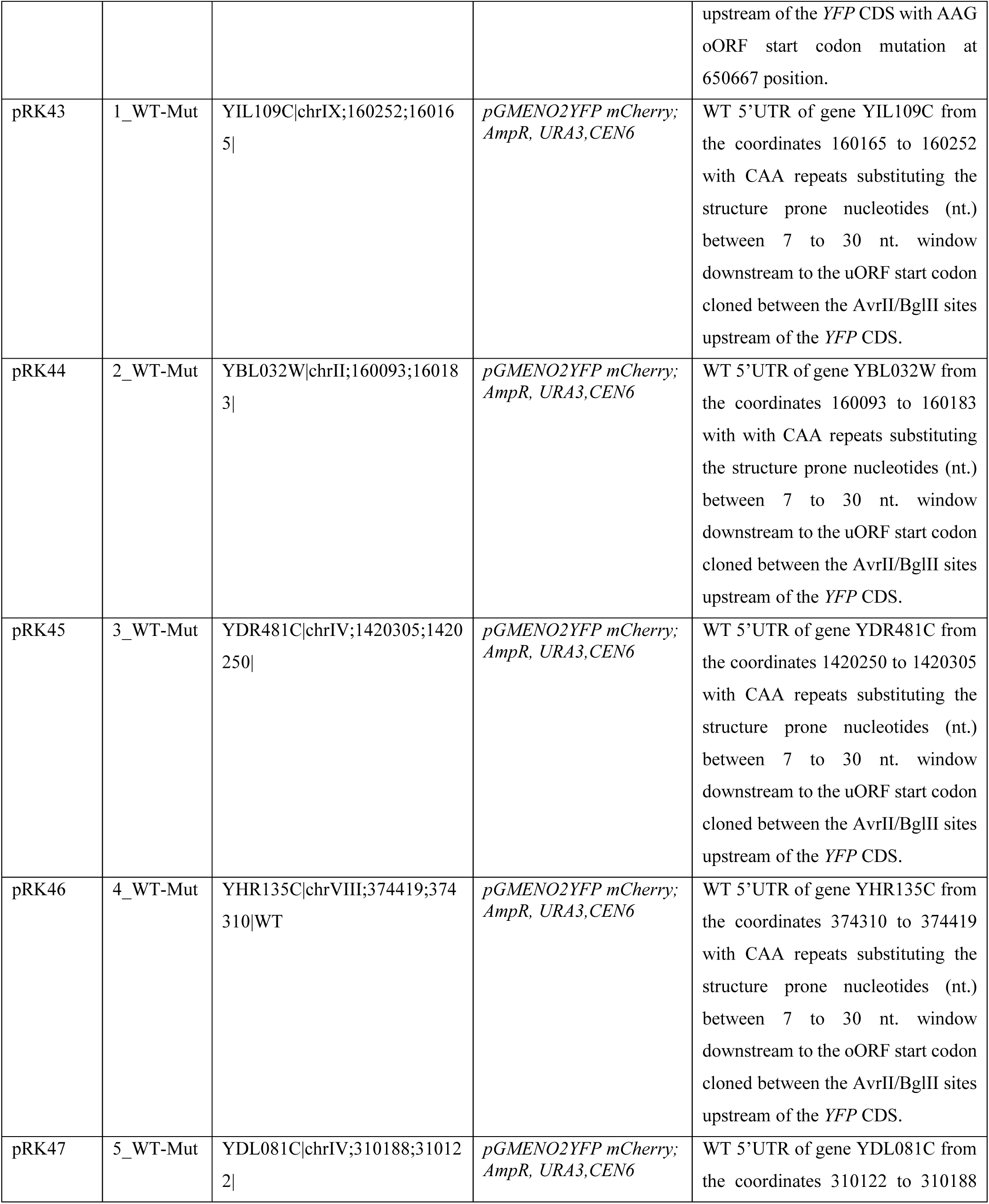
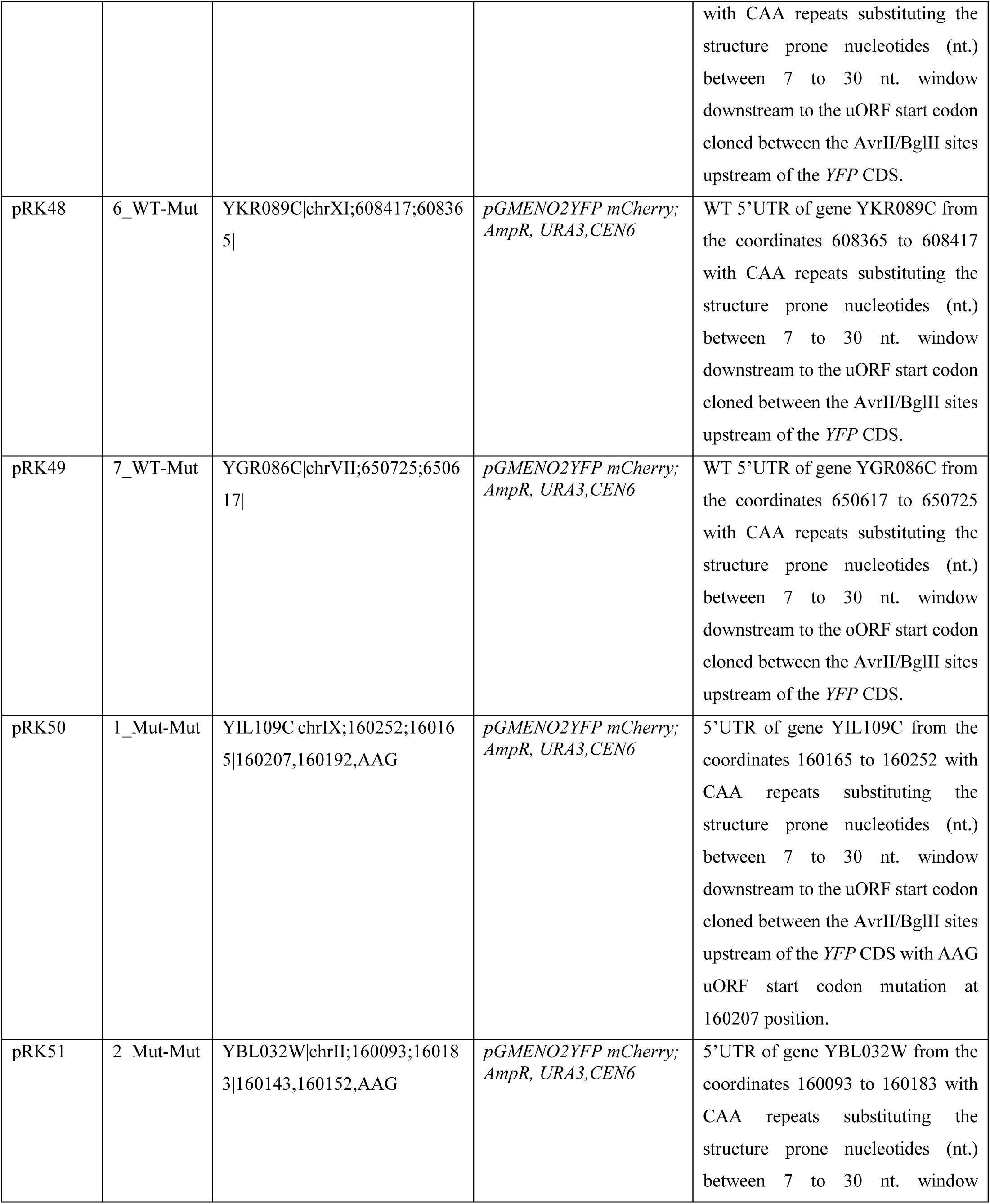
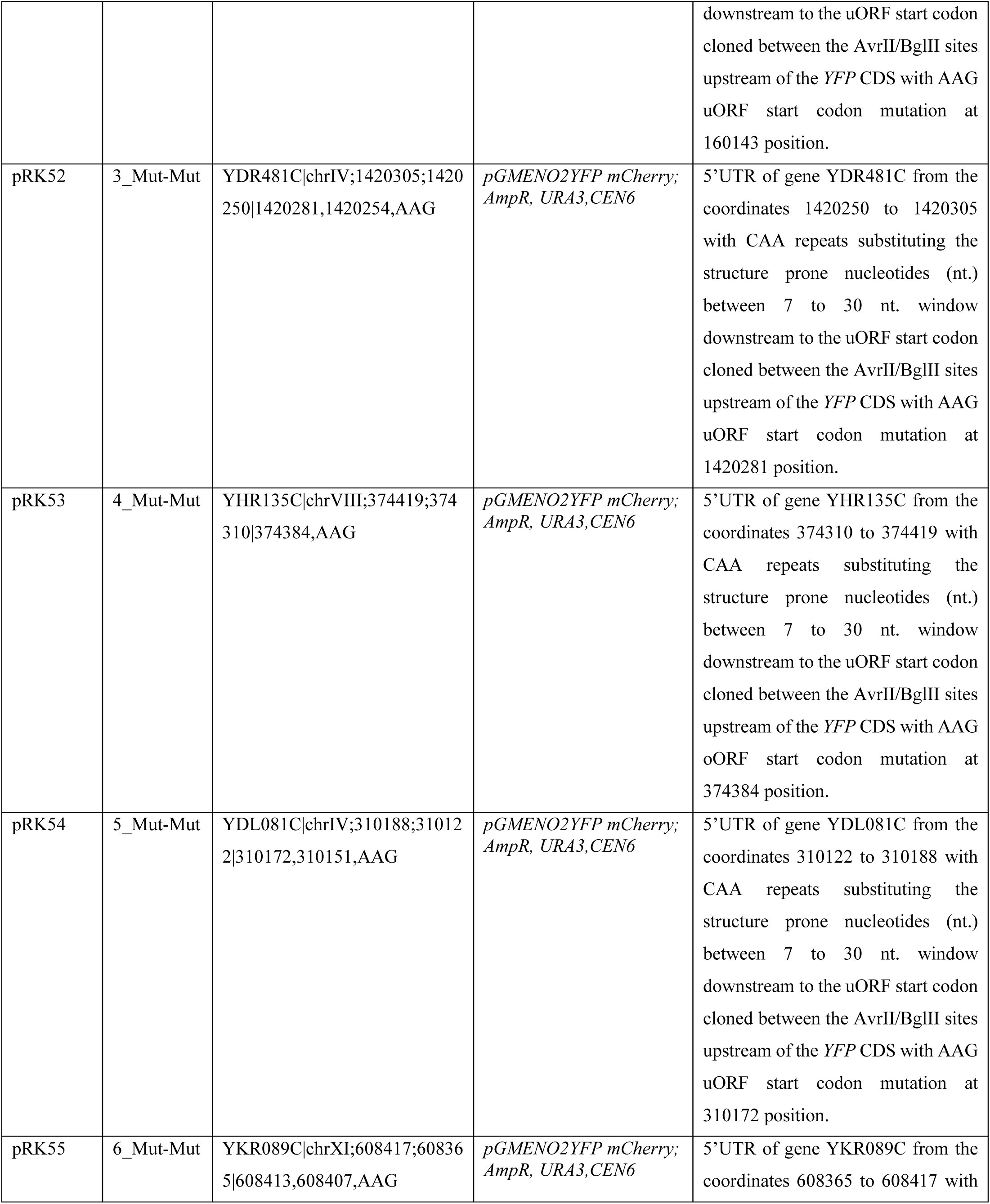
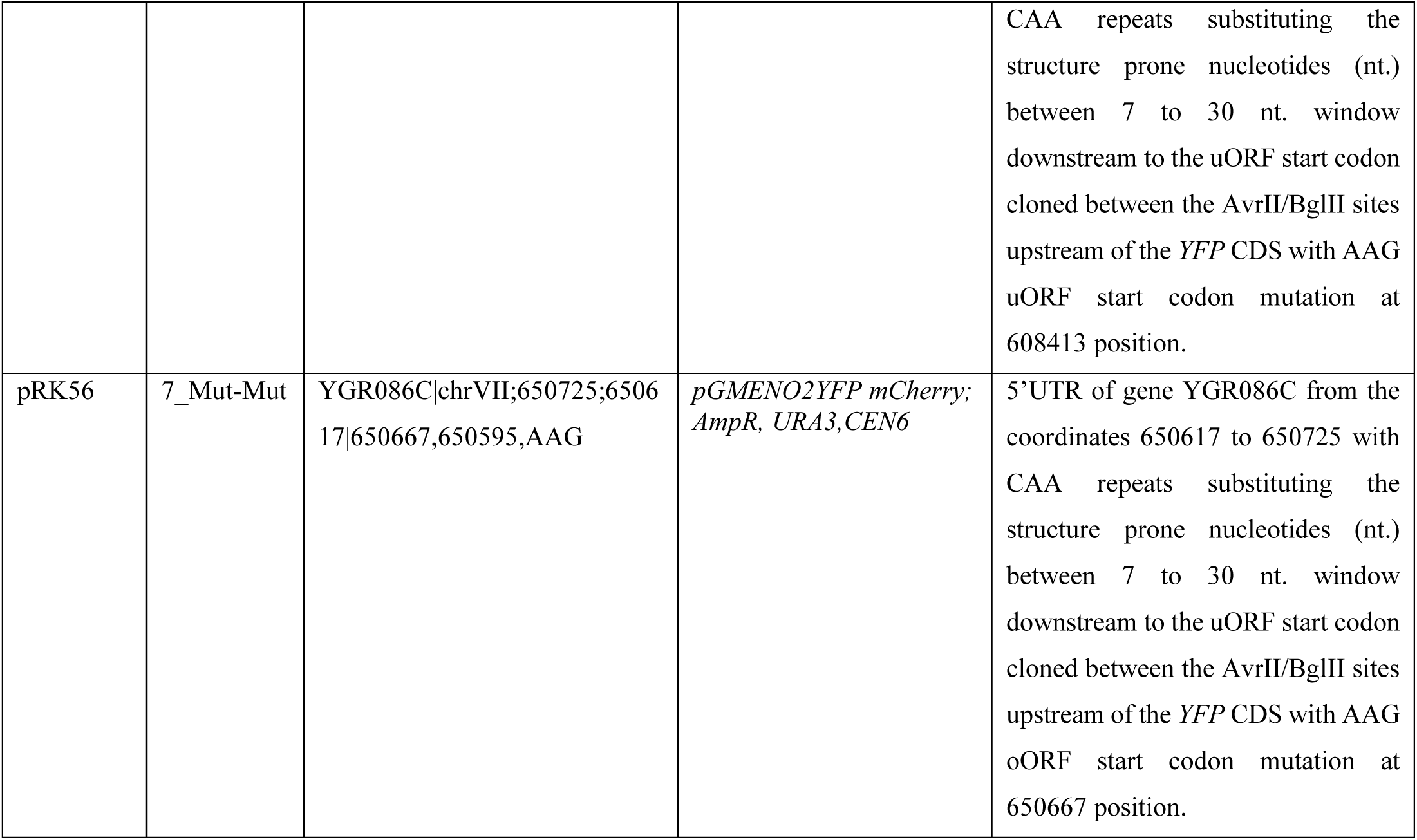
YFP reporter plasmid descriptions.

## SUPPLEMENTARY DATA FILES

**File S1. Source Data for Figs. 2A-2D**. This file lists the TE changes in mORFs and uORFs for all expressed mRNAs in *ded1-cs* versus *DED1* cells from (-CHX) Ribo-seq experiments, determined by DESeq2 analysis (sheets 2-3). The data and analyses used to generate Figs. 2A-D are provided in sheets 4-6.

**File S2. Source Data for Figs. 3A-E and Figs. S8B-G.** This file lists mRNAs containing unique AUG- initiated uORFs or NCC uORFs along with the data and analyses used to generate Figs. 3A-E (sheets 2-8) and Figs. S8B-G (sheets 9-14) respectively.

**File S3. Source Data for Fig. 4B**, **Figs. 7A-E, Figs. S10A-D, and Figs. S12D-G.** This file contains all primary FACS-uORF data for the *DED1* and *ded1-cs* strains and supporting analysis used to generate Figures 4B, 7A-E, S10A-D, and S12D-G.

**File S4. Source Data for Fig. 5**. This file contains all YFP reporter data and supporting analysis used to generate Figure 5.

**File S5. Source Data for Fig. 6 and Fig. S11B-D.** This file contains all YFP reporter data and supporting analysis used to generate Figures 6 and S11B-D.

**File S6. Source Data for Figs. S1A-D.** This file lists the TE changes in mORFs for all expressed mRNAs in response to the *ded1-cs* or *ded1-ts* mutations under +CHX and -CHX conditions. Expression values for all expressed mRNAs obtained from DESeq2 analysis of the corresponding ribosome profiling datasets are listed. Data for +CHX experiments was taken from Sen et al. (1) (GEO file GSE111255).

**File S7. Source Data for Figs. S2A-E and Figs. S3A-B.** This file lists number of RPF reads aligned to transcriptome in *ded1* mutants and respective *DED1* strains under -CHX and +CHX conditions (sheet 2), TE changes in uORFs for all expressed mRNAs in *ded1* versus respective *DED1* cells from (-CHX) Ribo-seq experiments, determined by DESeq2 analysis (sheets 3-4). The data and analyses used to generate Fig.S2E and Figs. S3A-B is given in sheet 5 and sheets 6-7 respectively.

**File S8. Source Data for Figs. S4A-D:** This file lists the changes in TE and center of ribosome density (CRD) of expressed mRNAs in response to the *ded1-cs* or *ded1-ts* mutations under +CHX or -CHX conditions.

**File S9. Source Data for Figs. S7A-C and Fig. S8A.** This file contains list of mRNAs containing unique AUG-initiated uORFs or NCC uORFs identified by May et al. (5), Spealman et al. (4), and Zhou et al. (3) (sheets 2-7) used to generate Fig. S7A-C and the data used to generate Fig. S8A (sheet 8).

**File S10. Source Data for Figs. S6A-B.** This file contains the data to generate Figs. S6A-B.

**File S11. Source Data for Fig. S11A.** This file contains the 5’UTR DMS MapSeq data from Guenther et al (2) used to generate Fig. S11A.

**File S12. Source Data for Figs. S12A-C.** This file contains the FACS-uORF used data to generate Figures S12A-C.

